# Scalable localization of clustering instability for spatial transcriptomics

**DOI:** 10.64898/2026.10.08.738042

**Authors:** Wenshan Wu, Joe Nguyen, Erin K. Molloy

## Abstract

Standard spatial transcriptomics workflows rely on clustering methods to define regions of interest for down-stream analyses, such as differential expression and pathway enrichment. Although researchers are presented with a single partition, most clustering methods are stochastic, meaning their output depends on random algorithmic choices, typically controlled by a seed; thus, repeating the analysis changing only the seed can yield a different result. Spatial transcriptomics data may be especially sensitive to algorithmic perturbations due to high dimensionality, sparsity, and mixed expression profiles at tissue boundaries and transitional zones; however, the extent of seed-dependent variation and its downstream impact have yet to be systematically evaluated. To bridge this gap, we present InSTability, a framework that localizes disagreement among partitions at the observation level. InSTability is built on (1) an interpretable, per-spot score that quantifies the variability of a spot’s cluster assignment across partitions, and (2) an algorithm with time complexity linear in the number of spots for a specified number of runs for computing the instability score at all spots. Together, these advances enable spatially-resolved clustering instability maps for large, high-resolution datasets. Applying InSTability to repeated runs of seven established methods (BayesSpace, GraphST, Leiden, Louvain, SEDR, STAGATE, and SpiceMix) on 18 benchmark samples reveals that clustering instability varies across tissue sections, with shared spatial patterns emerging among different methods applied to the same sample. Furthermore, clustering instability correlates with poor agreement against reference annotations and propagates downstream into differential expression analyses, resulting in inconsistent gene sets across seeds. Thus, InSTability can strengthen spatial transcriptomics studies by highlighting regions where clustering is seed-sensitive and further validation is warranted. InSTability is open-source and available at https://github.com/molloy-lab/InSTability.

---

Clustering algorithms are widely employed in spatial transcriptomics (ST), with high-performance implementations available in Python (e.g., Scanpy [1]), R (e.g., Seurat [2]), and 10x Genomics’ Space Ranger pipeline [3, 4]. The goal is to partition spots, or micron-sized bins, into coherent transcriptional domains that serve as the basis for downstream analyses, such as differential gene expression or pathway enrichment. Clusters also inform manual annotation, during which they are assigned biological interpretations. As a result, clustering algorithms play a central role in shaping the direction of investigation and the biological conclusions drawn from ST data.

Although standard ST workflows rely on a single reported clustering realization for downstream analyses [5–10], most methods are stochastic, meaning their output depends on random algorithmic choices, typically controlled by a seed. As an example, Louvain clustering [11], used by Space Ranger, initializes each spot as its own cluster and iteratively moves spots between clusters when doing so improves the modularity objective. Spots are processed in a random order, and this order, along with other randomized choices made during optimization, depends on the seed. Consequently, different seeds can yield different clustering results. We say that a cluster is *stable* when its constituent spots consistently cluster together across different seeds, or algorithmic perturbations, extending the notion of cluster stability under data perturbations [12, 13].

Spatial transcriptomics presents computational challenges arising from its spatial structure and scale [14]. These analyses may be especially sensitive to algorithmic perturbations due to high dimensionality, sparsity, and mixed expression profiles at tissue boundaries or transitional zones, such as tumor leading edges and tumor-normal interfaces, where molecular signatures derived from clusters guide clinical interpretation and decision making [15– 17]. Consequently, researchers analyzing the exact same dataset with the exact same methodology could arrive at different partitions simply by choosing different seeds. Whether these differences alter downstream analyses or biological conclusions is directly tied to the magnitude and spatial localization of clustering instability, which is not readily assessable with current software tools. To date, the prevalence and spatial distribution of seed-dependent variability in ST clustering, and its consequences for downstream analyses and biological conclusions, have not been systematically characterized.

## Previous work

Clustering gene expression data has been the subject of extensive research, and it is well appreciated that clusters derived from single-cell and spatial data can exhibit inherent ambiguity. Soft or fuzzy clustering methods address this issue by assigning membership weights to each spot or cell, indicating partial membership in multiple clusters [18–23]. When combined with probabilistic frameworks, including Bayesian, mixture, topic, or latent factor models, these weights can be interpreted as probabilities of cluster membership, enabling observation-level quantification of uncertainty [24–28]. Examples of such methods designed for ST data include BayesSpace [29], BASS [30], BayeSMART [31], ACSpot [32], and SpaTM [33]. Although the outputs of these methods can be visualized to reveal regions of high or low membership entropy, this does not indicate whether repeating the analysis with a different seed would yield the same result. Moreover, model-derived uncertainty estimates are restricted to specialized methods, whereas algorithmic stochasticity affects many methods, including graph-based community detection methods, Leiden and Louvain, which lack internal uncertainty metrics but dominate standard ST workflows due to their speed and simplicity.

Consensus frameworks complement soft clustering, although they are more commonly applied in single-cell rather than spatial transcriptomics. Ensemble methods, such as SC3 [34] and SAFE-clustering [35], combine results from multiple base methods or parameter settings into a unified consensus partition. Similarly, bootstrapping and resampling approaches identify stable clusters from random perturbations of the input data [13]. Significance-based hierarchical clustering methods, such as sc-SHC [36], apply statistical tests along a hierarchical clustering tree to guide cluster resolution; this strategy is used by Space Ranger to merge clusters produced by Louvain [4]. Thus, the goal of consensus-based frameworks is to produce a single, improved final partition—which is still dependent on the seed choice when the underlying base methods are stochastic—rather than to localize variation across the input partitions.

### Our contribution

We present *InSTability*, a general computational framework that localizes partition disagreement at the observation level. InSTability is based on two core advances. First, we formulate an interpretable, per-spot score that captures the variability of a spot’s cluster assignment across partitions. This approach enables spatial visualization while overcoming the inherent limitation of model-derived uncertainty estimates, which are tied to specific models or algorithms, and thus cannot be broadly deployed. Second, we give a linear-time algorithm for computing the instability score at all spots, bypassing the computational bottleneck of pairwise spot comparisons. Together, these advances enable spatially-resolved instability mapping for large, high-resolution datasets.

Application of InSTability to repeated runs of seven established clustering algorithms—spanning probabilistic, graph-based, and deep learning frameworks—on 18 benchmark samples leads to several key findings. First, seed-dependent clustering instability varies across tissue sections, with shared spatial patterns emerging among different methods applied to the same sample. Second, clustering instability correlates with poor agreement against reference annotations. Third, clustering instability propagates downstream: stable clusters yield consistent sets of differentially expressed genes across seeds, whereas gene sets from unstable clusters are highly variable. Together, these findings demonstrate the utility of InSTability in spatial transcriptomics workflows, from providing a fast, unsupervised check on overall clustering quality to pinpointing tissue regions where clustering is seed-sensitive and thus further validation or alternative analytical approaches going beyond clustering are warranted.

## Results

### InSTability is interpretable and scalable

Our framework, InSTability, takes as input a set of *R* partitions, for example those created through repeated runs of the same method on the same input, differing only by the random seed (Fig. 1). Intuitively, a spot’s cluster assignment is *stable* when the spots that cluster with it in one run tend to cluster with it in the other runs, while spots assigned to different clusters tend to remain separate; otherwise, the cluster assignment is *unstable*.

**Fig. 1.**
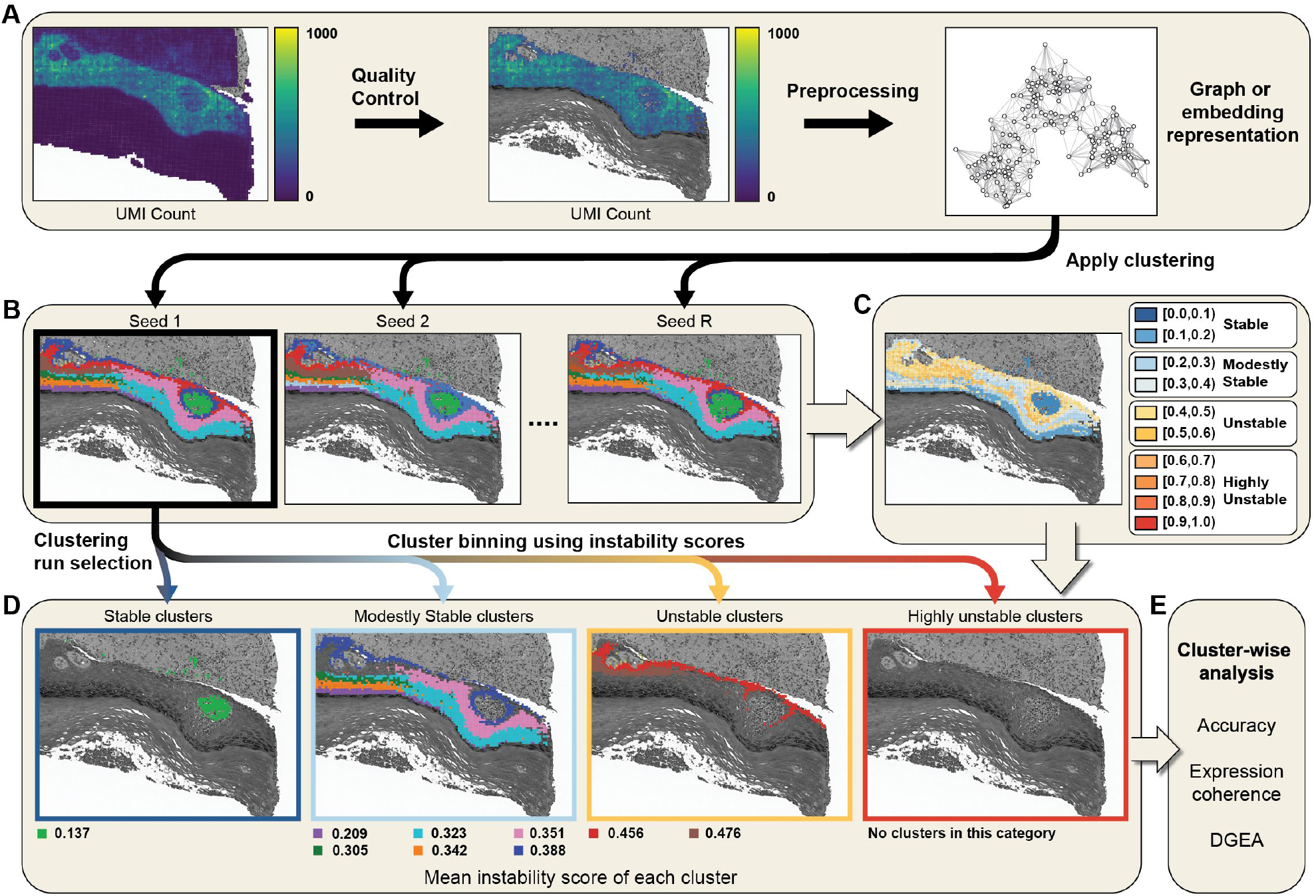
Cluster InSTability Framework. **A)** Standard spatial transcriptomic workflows include data quality control and preprocessing steps, ultimately producing a graph or reduced embedding representation that can be used for clustering. **B)** InSTability is applied to partitions generated from repeated clustering runs on the same input, changing only the seed. **C)** The resulting instability map reveals how (in)consistently each spot co-clusters with the other spots across the input partitions. **D)** For a particular partition of interest, which may be selected on the basis of biological plausibility, domain knowledge or other study-specific criteria, the stability of each cluster can be categorized based on the mean score of its spots. **E)** Instability scores can inform cluster ratings for downstream analyses.

We quantify this behavior using pairwise cluster co-membership, called co-association, across runs. Let 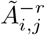 denote the fraction of runs *excluding r* in which spots *i* and *j* are assigned to the same cluster. Let 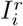 be the set of spots assigned to the same cluster as spot *i* in run *r*, excluding spot *i*, and let 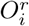 be the set of spots assigned to different clusters than spot *i* in run *r*. We then define the *instability score* of spot *i* as

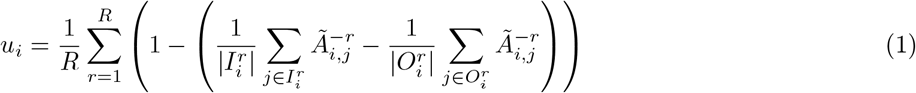

where the first term in the innermost brackets represents the *intra*-cluster co-association and the second term represents the *inter*-cluster co-association, both for spot *i* with respect to run *r*.

A key advantage of our instability score is that its value range is well-behaved. The minimum score of 0 occurs when the cluster containing the spot is identical across the input partitions, as intra-cluster co-association is 1 and inter-cluster co-association is 0. When cluster assignments are random, the expected intra- and inter-cluster co-associations are equal, resulting in an expected instability score of 1. Although the score can be greater than 1 when the partitions differ by more than what is expected by chance, this does not occur in our experiments, suggesting an empirical maximum of 1. We categorize scores between 0.0 and 0.2 as *stable* because this corresponds to high intra-cluster co-association (*>*0.8) and low inter-cluster co-association (*<*0.2) on average across runs. We further categorize scores as *modestly stable* (between 0.2 and 0.4), *unstable* (between 0.4 and 0.6), and *highly unstable* (between 0.6 and 1.0). Individual clusters and entire samples are assigned to these categories based on the mean instability score of their spots.

Instability score can be naively computed from the pairwise co-association matrix, requiring quadratic storage in the number *n* of spots, i.e., Θ(*n*^2^). The largest Visium HD sample used in our study has 503,622 spots, so storing strict upper right triangle of the matrix would require more than 500 GB of storage at 4-bytes (i.e., float32) per entry. To address this bottleneck, we give an algorithm for computing the complete spatially-resolved instability map in Θ(*R*^2^*n*) time and Θ(*Rn*) space, where *R* is the number of runs and *n* is the number of spots. This formulation is critical because *R* can be treated as a fixed constant: it is capped at 50 in our experiments, making *R* approximately four orders of magnitude smaller than the size of the largest Visium HD sample.

### InSTability can be leveraged to localize seed-dependent clustering instability

To evaluate the extent and potential downstream impact of clustering instability, we applied InSTability to 18 publicly available ST benchmark samples with reference annotations. Samples varied in tissue type, disease condition, and sequencing technology. Henceforth, we refer to these samples using the following abbreviations: DLPFC denotes a collection of 12 human dorsolateral prefrontal cortex Visium samples [37]; HBC denotes a human breast cancer Visium sample [38]; MB denotes a mouse brain Visium sample [39]; MBC denotes a mouse brain cerebellum Slide-seq sample [40]; CRC denotes a human colorectal cancer Visium HD sample [41]; COAD denotes a human colon adenocarcinoma Visium HD sample [42]; and OV denotes a human ovarian cancer Visium HD sample [42].

Clusters were computed with seven methods spanning classical non-spatial graph community detection (Leiden [43] and Louvain [11]), Bayesian spatial modeling (BayesSpace [29]), deep graph-based spatial representation learning (STAGATE [44], SEDR [45], and GraphST [46]), and probabilistic spatial mixture modeling (SpiceMix [47]). This selection was designed to encompass both traditional and modern approaches for ST clustering while avoiding methods that rely heavily on histological images, although evaluating the stability of such approaches is an important direction for future work. Prior to clustering, we applied standard quality-control and preprocessing procedures.

To summarize, the primary benchmark comprised 125 method–sample configurations after omitting one failure: STAGATE failed to run on the MBC sample due to a memory error. To evaluate the impact of backend clustering algorithms, we re-ran SEDR substituting the Leiden backend with the mclust backend (i.e., Gaussian mixture modeling) on the 12 DLPFC samples only; this is important because many ST clustering methods, including STAGATE and GraphST, apply traditional clustering algorithms to learned representations (all methods were run with the Leiden backend unless otherwise noted). To assess computational scalability, we ran the two fastest clustering methods, Leiden and Louvain, on three full-resolution Visium HD datasets. Clustering was repeated *R* = 50 times for each method–sample configuration, changing only the seed, for a total 7,150 clustering runs.

### Clustering instability varies across tissue sections, with shared spatial patterns emerging among methods applied to the same sample

#### Trends for DLPFC sample 151674

Taking DLPFC sample 151674 as an example (Fig. 2), clustering instability varied across spatial coordinates, with at least ~10% of spots having stable cluster assignments and more than ~20% of spots having highly unstable cluster assignments (BayesSpace was an exception exhibiting greater stability than the other methods). Rather than being confined to isolated spots, instability was often spatially localized, affecting entire clusters in the example run. As a consequence, every method produced at least one stable cluster (mean = 1.5) and at least one unstable cluster (mean = 2.4). Spatial patterns of instability were consistent across many of the methods tested. The pink cluster corresponding to white matter was highly stable (mean score: *<* 0.1) for all methods, except BayesSpace for which it was still stably recovered but with mean score in the interval range [0.1, 0.2). In contrast, the right of the pink cluster was highly unstable for GraphST, STAGATE, SpiceMix, and SEDR. Leiden and Louvain also exhibited high instability in this region but produced fragmented patterns, reflecting scattered spot reassignments. Although the fragmented patterns from Leiden and Louvain may indicate poor clustering quality, the other methods produced clusters that were more spatially coherent, yet had high instability and discordance from the reference annotation. SEDR-mclust was distinctive from the other methods, with clustering instability largely confined to cluster boundaries; it also was the most consistent with the reference annotations, based on Adjusted Rand Index (ARI). Similar trends were observed across the other benchmark samples (Supplementary Figs. S1–S17). Although the degree of instability varied across samples (Fig. 3A), at least one method produced an unstable cluster for all 18 benchmark samples.

**Fig. 2.**
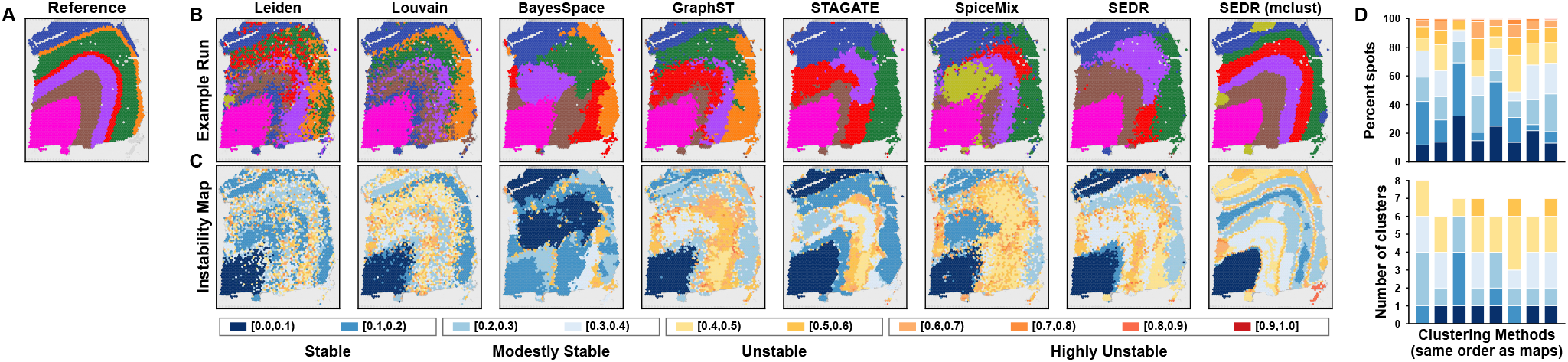
Clustering instability maps for DLPFC sample 151674. **A)** Reference annotation distributed with the benchmark. **B)** Example clustering run is shown for all methods tested. Adjusted Rand Index (based on reference annotations) going from left to right is 0.33, 0.36, 0.34, 0.43, 0.48, 0.37, 0.46, and 0.55. **C)** Spatial instability maps estimated from 50 stochastic runs for each method. Mean instability going from left to right is 0.28, 0.32, 0.18, 0.34, 0.23, 0.35, 0.31, and 0.32. Clustering instability was localized across spatial regions, with many methods exhibiting similar patterns. The hot pink cluster is stably recovered (mean instability score *<* 0.1) for all methods except BayesSpace. To the right of this region, many methods (e.g., STAGATE, SpiceMix, and SEDR) exhibited substantial instability. **D)** Stacked bars (one per method in the same order as the instability maps) show the percentage of spots assigned to each instability interval (top) and the number of clusters assigned to each instability interval based on the mean score across spots (bottom).

**Fig. 3.**
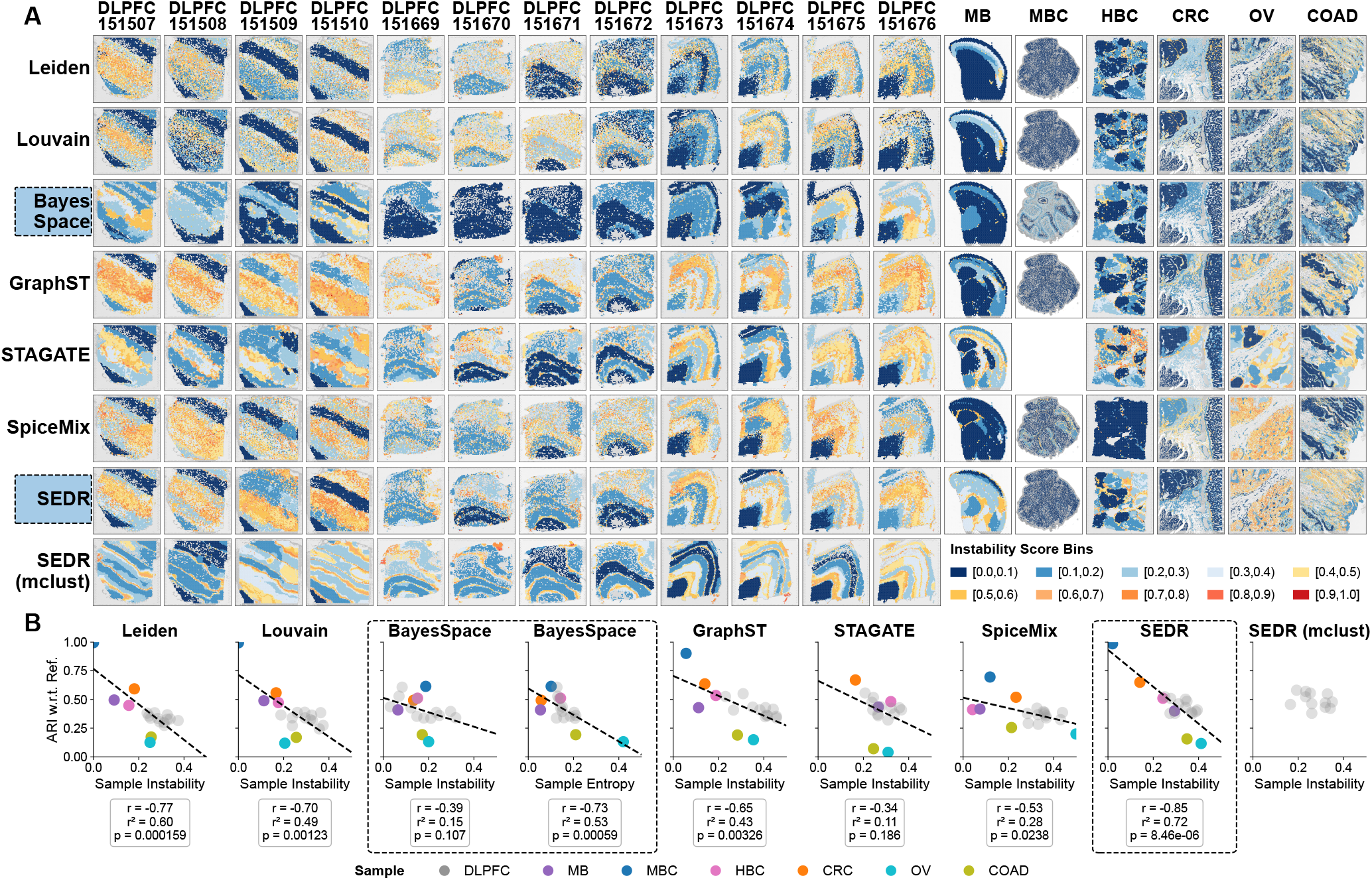
Instability across methods and samples. **A)** Spatial instability maps for 8 clustering configurations (rows) and 18 samples (columns). Spatial patterns and magnitudes of clustering instability share similarities across methods (rows) and vary across samples (columns). Missing panels are due to STAGATE encountering a memory error on the MBC sample and SEDR with mclust backend being run on the DLPFC samples only. **B)** Pearson correlation of clustering performance (mean ARI across 50 stochastic runs) versus instability (mean instability score across spots) for each clustering method. For BayesSpace, the dashed-box portion contains two comparisons: mean ARI versus mean instability and mean ARI versus cluster membership entropy, where entropy is averaged across all spots within each run and then across all 50 runs. Correlation analysis was not performed for SEDR with mclust backend, as it was only applied to the 12 DLPFC samples.

#### Selection of an example clustering run

As previously noted, instability of individual clusters was evaluated with respect to an example run; we selected the example, for each method–sample configuration, to correspond to the first run out of 50 achieving the lowest number of clusters. The number of clusters, required as input for BayesSpace and SEDR-mclust, was set to match the reference annotation, so the example run was the first run. Otherwise, the range in the number of inferred clusters across runs had a mean of 2.01 (s.d. = 1.38). As examples, Leiden produced 8–11 clusters on DLPFC sample 151674 (mean = 9.16, s.d. = 0.68) and 21–25 clusters on the HBC sample (mean = 22.42, s.d. = 1.01). Thus, the number of clusters produced by Leiden was slightly higher than the number of reference labels (7 for DLPFC 151674 and 20 for HBC), as expected based on our protocol for tuning the clustering resolution parameter.

#### Clustering instability is highly variable across samples

Sample-level instability was evaluated based on the percentage of spots with unstable cluster assignments, averaged across all methods applied to the same sample. Samples exhibited substantial variation in clustering stability (Supplementary Table S2). Only 3% (s.d. = 4%) of spots had unstable cluster assignments in MBC, the most stable sample, followed by MB, CRC, and HBC, for which 9–13% (s.d. = 7–9%) of spots had unstable assignments. The percentage of unstable spots increased to 25% (s.d. = 10%) for COAD and 35% (s.d. = 20%) for OV. The 12 DLPFC samples also exhibited substantial variation in instability, with the percentage ranging from 20% to 40%. DLPFC sample 151674 (Fig. 2) was near the middle of this range, with 30% of spots having unstable assignments.

Variation in clustering stability across samples was also apparent at the cluster level, evaluated as the percentage of stable clusters among the example runs of all methods applied to the same sample. At least 69% of clusters were stable for MBC, CRC, and MB, compared with 55% for HBC, 34% for COAD, and 36% for OV. Among the 12 DLPFC samples, the percentage of stable clusters ranged from 15% to 45% (Supplementary Table S2).

#### Clustering methods exhibit similar levels of instability with some notable exceptions

Clustering method instability was evaluated based on (1) the percentage of spots with unstable cluster assignments, averaged across all samples, and (2) the percentage of stable clusters among those inferred across all example runs (one per sample). Most methods exhibited similar levels of instability (Supplementary Table S1). The percentage of spots with unstable cluster assignments ranged from 24–27% (s.d. = 11–13%) for Leiden, Louvain, and STAGATE to 31–35% (s.d. = 16–20%) for SEDR, SpiceMix, and GraphST; thus, differences among methods were modest relative to variability across samples. BayesSpace was a notable exception, with just 9% of spots (s.d. = 9%) having unstable cluster assignments. Moreover, 69% of its inferred clusters were stable across stochastic runs, compared with 32–33% for SEDR, GraphST, STAGATE, and SpiceMix and ~45% for the community detection methods Leiden and Louvain.

These trends were apparent from spatial instability maps (Fig. 3A). BayesSpace was atypically stable for DLPFC samples 151509 and 151670–151672, and SpiceMix was atypically stable for the HBC sample and, to a lesser extent, the MB sample. Critically, being the most stable clustering method did not necessarily translate to better performance with respect to reference annotations. Although SpiceMix was among the most stable methods for the MB sample and was the most stable on the HBC sample, it had worse agreement with the reference annotations than GraphST, based on ARI averaged across all clustering runs. The mean ARI across samples ranged from approximately 0.38 for the worst methods to 0.45 (s.d. = 0.18) for the best method SEDR. BayesSpace fell in the middle with mean ARI 0.41 (s.d. = 0.13), despite being the most stable method overall (Supplementary Table S3).

#### Backend clustering algorithms impact stability of representation learning methods

In our primary benchmark, Leiden was used as the backend clustering algorithm for representation-learning methods. To evaluate the impact of the backend clustering algorithm on stability, we re-ran SEDR on the 12 DLPFC samples, replacing the Leiden backend with mclust [48]. Because mclust was provided with the target number of clusters set to match the number of reference domains, the total number of inferred clusters across the 12 example runs increased from 68 with Leiden to 76 with mclust (note that since the example run was selected as the first run with the minimum number of inferred clusters, the Leiden-backend example run could therefore contain fewer clusters than the target, even though resolution tuning generally favored cluster counts near or slightly above the target). Despite an increase in the number of clusters when switching from Leiden to mclust, the number of unstable clusters decreased from 21 to 18, corresponding to a reduction from 30.9% to 23.7% (Supplementary Table S1). Moreover, instability became more localized to cluster boundaries rather than spanning multiple clusters (e.g., Fig. 3A; Supplementary Figs. S1–S11), and mean ARI across the 12 DLPFC samples increased from 0.44 with the Leiden backend to 0.49 with mclust (Supplementary Table S3).

### Clustering instability is associated with poor agreement against reference annotations

Next, we evaluated whether clustering instability was associated with clustering performance. For each sample– method configuration, performance was quantified as the ARI between the inferred partition and the reference annotation, averaged across all 50 runs. Pairing the mean ARI for each sample with the mean per-spot instability score (i.e., averaged across all spots) revealed a negative association between clustering instability and performance, although the strength of this relationship depended on the method (Fig. 3B; Supplementary Tables S3–S4). The association was strongest for SEDR (Pearson *r*^2^ = 0.72), followed by Leiden (*r*^2^ = 0.60), Louvain (*r*^2^ = 0.49), and GraphST (*r*^2^ = 0.43). In contrast, the association was weakest for SpiceMix (*r*^2^ = 0.28), BayesSpace (*r*^2^ = 0.15), and STAGATE (*r*^2^ = 0.11). STAGATE failed to run on one sample (MBC), which may influence the correlation strength. The weaker association for SpiceMix may reflect its atypically stable behavior on the HBC and MB samples (Fig. 3A; Supplementary Figs. S12–S13). BayesSpace was an outlier in terms of stability, as previously discussed; it was also the only method that returned soft cluster assignments. Cluster membership entropy yielded a distinct pattern from instability, with high uncertainty near cluster boundaries and low values at cluster interiors (Supplementary Fig. S18), and was more strongly associated with poor performance (*r*^2^ = 0.53 versus 0.15) (Fig. 3B; Supplementary Tables S3–S5).

To investigate the connection between sample characteristics, specifically sequencing platform, we examined the results from SEDR, with the strongest association between clustering stability and performance. Both were high for the MBC and CRC samples: mean ARI of 0.99 and 0.65 versus mean instability of 0.02 and 0.14, respectively (these samples were generated with Slide-seq and Visium HD, respectively). Consistent with this, only 3% and 10% of spots had unstable assignments for MBC and CRC, respectively, averaged across methods (Supplementary Table S2). The HBC and MB samples, both generated with Visium, exhibited lower stability (mean scores = 0.24 and 0.29) and lower performance (mean ARI = 0.51 and 0.40). Similarly, the 12 DLPFC samples, also generated with Visium, exhibited lower stability and performance, although results varied across samples: the three samples on the high end (mean ARI = 0.50–0.52) had mean instability scores ranging from 0.25 to 0.34, whereas the three samples at the low end (mean ARI *≈*0.38) had mean instability scores ranging from 0.27 to 0.37. The worst samples COAD and OV exhibited low performance (mean ARI = 0.16 and 0.12) and low stability (mean scores = 0.35 and 0.41), despite also being generated with Visium HD. This may partly reflect their higher spatial resolution compared with the second best sample CRC (micron bins of size 8 *×*8 versus 16 *×*16), as smaller bins may contain fewer transcripts per observation and therefore provide less signal for clustering; it could also reflect the pronounced size imbalance among reference labels for OV and COAD compared with CRC.

### Cluster instability reflects poor cross-run cluster alignment and propagates to differential expression

Next, we evaluated whether cluster instability propagated to differential gene expression analyses (DGEA). To perform this analysis, we first aligned each cluster in the example run to its “best match” in each of the remaining runs based on maximum Jaccard similarity (Fig. 4A illustrates this matching procedure for SEDR applied to the HBC sample). Aligning clusters across runs allowed us to examine whether our instability scores agreed with traditional definitions of cluster stability [12]. We found this to be the case: stable clusters persisted across runs and unstable clusters did not (Fig. 4B; Supplementary Fig. S19). For SEDR, clusters with mean instability in range [0.0, 0.1) had high cross-run consistency: median best-match Jaccard of ~1. In contrast, clusters with mean instability in range [0.6, 0.7) had poor cross-run consistency: median best-match Jaccard of ~0.5, indicating that the intersection contained only about half as many spots as the union.

**Fig. 4.**
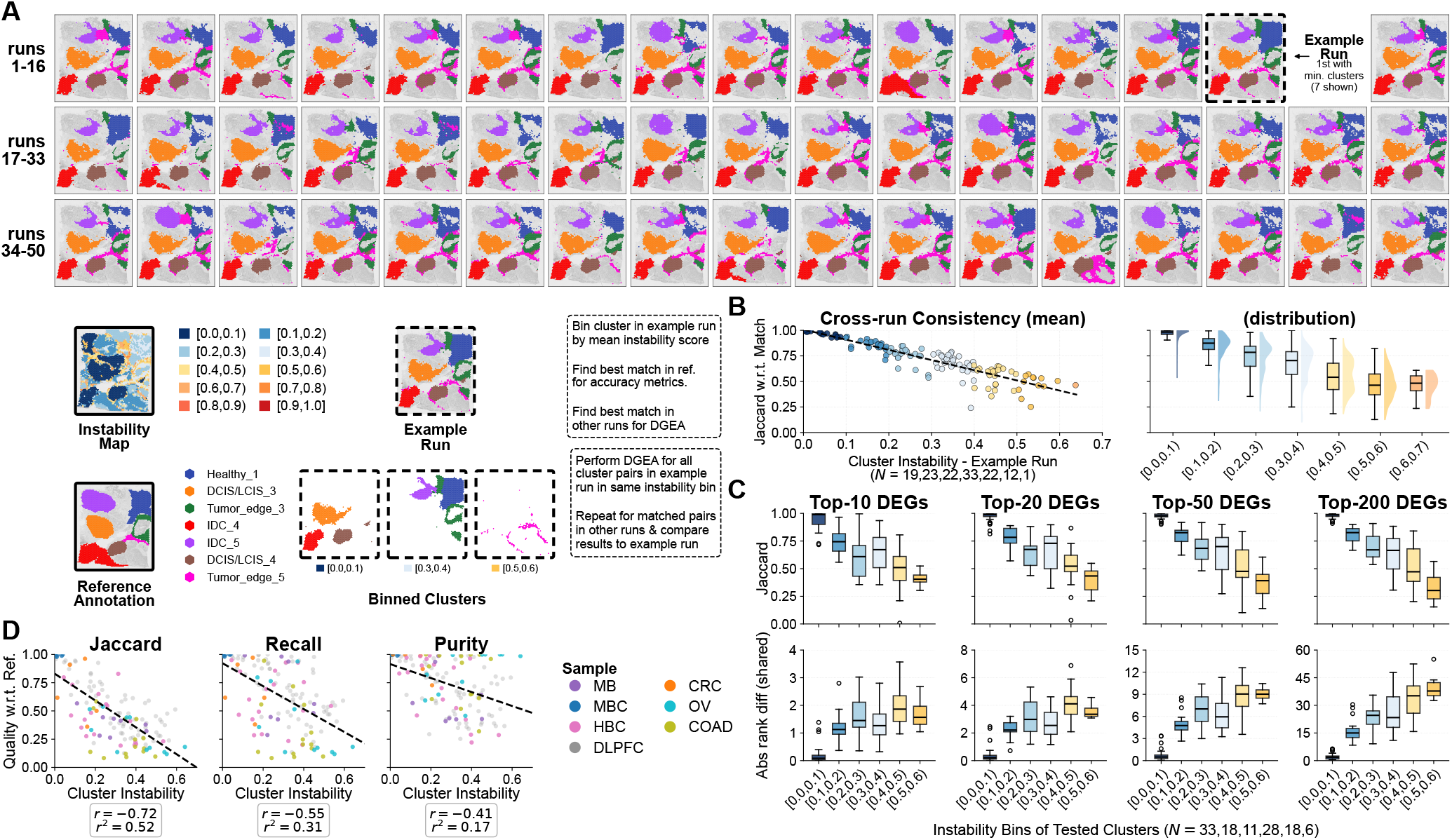
Cluster instability is negatively associated with cross-run reproducibility and quality based on reference annotations. All data shown are for SEDR. **A)** Analysis of SEDR on the HBC sample (only 6 clusters shown). Clusters are colored by best match to the example run, determined by maximum Jaccard similarity. **B)** Cluster cross-run consistency versus cluster instability. Left: Each dot is a cluster in the example run for a sample. *N* is the number of clusters in each instability bin (indicated by dot color). The *y*-axis is the Jaccard similarity between the spots in an example-run cluster and its best-matching cluster in another run, averaged across all non-example runs. Dashed line is the linear best fit. Right: Same underlying cross-run best-match Jaccard values as the left panel, shown as distributions without averaging across runs. **C)** DGEA cross-run consistency for varying numbers of DEGs. *N* is the number of cluster-pair contrasts summarized in each instability bin for the top-200 analysis. Top: The *y*-axis is the Jaccard similarity between the top-*K* genes produced by DGEA on the example run and the top-*K* genes produced by DGEA on another run using the best-matched clusters, averaged across all other valid runs. Bottom: The *y*-axis is the absolute rank difference between top-*K* DEGs found in both the example run and another run, averaged across all valid runs. **D)** Pearson correlation between cluster instability and cluster quality with respect to the reference annotation. Cluster quality is measured by Jaccard similarity, recall, or purity after matching each example-run cluster to its best-matching reference-annotation cluster. Dots are colored by the sample from which the cluster was obtained.

The best-matching clusters were used to evaluate whether DGEA produced consistent results across runs. DGEA was performed between pairs of clusters, drawn from the example run such that they had the same instability categorization. For example, DGEA would be performed between two clusters in the example run with mean instability score in bin [0, 0.1). Then, this contrast would be repeated for each of the remaining runs, using the respective best-match clusters. The result is one list of differentially expressed genes (DEGs) for each run (with some exclusion criteria, for example, the minimum cluster size was 100 spots; see Methods for details). We found that cluster instability was indicative of poor cross-run consistency in DGEA results and vice versa (Fig. 4C; Supplementary Figs. S20–S21). Gene-list agreement was highest for contrasts in the [0.0, 0.1) instability bin, with mean gene list Jaccard close to 1, and declined as instability increased. For example, at top 200, the mean gene list Jaccard decreased from *≈*0.98 in the lowest instability bin [0.0, 0.1) to*≈* 0.33 in the highest instability bin [0.5, 0.6). The complementary rank analysis agreed with this pattern: the mean absolute rank difference among shared top-200 genes increased from*≈* 2 positions to *≈*40 positions. Rank differences were calculated only for genes present in both top-*K* lists because gene-set disagreement was already quantified by Jaccard similarity; however, the shared genes retained their original ranks (i.e., shared genes were not re-ranked after restricting the intersection).

The number of top-ranked DEGs carried forward as markers varies with the downstream analysis. Compact lists of approximately 5–20 genes are commonly used for cluster annotation and focused interpretation, whereas larger signatures containing up to 200 genes may provide better performance for tasks such as cell-type characterization, classification, and deconvolution [49, 50]. We therefore evaluated cross-run consistency of the top 10, 20, 50, and 200 DE genes, spanning both compact marker sets and broader transcriptional signatures, although the former is more commonly combined with clustering methods. Cross-run gene list consistency varied with list length *K*. For stable clusters with instability scores in bin [0.1, 0.2), mean gene-list Jaccard increased from 0.75 for the top 10 genes to 0.81 for the top 200 genes. Meanwhile, the mean absolute rank difference increased from 1.18 to 16.25 positions. This increase reflects a scale effect where ranks span only 1–10 for a top-10 list but 1–200 for a top-200 list, so a similar proportional change in rank could produce a much larger absolute difference for the longer list. This increase may also reflect the inclusion of lower-ranked genes whose ordering is less consistent. Importantly, the same overall pattern was observed at every evaluated list length: higher-instability bins exhibited lower gene-list Jaccard and greater rank differences.

Lastly, we evaluated whether instability at the cluster-level was indicative of performance based on reference annotations. We aligned each cluster in the example run to the reference annotation, again selecting the best-match based on maximum Jaccard similarity. Taking SEDR as an example (Fig. 4D), higher cluster instability was negatively correlated with agreement to the reference annotations, based on Jaccard similarity (*r*^2^ = 0.52) and to a lesser extent recall (*r*^2^ = 0.31) and purity/precision (*r*^2^ = 0.17). Thus, clusters that were unstable across runs had worse agreement to the reference annotation; similar trends were observed for other methods (Supplementary Figs. S24–S26).

### InSTability is fast on large, high-resolution data sets

#### Instability map computation adds minimal overhead

Next, we evaluated the computation time required to run InSTability, as this is a limiting factor in incorporating the software tool into spatial transcriptomics workflows. We found that calculation of the instability maps from *R* = 50 partitions was extremely fast (Supplementary Table S6). Even on the largest VisiumHD data set, with 503, 622 spots, instability map computation took under 5 seconds, from end-to-end, including loading the input clustering results and writing the output file. Thus, instability map computation adds negligible overhead to spatial transcriptomics workflows. Runtimes were nearly identical across clustering methods applied to the same sample, indicating that the computational cost was driven primarily by sample size rather than by the partitions or the clustering method used to produce them.

The dominant cost of running InSTability was the time to run clustering. Clustering runtime varied substantially across methods and increased with dataset size (Supplementary Table S7). For standard Visium-sized datasets, mean clustering runtime ranged from a few seconds for Leiden, Louvain, and BayesSpace to tens or hundreds of seconds for GraphST, STAGATE, and SEDR, while SpiceMix required thousands of seconds, although we did not run it with optimal hardware specifications. Runtime increased dramatically with data set size, but for the largest Visium HD sample with 503, 622 spots, we could still run both Leiden and Louvain in ~0.66 hours.

#### Trends in clustering instability emerge after just 5 runs with scores stabilizing after 10-20

As the dominant cost of running InSTability is the time to run clustering methods, the number of runs needed is an important consideration. As an example, the time to run both Leiden and Louvain on the largest VisiumHD sample with 503, 622 spots, would be *<* 4 hours if *R* = 5 runs and *>* 30 hours for *R* = 50 runs. Therefore, we evaluated the sensitivity of our results to the number of clustering runs. Recalculating instability scores from 5 runs, and then adding 5 runs incrementally up to 50 clustering runs, showed spatial patterns observed in the 50-run analysis were visible at five runs and became increasingly consistent with the 50-run analysis as additional runs were included (Fig. 5A; Supplementary Figs. S28–S34).

**Fig. 5.**
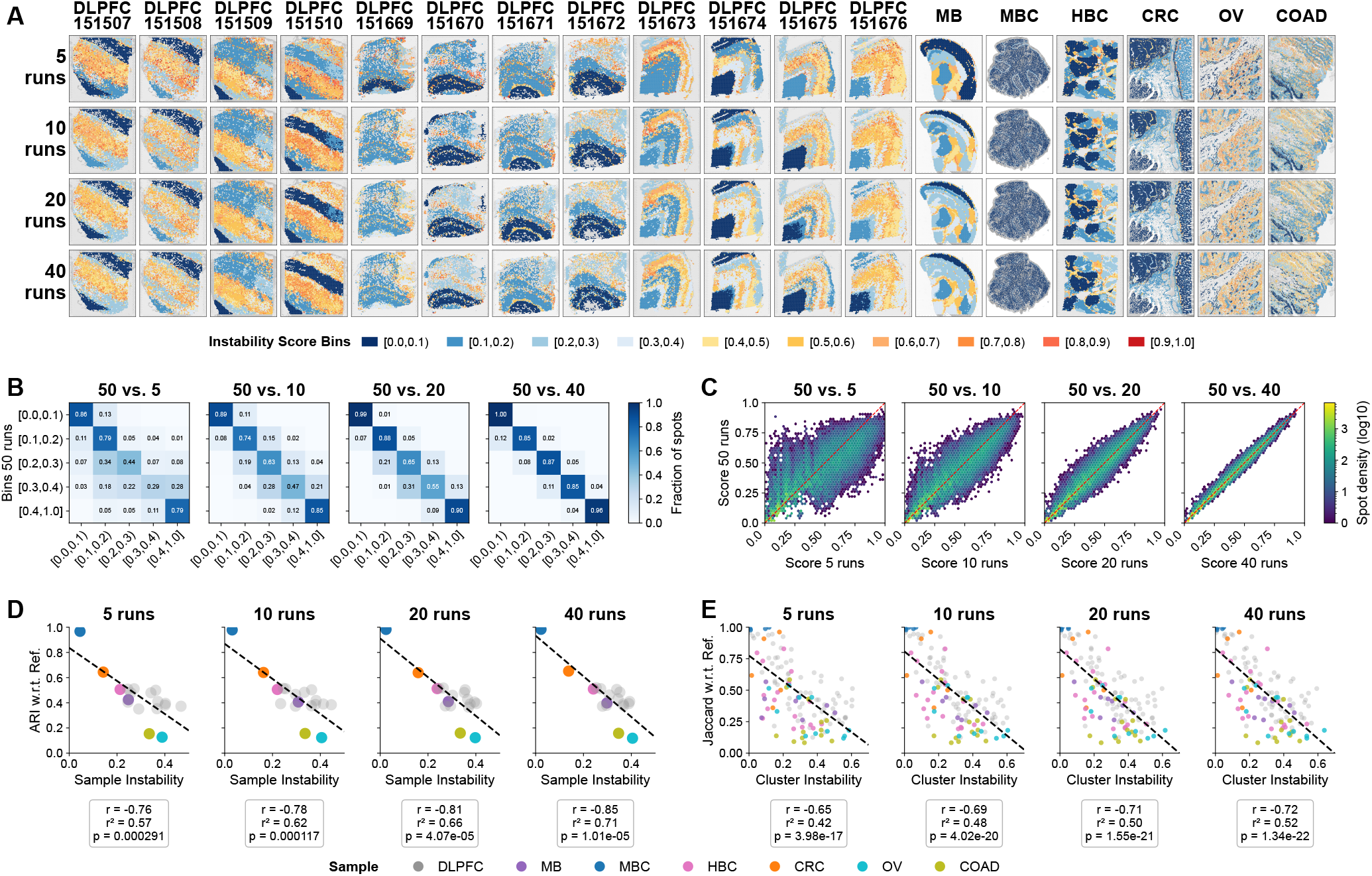
Trends in clustering stability emerge in as few as five runs and instability estimates progressively converge as additional runs are included, showing substantial convergence by 20 runs. All data shown are for SEDR. **A)** Spatial instability maps for SEDR computed from different numbers of runs (rows) for each of the 18 samples (columns). **B)** Comparison of instability-category assignments computed from 50 versus *R* runs. Each row shows the distribution of *R*-run categories among spots assigned to that category using 50 runs. Heatmap shows the fraction of spots assigned the category on the *y*-axis for 50 runs and assigned the category on the *x*-axis for *R* runs (rows sum to 1). Diagonal boxes indicate no change in score category. **C)** Comparison of spot-level instability scores computed from 50 runs (y-axis) and *R* runs (x-axis). Color indicates spot density on a log_10_ scale (i.e., a colorbar value of *k* corresponds to 10^*k*^ spots). Spots on the line have identical scores. **D)** Pearson correlation of sample accuracy (mean ARI across the *R* runs) versus sample instability (mean score across spots) for each number of runs. Dots are colored according to sample. **E)** Pearson correlation of cluster quality (Jaccard), evaluated by matching each cluster in the 50-run-defined example run to the reference annotation, versus cluster instability for each number of runs. Dots are colored according to the sample from which the cluster was obtained. Corresponding analyses for the other clustering methods are shown in Supplementary Figures S28–S34.

Discretized instability score assignments showed progressive convergence, taking the 50-run analysis as the convergence reference (Fig. 5B). For 5 runs, the number of spots with matching score categories as the 50-run analysis ranged from ~0.29 to ~0.86, with the greatest disagreement occurring among intermediate instability categories. This range improved to 0.47–0.89 with 10 runs, 0.55–0.99 with 20 runs, and 0.85–1.00 with 40 runs. The slower apparent convergence of the intermediate categories likely reflects their sensitivity to bin boundaries, as small numerical changes can move spots between adjacent categories even when their continuous instability scores are similar.

Instability scores themselves also converged to the 50-run analysis, becoming progressively concentrated around the identity line, with a marked reduction in dispersion between 10 and 20 runs and near agreement by 40 runs (Fig. 5C). Thus, a small number of runs was sufficient to identify the major spatial trends, whereas 10 *−*20 runs produced estimates substantially closer to the 50-run reference. A complementary sensitivity analysis using 25 random *R*-run subsets drawn from the same set of 50 runs showed a nearly identical convergence pattern, indicating that the observed convergence was not specific to the ordering of clustering runs (Supplementary Fig. S27).

Importantly, the relationship between instability and clustering performance (i.e., agreement against reference annotations) was also detectable with as few as 5 runs. At the sample level, instability was negatively correlated with ARI at 5 runs (*r* = *−*0.76, *p* = 2.91 *×* 10^*−*4^), and the association strengthened gradually to *r* = *−*0.85 at 40 runs (*p* = 1.01 *×* 10^*−*5^; Fig. 5D). Likewise, cluster-level instability was negatively correlated with cluster accuracy measured by Jaccard similarity at 5 runs (*r* = *−*0.65, *p* = 3.98 *×*10^*−*17^), increasing in magnitude to *r* = *−*0.72 at 40 runs (*p* = 1.34*×* 10^*−*22^; Fig. 5E). Together, these results indicate that the principal spatial and accuracy-related trends can be recovered from as few as 5 runs, making it computationally feasible to incorporate InSTability into spatial transcriptomics workflows, especially those based on fast graph community detection methods.

## Discussion

In this work, we introduced InSTability, a general computational framework for quantifying differences in partitions derived from ST datasets at the observation-level. We then demonstrated the effectiveness of this framework for localizing cluster instability due to algorithmic perturbations in an extensive benchmarking study of 7 methods and 18 samples. Clustering instability maps incurred little computational overhead compared to clustering, with key trends emerging in as few as five independent runs, which can be executed in parallel. Thus, InSTability is fast, even on large VisiumHD datasets, and can strengthen spatial transcriptomics workflows in the following ways.

### Using InSTability for reference-free evaluation of clustering quality

A key finding of our study is that clustering instability correlates with poor agreement with reference annotations for many methods tested. Thus, researchers can leverage InSTability to gauge whether a particular clustering analysis is likely to be of poor quality in the absence of ground truth. InSTability provides the greatest utility for this purpose when clusters do not display other signs of being poor quality, such as spotty-ness or spatial incoherence. As an example, clusters produced by SEDR on DLPFC sample 151674 appear spatially coherent but are unstable and have poor agreement with the reference annotation (Fig. 2). An important caveat when using InSTability is that the correlation strength depends on the method; we therefore recommend pairing InSTability with 5–10 runs of SEDR, Leiden, Louvain, or GraphST.

### Using InSTability for reference-free evaluation of clustering difficulty

Our study also revealed that clustering instability varies across tissue sections, with shared spatial patterns emerging among many methods. As an example, considerable heterogeneity was apparent among the individual DLPFC samples, with large regions of low and high instability shared across multiple clustering methods. Additionally, clustering instability correlated with sequencing technologies for SEDR: Slide-seq resulted in the best stability and Visium HD with 8 *×*8 micron bins resulted in the worst (tied with Visium on some of the DLPFC samples). This result is expected as dataset-specific factors such as transcriptomic complexity, spatial organization, signal-to-noise ratio, and the separability of spatial domains may jointly influence clustering sensitivity to algorithmic perturbations and challenge the accuracy of clustering methods. For Visium HD, this issue is particularly relevant because the choice of bin size changes both the amount of transcriptomic information available per observation and the spatial scale at which tissue structure is represented, with decreased resolution leading to higher numbers of transcripts but also cellular mixtures. In such contexts, researchers can leverage InSTability, paired with just 5–10 runs of Leiden or Louvain to quickly gauge clustering difficulty for a particular dataset after preprocessing.

### Using instability to guide ST analyses

Our study demonstrates that clustering instability propagates to downstream analyses, as may be expected. With as few as five repeated runs, instability scores can be used to identify clusters for which downstream analyses may be sensitive to the seed choice. Stable clusters, with mean scores between 0.0 and 0.1, are unlikely to yield different downstream results across runs, whereas unstable clusters, with mean scores greater than 0.4, are more likely to be affected. Clusters with intermediate instability showed greater variability in how sensitive downstream results were to the seed choice and thus may require further validation. We encourage researchers to report the clustering instability map alongside the final clustering result used for downstream analyses for greater transparency.

High clustering instability should not, however, be interpreted as evidence that a cluster is biologically meaningless, as a highly unstable analysis may produce a favorable partition under a particular seed. Rather, high instability is an indication that the inferred domains warrant further validation, for example review by pathologists. High instability could also indicate that the tissue does not readily decompose into discrete domains. In such settings, methods that explicitly model continuous spatial variation or jointly represent spatial gradients and domains may provide complementary descriptions of tissue organization [51, 52] If clustering was based on gene expression alone, approaches that incorporate histological information, including spatial domain segmentation methods [53, 54] and cell type prediction methods [55], could be explored as an alternative. Additional analyses that do not require discrete clusters, such as spatial autocorrelation analyses (e.g., those implemented in MERINGUE [56] InSTAnT [57]) may provide a more direct way to test spatially structured biological hypotheses.

### Using InSTability with soft clustering methods

Our instability framework was primarily designed for clustering methods that do not provide their own estimates of uncertainty, such as community detection methods Leiden and Louvain. However, we included BayesSpace as an informative comparison because its probabilistic clustering framework allows uncertainty to be calculated directly as the entropy of posterior cluster-assignment probabilities. Entropy and instability are distinct: entropy characterizes ambiguity in the cluster assignment of an individual spot within a fitted model, whereas instability measures whether cluster assignments are reproduced across repeated stochastic runs. The two measures need not identify the same spots, as we observed from our application of InSTability to BayesSpace.

In BayesSpace, low-entropy spots were concentrated in cluster interiors and high entropy spots were concentrated near inferred cluster boundaries, which did not always coincide with reference tissue annotations, such that changing these assignments would improve agreement with the reference. We found that high entropy spots could have low instability, indicating that the conversion to hard cluster assignments was consistent across seeds. As a result, BayesSpace was generally more stable than the other methods tested, although greater stability did not translate to greater agreement with reference annotations, resulting in a weak inverse relationship between instability and reference agreement. We therefore do not recommend using InSTability to assess the potential quality of BayesSpace clustering. InSTability could still be used to gauge the sensitivity of downstream analyses to seed-dependent variation, although our study suggests this would be limited for BayesSpace.

### Using InSTability in ST method benchmarking and development

In our study, the tissue section being analyzed appeared to have the strongest influence on clustering instability, but method choice still mattered and could alter its spatial pattern; for example, changing SEDR’s backend from Leiden to mclust concentrated instability at cluster boundaries, while improving agreement against reference annotations. Such comparisons provide a way to examine how methodological choices—including spatial smoothing, representation learning, probabilistic modeling, and optimization—shape both clustering stability and performance, and raise questions of how stability should be considered in method development. Robustness to seed choice is desirable because it promotes reproducibility and can be associated with quality with respect to reference annotations, but stability alone does not imply quality: a method could consistently produce the same biologically meaningless cluster. Conversely, instability may be informative in regions where the underlying data do not support a single partition so clustering is ambiguous. An ideal method would exhibit greater instability where the inferred structure is ambiguous and exhibit stability where the inferred cluster structure is well supported by the data and ideally biology, although the appropriate level of granularity must also be considered, for example clusters having high purity but low recall with respect to reference annotations. Thus, InSTability can complement traditional reference-based measures in method benchmarking by assessing whether a method’s stability reflects its performance.

### Study Limitations

#### Parameter tuning was limited

Some prior benchmarking and method-development studies have used reference annotations to guide parameter selection, for example choosing settings that maximize ARI relative to reference annotations [58–60]. Instead, we performed limited parameter tuning for each method–sample configuration, with the goal of ensuring that clustering granularity was close to the number of labels in the reference. Our tuning procedure reflects practical scenarios, where researchers do not have access to reference annotations but have a target for the number of reference labels.

#### Reference annotations are not unique biological ground truth

Our study leveraged reference annotations as a quality metric, although they do not reflect a unique biological ground truth. Disagreement with a reference annotation can reflect differences in the granularity of the biological structures being represented, especially if the annotation convention gives transcriptionally similar but spatially disconnected regions distinct labels. Conversely, low ARI can indicate failures to recover annotated tissue structures in ways that cannot be reconciled through cluster splitting or merging and thus are genuinely inaccurate, as observed in DLPFC samples (e.g., Fig. 2). Thus, reference-based quality metrics continue to be an important aspect of benchmarking clustering performance.

### Future Work

Our study also raises several opportunities for future work leveraging InSTability. Because we evaluated each clustering method in isolation, an important next step is to examine how choices throughout the data processing stack influence clustering instability. These choices begin with how the data are represented: for Visium HD data, researchers can select different bin resolutions, while nuclei segmentation maps transcripts from bins to nuclei, both of which change transcript counts and expression mixtures at the observation level. Subsequent steps require additional choices, such as the number of nearest neighbors for graph-based methods. Finally, clustering itself involves choices, such as the resolution parameter, which controls the granularity of the initial partition, which are sometimes combined with postprocessing steps, such as merging statistically indistinguishable clusters, as implemented in the Space Ranger pipeline. These choices have the potential to affect clustering stability and quality. Thus, combining InSTability with reference-based evaluation frameworks could identify analytical choices that produce more stable and biologically meaningful results, informing best practices.

Another direction for future work is to leverage InSTability on methods that leverage histological images, in addition to gene expression data. Imaging data, increasingly used in spatial domain segmentation [61–63] and cell type prediction [55], may improve stability. Cell type prediction methods, in particular, report probabilities for individual class assignments, providing a measure of uncertainty for each prediction [64, 65]. InSTability complements entropy from cell type prediction methods by assessing whether the same groups of spots or nuclei receive the same cell type labels across repeated runs. Lastly, InSTability makes no assumptions about how input partitions are generated; thus, it can be used to compare partitions produced by different methods or analytical workflows, in addition to run-to-run variation within a single method, under either algorithmic or data perturbations. In this way, InSTability provides a framework for understanding where in the tissue consistency breaks down, thereby indicating where further validation or alternative analyses beyond clustering may be warranted.

## Methods

### Instability Score for Spatial Mapping

Our per-spot instability score *u*_*i*_ captures the variability of spot *i*’s cluster assignment across the *R* input partitions, which could be created through repeated runs of the same method on the same input, differing only by the random seed (Fig. 1). As previously mentioned, a spot’s cluster assignment is *stable* when the spots that cluster with it in one run tend to cluster with it in the other runs, while spots assigned to different clusters tend to remain separate; otherwise, the cluster assignment is *unstable*. We quantify this behavior using spotwise co-association across partitions. The *n× n* co-association matrix contains the fraction of times that spots *i* and *j* cluster together across the *R* partitions, i.e.,

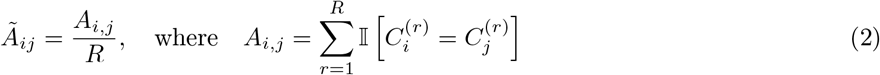

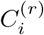 is the cluster label of spot *i* in partition *r*, and I denotes the indicator function. We then consider *intra*-cluster and *inter*-cluster co-association for spot *i* with respect to partition *r ∈ {*1, 2, …, *R}* as

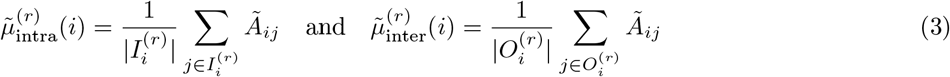

where 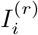 is the set of spots assigned to the same cluster as *i* in partition *r*, excluding *i*, and 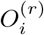 is the set of spots assigned to a different cluster from *i* in partition *r* (i.e., the *complement* of the cluster containing spot *i* in partition *r*).

The instability score of spot *i* is formulated from the *difference* between spot *i*’s intra-cluster and inter-cluster co-association, both with respect to run *r*. Because partition *r* itself contributes to the co-association matrix, pairs that are assigned to the same cluster in partition *r* receive a deterministic contribution of 1*/R*, whereas pairs assigned to different clusters receive no contribution from that partition. Without correction, this induces a finite-*R* downward bias. We therefore apply the normalization factor *R/*(*R −*1), defining the partition-specific spot-level instability score as:

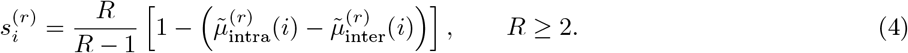

The normalization factor is equivalent to excluding partition *r* when computing the co-association structure against the clustering assignment from that partition, i.e., substituting *Ã*_*i,j*_ with 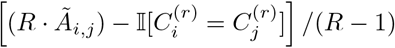 in Equation 3. Thus, each clustering result is effectively evaluated against the consensus of the remaining *R –* 1 partitions rather than all *R* partitions including itself. Finally, averaging the 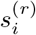 values across partitions gives the local instability score for spot *i*:

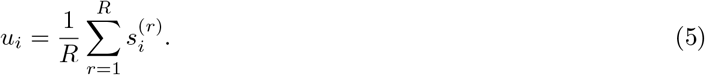

Spotwise scores *u*_*i*_ provide a spatially-resolved measure of clustering instability across the input partitions.

#### Degenerate cases

Intra-cluster co-association 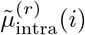 is undefined when the cluster containing spot *i* is a singleton cluster in partition *r*, and inter-cluster co-association 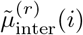 is undefined when partition *r* is just a single cluster. In these degenerate cases, we use the following conventions: if spot *i* forms a singleton cluster in run *r*, we set 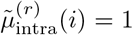, and if all spots belong to a single cluster in run *r*, we set 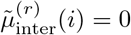. These conventions ensure that the per-spot instability score remains well-defined while retaining all available co-association information on the nondegenerate side of the comparison. In particular, if all partitions produce the same degenerate partition—for example, all singleton clusters in every partition or a single global cluster in every partition—then the resulting per-spot instability scores are 0 for all spots, since the clustering output is perfectly consistent across partitions.

### Analysis of instability score range

Although both 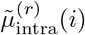 and 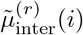 lie in [0, 1], tighter bounds follow from the fact that the run *r* is included in *Ã*. For any non-singleton cluster, every spot assigned to the same cluster as *i* in run *r* co-clusters with *i* in at least that run and no more than all *R* partitions, implying

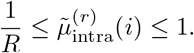

Similarly, a spot assigned to a different cluster from *i* in run *r* can co-cluster with *i* in 0 of the remaining runs and at most the remaining *R −* 1 runs, so

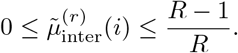

Putting these bounds together gives

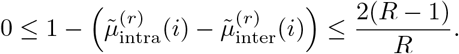

and multiplication by *R/*(*R −*1) gives 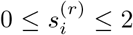. The same bounds are preserved under averaging 0 *≤u*_*i*_ *≤*2. The conventions defined above for singleton clusters and single-cluster partitions also preserve these bounds.

#### Interpretation of extreme cases

The lower bound 0 corresponds to perfect stability. When the cluster containing spot *i* is the same across all partitions, we get 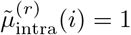 and 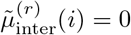. At the other extreme, the theoretical upper bound 2 corresponds to complete contradiction between a run-specific assignment and the consensus of the remaining runs. When spots assigned to the same cluster as *i* in run *r* never co-cluster with *i* in any of the remaining runs, and spots assigned to different clusters always co-cluster with *i* in the remaining runs, we get

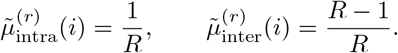

Such a configuration represents a pathological worst-case scenario and is not expected for meaningful clustering outputs.

For nondegenerate clusters, an intermediate reference point is random clustering across partitions. Suppose cluster assignments in the remaining partitions are effectively independent of the run *r*, and let *p* denote the probability that any two spots cluster together. Because the partition *r* necessarily contributes one co-association count to intra-cluster pairs *A*_*i,j*_ and zero to inter-cluster pairs *A*_*i,j*_,

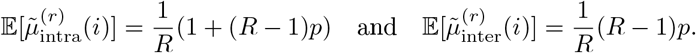

Their expected difference is thus 1*/R*, giving 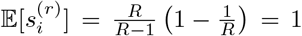. Thus, when partitions are generated via random clustering, the instability score has a reference level of 1 regardless of the number of clustering runs. A score near 1 can therefore be interpreted as indicating little structured cross-run agreement, approaching the behavior expected under effectively random assignment.

#### Practically, 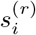 values are expected to lie predominantly in [0, 1]

By definition, 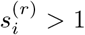 1 if and only if 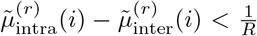. The value 1*/R* represents the deterministic within-cluster contribution from partition *r* itself. Therefore, a score greater than 1 indicates that, beyond this self-contribution, spot *i* shows greater agreement across the remaining partitions with spots outside its cluster in partition *r* than with spots inside it. This represents a locally anti-consensus assignment. In meaningful clustering solutions, the agreement with spots inside the reference cluster is generally expected to remain stronger than the agreement with spots outside it after accounting for this self-contribution, leading to 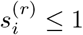 for most spots. Stable cluster cores typically have a large difference between 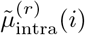 and 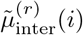, producing scores close to 0, whereas regions with little structured cross-run agreement have 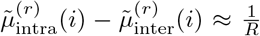, producing scores near 1. Values above 1 indicate stronger local disagreement than would be expected under effectively random assignment.

### Linear time and space algorithm for computing spatial instability maps

A naive implementation of the instability score requires the *n× n* count co-association matrix *A*, resulting in Θ(*n*^2^) storage and Θ(*Rn*^2^) time complexity. Our implementation avoids explicit construction of the co-association matrix when computing intra- and inter-cluster co-association. The former can be written as

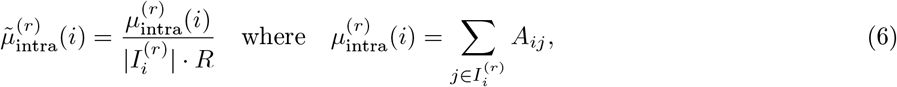

and similarly for the latter. The computation can then be restructured

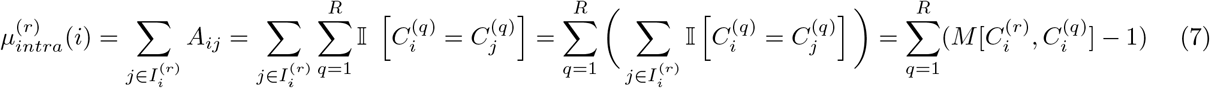

where 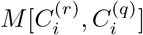 is the overlap between the cluster containing spot *i* in partition *r* and the cluster containing spot *i* in partition *q*. Cluster overlaps for a fixed pair of partitions (*r, q*) can be computed in *O*(*n*) time and space by looping over each spot and counting its cluster-pair membership on the fly.

Leveraging this cluster-pair membership computation, the auxiliary quantity 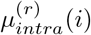 can be computed in Θ(*R*^2^*n*) time and Θ(*Rn*) space; see lines 1–26 of Algorithm 1 in the Supplemental Methods. During this preprocessing phase, the size of every cluster *c* in every partition *q* is stored as auxiliary quantity *N* ^(*q*)^[*c*]. Together, 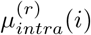 and *N* ^(*r*)^[*c*] enable us to compute any intra-cluster co-association term 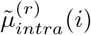 on the fly via Equation 7, since 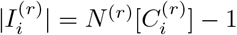.

Likewise, the inter-cluster co-association term 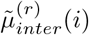 can be computed from the corresponding count 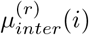 with 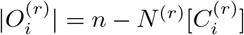. Computing 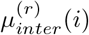 requires one additional auxiliary quantity: the row sums for the co-association count matrix *A*. The intra- and inter-cluster co-association counts for spot *i*, with respect to any run *r*, split the row sum

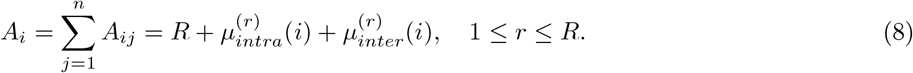

where *R* is the contribution from *A*_*ii*_. The row sum *A*_*i*_ can be computed from *N* ^(*q*)^[*c*] during the preprocessing phase, since it equals the total size of the clusters containing spot *i* across all runs.

After preprocessing, the instability score *u*_*i*_ can be computed for every spot *i* via Equations 4–5 in Θ(*nR*) time and Θ(*n*) space; see lines 27–44 of Algorithm 1 in the Supplemental Methods. The total time and storage complexity is Θ(*R*^2^*n*) and Θ(*Rn*), respectively. Related techniques have been used in prior work to improve the computational efficiency of algorithms for comparing partitions, such as the adjusted Rand index [66].

In comparison to the naive implementation, our algorithm substitutes a factor of *n* with a factor of *R* in both time and space complexity; this trade-off is favorable because *R ≪n* in our experiments. The largest Visium HD sample we analyzed had *n* = 503,622 spots, but *R* was capped at 50 runs, making *R* approximately four orders of magnitude smaller than *n*. Moreover, we can treat the number of runs as a fixed constant, which is reasonable as our study recommends just 5–10 runs regardless of the dataset, giving our algorithm linear time and storage complexity.

### Datasets

We evaluated InSTability on 18 samples drawn from seven publicly available spatial transcriptomics datasets; see the Supplemental Methods for detailed descriptions. All datasets were represented as AnnData objects and analyzed using Scanpy [1]. Each AnnData object contained a gene-expression matrix with spatial locations as rows and retained genes as columns, together with spatial coordinates and reference annotations. Histological images were retained for visualization only. Genes detected in fewer than 20 spatial locations were removed, mitochondrial genes were removed, and spatial locations were filtered based on minimum total UMI count and maximum mitochondrial gene percentage thresholds. Dataset-specific thresholds were chosen to accommodate differences in sequencing depth, tissue type, and spatial resolution. After quality control, CRC, COAD, and OV datasets were cropped, defining a spatially contiguous subset of spots, to ensure that all seven methods could be run for the primary instability benchmark. The full-resolution, quality-controlled, datasets were retained for scalability analyses using Leiden and Louvain. The quality-controlled datasets (AnnData objects), summarized in Table 1, served as the common input for clustering analyses, after method-specific preprocessing.

**Table 1.** Summary of spatial transcriptomics datasets used in the study. Primary benchmark comprised 18 samples.Full-resolution quality-controlled Visium HD datasets were additionally used for computational scalability analyses with Leiden and Louvain only.

| Dataset | Platform | Resolution | N | # Obs.<br>Retained | # Genes<br>Retained | # Ref.<br>Domains | UMI<br>threshold | Mito. gene<br>threshold |
| --- | --- | --- | --- | --- | --- | --- | --- | --- |
| **Primary benchmark** |  |  |  |  |  |  |  |  |
| MBC [40] | Slide-seq | locations | 1 | 25,548 | 11,970 | 8 | 200 | 20% |
| MB [39] | Visium FFPE | spots | 1 | 2,264 | 14,022 | 10 | 500 | 25% |
| DLPFC [37] | Visium | spots | 12 | 2,731–4,696 | 13,821–15,875 | 5 or 7* | 600 | 25% |
| HBC [38] | Visium | spots | 1 | 3,795 | 18,233 | 20 | 1,000 | 20% |
| CRC [41] | Visium HD | 16 μm bins | 1 | 21,656 | 17,823 | 6 | – | – |
| COAD [42] | Visium HD FFPE | 8 μm bins | 1 | 27,494 | 17,757 | 15 | – | – |
| OV [42] | Visium HD FFPE | 8 μm bins | 1 | 27,291 | 17,757 | 13 | – | – |
| **Full-resolution Visium HD datasets used for scalability analyses** |  |  |  |  |  |  |  |  |
| CRC | Visium HD | 16 μm bins | 1 | 110,402 | 17,823 | 6 | 400 | 20% |
| COAD | Visium HD FFPE | 8 μm bins | 1 | 503,622 | 17,757 | 16 | 200 | 20% |
| OV | Visium HD FFPE | 8 μm bins | 1 | 410,006 | 17,757 | 13 | 200 | 20% |

\*DLPFC sections 151669–151672 contained five reference domains, and the remaining eight sections contained seven (a small number of DLPFC spots lacked reference annotations and thus were excluded from analyses requiring reference labels, although they were retained for clustering and instability analyses). N: Number of samples; Obs: Observations (i.e., spatial locations); MBC: Mouse Cerebellum; MB: Adult Mouse Brain; HBC: Human Breast Cancer; DLPFC: dorsolateral prefrontal cortex; CRC: colorectal cancer; COAD: colon adenocarcinoma; OV: ovarian cancer.

### Clustering Methods

We evaluated seven ST clustering methods spanning several commonly used approaches, summarized in Table 2. Unless method-specific preprocessing was required, highly variable genes were selected from raw counts using the seurat_v3 method, expression was normalized to a total of 10^4^ per spot, log1p-transformed, and scaled with a maximum absolute value of 10. This preprocessing follows a commonly used Scanpy-style workflow and closely matches preprocessing used by several of the spatial methods evaluated in this study. BayesSpace and SpiceMix were run with their own method-specific preprocessing procedures. We followed parameter choices from the original software implementations or author-provided tutorials when practical, while maintaining a common analysis framework across methods. If an implementation or tutorial specified a seed for reproducibility, we used that value as the base seed. We set the target number of clusters to match the number of reference domains. Scripts to run clustering methods and make figures are available on Github https://github.com/ws500981/InSTability_figures.

**Table 2.** Summary of clustering methods evaluated in this study Seed dependence refers to the implementations used in this study, which contained stochastic components and were therefore evaluated across repeated runs.

| Method (Year) | Method Family | Core Modeling Idea | Reason for Inclusion | Seed-Dependent |
| --- | --- | --- | --- | --- |
| Leiden (2019) [43] | Graph community detection | Community detection with local moving, partition refinement, and graph aggregation | Widely used non-spatial graph-clustering baseline | Yes |
| Louvain (2008) [11] | Graph community detection | Greedy modularity optimization for graph community detection | Classical non-spatial graph-clustering baseline, used in Space Ranger | Yes |
| BayesSpace (2021) [29] | Bayesian spatial modeling | Bayesian clustering of low-dimensional expression with a spatial Potts-model prior | Representative Bayesian spatial-clustering method | Yes |
| STAGATE (2022) [44] | Deep graph learning | Graph attention autoencoder integrating gene expression with spatial-neighborhood information | Representative graph-attention spatial method | Yes |
| SEDR (2024) [45] | Deep graph learning | Masked deep autoencoder coupled with a variational graph autoencoder for joint expression-spatial representation learning | Representative deep spatial-representation method | Yes |
| GraphST (2023) [46] | Deep graph learning | Graph neural network with self-supervised contrastive representation learning | Representative self-supervised graph method | Yes |
| SpiceMix (2023) [47] | Probabilistic spatial modeling | Probabilistic latent-variable modeling of gene expression and spatial relationships | Representative probabilistic spatial-modeling method | Yes |

### Leiden and Louvain

Leiden [43] and Louvain [11] are graph-based community-detection algorithms. Louvain performs iterative modularity optimization, whereas Leiden adds a partition-refinement procedure that improves the connectivity of the resulting communities. For both standalone methods, we used the implementations in Scanpy. To construct the input graph, we selected the top 3, 000 variable genes, computed a 50-component PCA representation (seed was 42), and then constructed a 50-nearest-neighbor graph. The number of highly variable genes used is consistent with the feature-selection scale used in several spatial-transcriptomics workflows, including STAGATE and GraphST.

For each dataset, we selected the clustering resolution parameter by tuning to the number of reference domains, similar to the approaches used in SEDR, GraphST, and STAGATE. The general approach is to run the method with a resolution that produces a larger number of clusters than reference domains so that the resolution parameter can be incrementally decreased until the target number of clusters is hit. Leiden and Louvain were both run with seed 0 during tuning. We then ran Leiden (or Louvain) 50 times on the KNN graph using the tuned resolution parameter and seeds 42 + *t* for *t ∈* 0, …, 49.

### BayesSpace

BayesSpace [29] is a fully Bayesian spatial-clustering method that incorporates neighboring spots through a spatial Potts-model prior to encourage spatially coherent cluster assignments. To construct the input for BayesSpace, we used its native preprocessing workflow, which takes the top 2, 000 variable genes and 15 principal components (seed was 102). Both are defaults of the BayesSpace preprocessing implementation, and the authors specifically recommend using the top 15 principal components in most applications. For each dataset, we ran BayesSpace for 50 stochastic runs, setting the seed to 149 + *t* for *t∈{* 0, 1, …, 49*}*. Each clustering run used 10, 000 MCMC iterations with a 1, 000-iteration burn-in, which was the minimum recommended number of iterations for spot-level clustering. The number of clusters was set to the number of reference domains. BayesSpace outputs hard cluster labels by default, under `spatial.cluster`. We ran BayesSpace with `save.chain=True` so that the post-burn-in MCMC states (i.e., a collection of partitions with common cluster labels) could be used to compute spotwise cluster membership probabilities and entropy.

### STAGATE

STAGATE [44] uses a graph attention autoencoder to learn low-dimensional representations that jointly capture gene-expression and spatial-neighborhood information. To construct the input, we selected the top 3, 000 variable genes. For each dataset, we ran STAGATE for 50 stochastic runs, setting *t ∈{*0, 1, …, 49*}* to change the seed. The STAGATE method includes the following steps: (1) calculating STAGATE spatial network (we set `rad_cutoff=150`), (2) training the STAGATE autoencoder with seed 2020 + *t*, (3) applying the trained STAGATE model to produce a learned embedding with dimension 30, (4) constructing a 50-nearest-neighbor graph, (5) tuning the resolution parameter for Leiden setting the seed to 0 and the target to match the number of clusters in the reference (part of their tuning search function), (6) clustering with Leiden given the tuned resolution parameter and seed 2020. The STAGATE network used hidden dimensions [512,30] and was trained for 1, 000 epochs with a learning rate of 0.001, following the STAGATE tutorial.

### SEDR

SEDR [45] learns spatially informed representations by combining deep gene-expression representation learning with a variational graph autoencoder incorporating spatial structure. To construct the input, we selected the top 2, 000 variable genes, calculated 200 principal components using seed 42, and constructed the SEDR spatial graph using 12 neighbors; these choices reproduce those used in the authors’ 10x Visium clustering tutorial (note that expression normalization followed the shared preprocessing protocol described above). For each dataset, we ran SEDR for 50 stochastic runs, setting *t ∈{*0, 1, …, 49*}* to change the seed. The SEDR method includes the following steps: (0) initializing SEDR with seed 2023 + *t*, (1) training the SEDR autoencoder, (2) applying the trained SEDR model to produce a learned embedding, (3) constructing a 50-nearest-neighbor graph, (4) tuning the resolution parameter for Leiden setting the seed to 0 and the target to match the number of reference domains (note that we used our own tuning approach implemented for standalone Leiden because the range in their own tuning function was not exposed as user inputs), and (5) clustering with Leiden given the tuned resolution parameter and seed 2023 + *t*. We *re-ran* SEDR on each dataset replacing steps (4) and (5) above with mclust given seed 2023 + *t*, same as Leiden. The mclust method takes the number of mixture components as input, which we set to match the number of reference domains, instead of a resolution parameter.

### GraphST

GraphST [46] uses graph neural networks and self-supervised contrastive learning to learn spot representations that incorporate both gene-expression and spatial-neighborhood information. To construct the input, we selected the top 3, 000 variable genes. We trained GraphST directly on the preprocessed expression data without an additional PCA step, as the 3, 000-gene feature dimension is consistent with the GraphST preprocessing implementation. For each data set, we ran GraphST for 50 stochastic runs, setting *t ∈{*0, 1, …, 49*}* to change the seed. The GraphST method includes the following steps: (1) initializing GraphST with seed 41+*t*, (2) training the GraphST autoencoder on the input dataset, (3) clustering, which encompasses applying the trained GraphST model to produce a learned embedding, constructing a graph representation, tuning the resolution parameter for Leiden using the number of annotation categories as the target, and then clustering with Leiden. We set `refinement=False`, which disables the optional spatial label-refinement step in clustering, following the GraphST tutorial.

### SpiceMix

SpiceMix [47] is an unsupervised probabilistic latent-variable model that represents gene expression using latent metagenes while modeling spatial relationships among neighboring spots. For each data set, we ran SpiceMix for 50 stochastic runs, setting *t ∈{*0, 1, …, 49*}* to change the seed. Specifically, we set the numpy and pytorch seeds, both to *t*, as numpy and pytorch are invoked by SpiceMix, and then ran SpiceMix commands with default parameters, unless otherwise noted, following the tutorial. SpiceMix executes a custom Louvain-based clustering method with built-in clustering resolution tuning over the default range of 0.5–2.0. The number of latent factors was 10 for all datasets (this is slightly smaller than the default of 15 but was necessary to improve speed and memory given our compute resources). Model fitting used 10 initialization iterations followed by 200 spatial optimization iterations, consistent with iteration settings documented in the SpiceMix repository.

### Evaluation Metrics

#### Instability Maps

Instability maps were visualized for each sample-method configuration (Fig. 2; Supplementary Figs. S1–S17). We also visualized the reference annotations and predicted clusters from the example run of each method. Colors of the predicted clusters were aligned to the reference colors to facilitate visual comparison of spatial patterns. A Jaccard-overlap matrix was constructed between all predicted example-run clusters and all clusters in the reference annotation. A globally optimal one-to-one assignment was obtained using the Hungarian algorithm by maximizing total Jaccard overlap. Predicted clusters assigned to a reference group inherited that group’s color, whereas additional unmatched predicted clusters were assigned unused colors from the same cluster palette. This alignment was used only for visualization and did not relabel clusters or affect ARI, cluster-quality metrics, cross-run matching, DGEA, or instability calculations.

Instability was summarized at both the spot and cluster levels. At the spot level, the proportion of spots in each 0.1-wide instability bin was calculated. At the cluster level, instability was averaged over all spots belonging to each cluster in the example run, 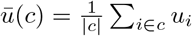, and the resulting cluster mean was used to assign the cluster to a corresponding instability bin. In the visualization, spot-level distributions were shown as percentages, whereas cluster-level summaries represented the number of clusters in the example run in each bin (Panel D of Fig. 2 and Supplementary Figs. S1–S17). We also report the fraction of unstable spots as the proportion with instability *≥*0.4 (Supplementary Tables S1–S2). Method-level summaries averaged this fraction across available benchmark samples, whereas sample-level summaries averaged it across available clustering methods. Cluster counts and the fractions of clusters in the four broader instability categories were calculated from the corresponding 50-run example-run clusters.

#### Adjusted Rand Index and Sample-Level Association

Clustering accuracy with respect to available reference annotations was quantified using the Adjusted Rand Index (ARI). For each sample, method, and stochastic run, ARI was calculated between the predicted labels and the reference annotations after excluding spots with missing reference labels (i.e. if reference labels are “N.A.”). For an analysis using *R* runs, clustering performance was summarized as the arithmetic mean of the *R* run-specific ARI values. Note that we used *R* = 50 for all analyses except for “Sensitivity to the number of clustering runs” analysis. Sample instability was summarized independently as the mean *R*-run instability score across all spots in the corresponding sample–method configuration. Within each clustering method, the association across samples was quantified using Pearson’s correlation coefficient *r*, with *r*^2^ and the corresponding *p*-value also reported (Fig. 3B, Panel D of Fig. 5 and Supplementary Figs. S28–S34, Supplementary Table S3).

#### Cluster-wise Quality Analysis

Because downstream interpretation frequently depends on individual clusters rather than an entire partition, we evaluated the relationship between cluster instability and several cluster-level quality measures. For the principal 50-run analyses, clusters were defined by the example run selected from all 50 runs. The same 50-run-selected example partition was also retained in the run-count sensitivity analysis so that differences across run counts reflected changes in the estimated instability rather than changes in the cluster partition being evaluated.

##### Concordance with reference annotations

For reference-based cluster metrics, spots with missing reference annotations were first excluded (i.e. reference label is not “N.A.”). Each predicted cluster was independently matched to the reference cluster having the highest Jaccard similarity (i.e., which is the number of spots common to both divided by the number of spots in the union of both clusters). Ties were broken among reference clusters by taking the one with the larger intersection. For each best-matched-reference and predicted cluster pair, we additionally computed purity and recall (Panel D of Fig. 4 and Supplementary Figs. S24–S26; Panel E of Fig. 5 and Supplementary Figs. S28–S34). Purity (i.e., precision) is the fraction of the predicted cluster recovered by the best-match reference cluster, recall is the fraction of the best-match reference cluster recovered by the predicted cluster, and Jaccard jointly penalizes both over- and under-extension relative to the matched annotation. Supplementary Table S1 reports method-level Pearson *r* and *r*^2^ values obtained by pooling eligible clusters across samples within each method.

##### Expression coherence and variance

Expression-based cluster quality was evaluated in a normalized PCA representation. A copy of the AnnData expression matrix was normalized to a total of 10,000 per spot and log-transformed. PCA was then fitted using up to 50 components, subject to the dimensions of the expression matrix, and the first 20 principal components were retained. Each resulting 20-dimensional spot vector was normalized to unit Euclidean norm. For a cluster containing at least two spots, expression coherence was defined as the average pairwise cosine similarity,

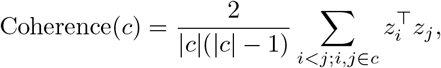

where *z*_*i*_ is the unit-normalized 20-dimensional PCA vector for spot *i*. Higher values therefore indicate greater similarity of the expression representations within a cluster. Within-cluster expression variance was calculated as the mean population variance across the same 20 normalized PCA dimensions,

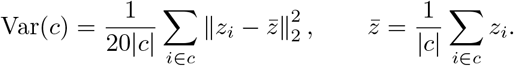

Lower values indicate a more compact cluster in the normalized PCA representation. The associations of cluster instability with expression coherence and expression variance across clustering methods are shown in Supplementary Figures S22–S23, respectively. Supplementary Table S1 additionally reports the pooled method-level Pearson *r* and *r*^2^ values for both expression metrics, together with those for purity, recall, and Jaccard.

#### Cross-run Cluster Consistency

We evaluated whether the mean instability score of each cluster from the example run reflected how reproducibly it could be recovered in the other stochastic runs. For each predicted cluster in the example run, one cluster is selected independently from each remaining run by maximum Jaccard similarity. Ties in maximum Jaccard were resolved by the largest intersection. No clusters were merged, and the mapping was not constrained to be one-to-one. Consequently, different clusters in the example run could have the same best-match cluster in a non-example run. For each cluster in an example run, the cross-run consistency measure was the mean best-match Jaccard across the remaining runs; the distribution is also shown (Fig. 4B and Supplementary Figure S19).

### Differential Expression Stability

To determine whether clustering instability propagated downstream, we evaluated the stability of differential gene expression analyses (DGEA) across stochastic clustering runs.

#### Cluster alignment and instability-stratified contrasts

Mean cluster instability was calculated for each cluster in the example run, binning each cluster to one of the ten 0.1-wide instability bins. A fixed set of candidate DGEA contrasts was then constructed by considering all unordered pairs of example-run clusters belonging to the same instability bin. A pair was retained only if both example-run clusters contained at least 100 spots. The fixed example-run contrast plan was then applied to each remaining run using the best-match clusters described above. A run-specific contrast was excluded if both example clusters matched to the same run cluster, producing a matching collision, or if either matched run cluster contained fewer than 100 spots. Matching collisions were rare, occurring in less than 1% of contrast–run comparisons overall. No matching collisions occurred for the SEDR HBC analysis shown in Figure 4A. Note that cluster matching and DGEA were based on the clustering assignments themselves and did not require reference annotations.

#### Pairwise differential gene expression analysis (DGEA)

For DGEA, we implemented the preprocessing protocol from [67]: expression values were normalized to 10,000 counts per spot, log-transformed, and restricted to genes detected in at least one spot. Differential expression was performed using the Wilcoxon test implemented in `scanpy.tl.rank_genes_groups`, with absolute-rank ordering, tie correction enabled, and *p*-values FDR-corrected with the Benjamini–Hochberg method. Differentially expressed genes were retained when the adjusted *p*-value was *<* 0.05, the absolute log fold-change was*≥* 0.5 (similar to [68]), and the gene was expressed in more than 10% of spots in at least one of the two groups being tested (similar to [69]). Retained genes were ordered by the absolute Wilcoxon statistic and truncated to at most 200 genes.

#### Cross-run DGEA consistency

For each retained cluster-pair contrast, defined from the example run, the DGEA result (ranked gene list) from the example run served as the reference and was compared to the DGEA result (ranked gene list) from each remaining run. Cross-run consistency was calculated using Top-*K* gene lists with 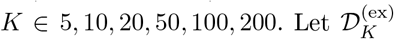 and 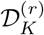 denote the Top-*K* filtered DE-gene sets from the example run and non-example run *r*, respectively. Gene-set reproducibility was quantified using the Jaccard index,

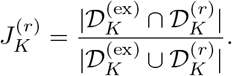

For each contrast and value of *K*, Jaccard values were averaged across non-example runs (Fig. 4C; Supplementary Fig. S20). Gene set rank-displacement metric was calculated only for genes shared by the two Top-*K* lists. Let 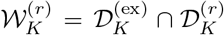 denote the set of shared genes. When at least one gene was shared, the run-specific mean absolute rank difference was

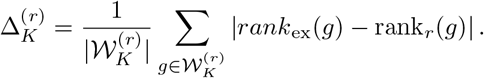

These run-specific values were then averaged across valid non-example runs for each contrast. Genes absent from either Top-*K* list were excluded from this displayed rank-difference calculation (Fig. 4C; Supplementary Fig. S21).

### Soft-label Entropy Comparison

Soft-assignment entropy was evaluated specifically for BayesSpace, which provides run-specific soft cluster-assignment weights. For a BayesSpace run with soft-assignment matrix *P ∈*ℝ^*n×M*^, where *M* denotes the total number of clusters in that run, the soft-assignment weights for each spot were normalized to sum to one so that they formed a probability distribution across the *M* clusters:

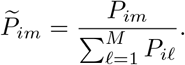

Spot-level Shannon entropy was calculated as

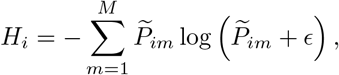

where *ϵ* = 10^*−*12^ was included for numerical stability. Entropy was retained on its natural, unnormalized scale rather than divided by log *M*. Low entropy indicates concentration of the assignment weights on one cluster, whereas high entropy indicates greater ambiguity among multiple clusters.

Two complementary entropy summaries were used. For the spatial comparison in Supplementary Fig. S18, entropy was calculated for the selected BayesSpace example run and visualized alongside the BayesSpace instability map. Its spot-level association with BayesSpace instability was summarized using both Spearman and Pearson correlations; the same entropy values were also compared with Leiden instability. For the sample-level BayesSpace analysis in Fig. 3B, entropy was first averaged across spots within each stochastic run and then averaged across the available runs. Thus, the 50-run sample entropy used in the main comparison and Supplementary Table S5 represents the mean of the 50 run-specific mean entropies rather than the entropy from the example run alone. Its association with mean ARI across the 50 BayesSpace runs was assessed using Pearson correlation.

### Sensitivity to the Number and Selection of Runs

We examined whether the estimated instability and its relationships with clustering quality were sensitive to the number and selection of stochastic runs.

#### Nested run-count analysis

For the main run-count analysis, *R*-run results were constructed from nested prefixes of the original stochastic runs, i.e., the 5-run analysis used the first five runs, the 10-run analysis used the first ten runs, and so forth. The principal visual comparisons used *R∈ {*5, 10, 20, 40 *}*, with the 50-run result serving as the reference.

Spatial maps at each *R* used the same ten 0.1-wide instability display bins as the principal visualization (Panel A of Fig. 5 and Supplementary Figs. S28–S34). To summarize movement between instability categories as *R* increased, transition analyses used five displayed categories, [0, 0.1), [0.1, 0.2), [0.2, 0.3), [0.3, 0.4), [0.4, 1.0]. For each *R*, spots were cross-tabulated according to their 50-run category and their *R*-run category. Transition counts were pooled across the analyzed biological samples for each method and row-normalized by the 50-run category, so each row represents the fraction of spots from a given 50-run instability category assigned to each category when fewer runs were used (Panel B of Fig. 5 and Supplementary Figs. S28–S34).

Continuous agreement with the 50-run instability estimate was evaluated by directly comparing spot-level instability scores obtained with fewer runs to those obtained with all 50 runs. For each biological sample, spots with finite instability values for all considered run counts (*R* = 5, 10, 20, 40, 50) were identified. To limit the contribution of very large samples and keep the density visualization computationally manageable, at most 10,000 such spots were randomly sampled from each sample using a fixed random seed; all available spots were retained when fewer than 10,000 were available. The selected spots were then pooled across samples within each clustering method. For each *R ∈ {*5, 10, 20, 40*}*, the instability score of every selected spot calculated from *R* runs was plotted against the corresponding instability score calculated from 50 runs. Hexagonal binning was used to show the density of pooled spots in each comparison, and an identity line indicated perfect agreement between the *R*-run and 50-run estimates (Panel C of Fig. 5 and Supplementary Figs. S28–S34).

At the sample level, for each *R∈ {*5, 10, 20, 40 *}*, mean instability was compared with mean ARI across the first *R* stochastic runs using Pearson *r, r*^2^, and the corresponding *p*-value (Panel D of Fig. 5 and Supplementary Figs. S28–S34). At the cluster level, mean cluster instability was compared with Jaccard overlap to the matched reference annotation using the same run counts (Panel E of Fig. 5 and Supplementary Figs. S28–S34). These cluster-level analyses retained the example partition selected from all 50 runs, ensuring that the evaluated clusters themselves remained fixed while the number of runs contributing to the instability estimate changed.

#### Random run-selection convergence analysis

As a complementary assessment of sensitivity to the particular stochastic runs selected, we generated 25 random permutations of the full set of 50 runs. Within each replicate, the *R*-run analysis used the first *R* runs of the corresponding random permutation, so run sets were nested across *R* within a replicate but differed across replicates. For each sample–method combination, run count *R*, and replicate *b* (out of 25), convergence was quantified by

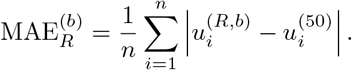

where 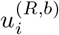 is the instability score for spot *i* calculated from *R ≠* 50 runs in sampling replicate *b* (out of 25) and 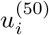 is the instability score calculated from the complete set of 50 runs. The spotwise MAE was then averaged across the 25 replicates for each sample–method combination and run count *R*. Method-level convergence curves summarized these sample-specific values across benchmark samples using the mean and standard error of the mean. Because selecting all 50 available runs yields the same complete set regardless of permutation, all 50-run sampling replicates reproduced the same reference instability vector. Convergence curves are shown in Supplementary Fig. S27.

### Computational Runtime

Computational efficiency was evaluated separately for the clustering procedures and for the instability-score calculation. Clustering runtime was summarized as the mean elapsed time, in seconds, for each method and sample (Supplementary Table S7). For Leiden and Louvain, the reported “Leiden+Louvain” runtime includes the preprocessing step followed by both clustering algorithms. Clustering analyses were performed on the NIH Biowulf cluster. Leiden, Louvain, BayesSpace, and the DLPFC SpiceMix analyses were run without GPU acceleration, whereas the other clustering analyses used NVIDIA Tesla V100 GPUs unless otherwise noted. Jobs were submitted through Slurm using `–exclusive` and `–mem=0` to minimize competition for computational resources. These timings therefore provide a comparison under the computational configurations used in this study rather than optimized benchmarks for each individual method.

The computational cost of the instability calculation itself was evaluated independently using clustering assignments from 50 runs (Supplementary Table S6). For each sample–method combination, the reported runtime is end-to-end wall clock time: reading the 50 partitions, computing instability scores, and writing the final map to disk. Each sample–method calculation was restricted to a single CPU thread, and independent sample–method calculations were run concurrently as separate processes to complete the benchmark efficiently. Benchmarks were performed on the NIH Biowulf cluster on a CPU-only compute node with two AMD EPYC 9454 processors, 96 physical CPU cores (192 hardware threads), and approximately 3 TB of memory. Jobs were submitted through Slurm using `–exclusive` and `–mem=0`.

## Supporting information

Supplemental Results and Methods

## Acknowledgments

W.W. was supported by a National Cancer Institute Cancer Research Training Award, in conjunction with its Partnership for Integrative Cancer Research with the University of Maryland, College Park (seed project award to E.K.M. and J.T.N.). J.T.N. discloses funding by the Surgical Oncology Program, Center for Cancer Research, National Cancer Institute, National Institutes of Health Intramural Research Program project number 1ZIABC012169. E.K.M. discloses funding from the State of Maryland. The contributions of the NIH author(s) are considered Works of the United States Government. The findings and conclusions presented in this paper are those of the author(s) and do not necessarily reflect the views of the State of Maryland, the U.S. National Institutes of Health, or the U.S. Department of Health and Human Services.

## Author Contributions

WW designed and analyzed the instability map algorithm, under the supervision of EKM. WW and EKM designed the computational experiments. WW performed all computational experiments. WW, JN, and EKM interpreted the results. WW and EKM created figures and drafted the manuscript. WW, JN, and EKM edited, reviewed, and approved the manuscript. JN and EKM secured funding for the project. WW conceptualized the project.

