## Supplemental Results and Methods for "Scalable localization of clustering instability for spatial transcriptomics"

### Scalable localization of clustering instability for spatial transcriptomics
#### Supplementary Materials

Wenshan Wu, Joe Nguyen*, Erin K. Molloy*

***

October 8, 2026

### Contents

**List of Tables** 1

**List of Figures** 1

**1 Supplemental Methods** 2

1.1 Algorithm . . . 2

1.2 Dataset Descriptions . . . 4

**2 Supplemental Results** 5

**References** 31

### 1 Supplemental Methods

#### 1.1 Algorithm

**Algorithm 1 Time and storage complexity of instability map with direct cluster-overlap computation.** Clusters are labeled by integers (e.g., int16) $0, 1, \ldots, L^{(r)} - 1$, where $L^{(r)}$ is the total number of clusters in partition $r$. The dictionary is indexed by ordered pairs of cluster labels. We assume dictionary operations (insertions and accesses) have cost $\Theta(1)$.

**Input:** Cluster assignments $C_i^{(r)}$ for spots $i = 1, \ldots, n$ and partitions $r = 1, \ldots, R$

Initialize $N^{(q)}(c) \leftarrow 0$ for all partitions $q$ and respective clusters $c$ ▷ Storage: $\Theta(Rn)$
Initialize $A_i \leftarrow 0$ for all spots $i$ ▷ Storage: $\Theta(n)$
Initialize $\mu_{\text{intra}}^{(r)}(i) \leftarrow 0$ for all partitions $r$ and spots $i$ ▷ Storage: $\Theta(Rn)$
Initialize $u_i \leftarrow 0$ for all spots $i$ ▷ Storage: $\Theta(n)$
**for** $q = 1, \ldots, R$ **do**
    **for** $i = 1, \ldots, n$ **do**
        $N^{(q)}[C_i^{(q)}] \leftarrow N^{(q)}[C_i^{(q)}] + 1$ ▷ Store the size of each cluster in each run
    **end for**
    **for** $i = 1, \ldots, n$ **do**
        $A_i \leftarrow A_i + N^{(q)}[C_i^{(q)}]$ for all $i$ ▷ Store the row sum $\sum_{j=1}^{n} A_{i,j}$
    **end for**
    **for** $r = 1, \ldots, q$ **do**
        $M \leftarrow \{\}$ ▷ Store size of cluster overlaps between partitions $r, q$; Temporary storage: $\Theta(n)$
        **for** $i = 1, \ldots, n$ **do**
            Try: $M[(C_i^{(r)}, C_i^{(q)})] \leftarrow M[(C_i^{(r)}, C_i^{(q)})] + 1$
            Except KeyError: $M[(C_i^{(r)}, C_i^{(q)})] \leftarrow 1$
        **end for**
        **for** $i = 1, \ldots, n$ **do**
            $X_i^{(r,q)} \leftarrow M[(C_i^{(r)}, C_i^{(q)})] - 1$
            $\mu_{\text{intra}}^{(r)}(i) \leftarrow \mu_{\text{intra}}^{(r)}(i) + X_i^{(r,q)}$
            **if** $r \neq q$ **then**
                $\mu_{\text{intra}}^{(q)}(i) \leftarrow \mu_{\text{intra}}^{(q)}(i) + X_i^{(r,q)}$
            **end if**
        **end for**
    **end for**
**end for** ▷ Time: $\Theta(R^2 n)$
**for** $r = 1, \ldots, R$ **do**
    **for** $i = 1, \ldots, n$ **do**
        $|I_i^{(r)}| \leftarrow N^{(r)}[C_i^{(r)}] - 1$
        **if** $|I_i^{(r)}| > 0$ **then**
            $\tilde{\mu}_{\text{intra}}^{(r)}(i) \leftarrow \dfrac{\mu_{\text{intra}}^{(r)}(i)}{|I_i^{(r)}| \cdot R}$
        **else**
            $\tilde{\mu}_{\text{intra}}^{(r)}(i) \leftarrow 1$
        **end if**
        $|O_i^{(r)}| \leftarrow n - N^{(r)}[C_i^{(r)}]$
        **if** $|O_i^{(r)}| > 0$ **then**
            $\mu_{\text{inter}}^{(r)}(i) \leftarrow A_i - \mu_{\text{intra}}^{(r)}(i) - R$
            $\tilde{\mu}_{\text{inter}}^{(r)}(i) \leftarrow \dfrac{\mu_{\text{inter}}^{(r)}(i)}{|O_i^{(r)}| \cdot R}$
        **else**
            $\tilde{\mu}_{\text{inter}}^{(r)}(i) \leftarrow 0$
        **end if**
        $s_i^{(r)} \leftarrow \dfrac{R}{R-1}\left[1 - \left(\tilde{\mu}_{\text{intra}}^{(r)}(i) - \tilde{\mu}_{\text{inter}}^{(r)}(i)\right)\right]$
        $u_i \leftarrow u_i + \dfrac{1}{R} s_i^{(r)}$
    **end for**
**end for** ▷ Time: $\Theta(Rn)$
**return** $\{u_i\}_{i=1}^{n}$
▷ **Overall time complexity:** $\Theta(R^2 n)$, **Peak storage complexity:** $\Theta(Rn)$

#### 1.2 Dataset Descriptions

##### Human dorsolateral prefrontal cortex (DLPFC) dataset

The human dorsolateral prefrontal cortex (DLPFC) dataset was originally generated by Maynard et al. [6] using the 10x Genomics Visium platform. The dataset comprises 12 tissue sections obtained from three independent neurotypical adult donors. For each donor, two pairs of spatial replicates were profiled; each pair consisted of two directly adjacent, 10 $\mu$m-thick serial sections, and the second pair was located approximately 300 $\mu$m posterior to the first. We obtained the processed expression data and reference annotations from the SDMBench resource [11].

Reference annotations correspond to cortical laminar structure and white matter. In the processed annotations used in our analysis, sections 151669–151672 contained five non-missing reference categories, whereas the remaining eight sections contained seven. Following quality control, the 12 sections contained between 2,731 and 4,696 spots and between 13,821 and 15,875 retained genes. A small number of spots in each section lacked reference annotations. These spots were retained during unsupervised clustering and instability estimation and were excluded only from analyses requiring reference labels.

##### Human colorectal cancer (CRC) dataset

We used the publicly available human colorectal cancer (CRC) Visium HD dataset released by 10x Genomics [3]. The sample consists of an FFPE sigmoid-colon tumor section profiled using the 10x Genomics Visium HD platform. We specifically used the dataset processed with Space Ranger v3.0.0 because the pathologist-provided annotations used in this study [5] correspond to this release.

Visium HD provides expression matrices at multiple spatial bin sizes. We used the 16 $\mu$m bin representation to balance spatial resolution, transcript coverage, and computational requirements. After quality control, the full CRC dataset contained 110,402 spatial bins and 17,823 retained genes. The quality-controlled data contained six non-empty reference annotation categories, and this count of six was used as the target reference-domain count in clustering procedures that required one.

We selected a contiguous spatial subset for the primary instability benchmark. Specifically, the quality-controlled tissue was divided horizontally at the midpoint of its spatial $y$-coordinate, the lower half was retained, and a further square region was selected from this half. The square side length was set to 90% of the smaller spatial extent of the retained half. This procedure yielded the final CRC benchmark region containing 21,656 spatial bins. The full quality-controlled CRC dataset was retained separately for computational scalability analyses using Leiden and Louvain.

##### Adult mouse brain (MB) dataset

We used the Adult Mouse Brain FFPE dataset released by 10x Genomics [2], accessed in annotated AnnData format through the `st_datasets` resource [9]. The dataset consists of one adult mouse brain section profiled using the 10x Genomics Visium FFPE workflow. The original 10x dataset contains 2,264 spots under tissue and has a reported median of 17,614 UMI counts per spot.

We used the 10-category reference partition distributed with the annotated dataset for evaluation. Following quality control, all 2,264 spatial spots were retained, with expression measurements for 14,022 genes.

##### Human breast cancer (HBC) dataset

We used the Human Breast Cancer (HBC) Block A, Section 1 dataset released by 10x Genomics [1], accessed in annotated AnnData format through the `st_datasets` resource [9]. The sample consists of fresh-frozen invasive ductal carcinoma tissue profiled using the 10x Genomics Visium platform and processed using Space Ranger v1.1.0.

For reference evaluation, we used the detailed histopathological annotations described by Xu et al. [10]. The tissue was manually segmented into 20 regions based on pathological features, which can be grouped into four broader morphotypes: ductal carcinoma in situ/lobular carcinoma in situ (DCIS/LCIS), invasive ductal carcinoma (IDC), healthy tissue, and tumor-edge regions. The 20-region annotation was retained as the reference partition in our analysis. Following quality control, the dataset contained 3,795 spatial spots and 18,233 retained genes.

##### Mouse cerebellum (MBC) dataset

We used a mouse cerebellum dataset generated using Slide-seq [8], accessed in annotated AnnData format through the `st_datasets` resource [9]. The original Slide-seq experiment did not provide a manually defined spatial-domain ground truth. We therefore used the eight-category reference partition supplied with the benchmark dataset distribution, which was derived for spatial-domain benchmarking and represents major anatomical structures of the cerebellum [4]. Following quality control, the dataset contained 25,548 spatial locations and 11,970 retained genes. These eight benchmark-derived spatial labels were used as the reference partition for analyses requiring an external annotation.

##### SPATCH Visium HD tumor (COAD and OV) datasets

We analyzed colon adenocarcinoma (COAD) and ovarian cancer (OV) samples from the SPATCH spatial-omics resource [7]. These samples were derived from treatment-naive human tumors and profiled from FFPE tissue using the 10x Genomics Visium HD platform. We analyzed expression aggregated at 8 $\mu$m spatial-bin resolution. Histology images and corresponding spatial coordinates were incorporated into the AnnData objects for visualization; histology images were not used as input to the clustering methods evaluated in this study. Following quality control, the full COAD and OV datasets contained 503,622 and 410,006 bins, respectively, with 17,757 retained genes in each.

Contiguous spatial subsets were used for the primary multi-method benchmark while the complete quality-controlled tissues were retained for scalability analyses. For each sample, the quality-controlled tissue was divided into a regular $4 \times 4$ grid according to its spatial-coordinate range, and one contiguous grid region was selected for downstream analysis. The selected region was then restricted further to a square crop. The square side length was set to 91.5% of the smaller spatial extent of the selected COAD grid region and 90% of that of the selected OV grid region. The resulting benchmark regions contained 27,494 bins for COAD and 27,291 bins for OV. The cropped COAD region contained 15 reference categories, whereas the cropped OV region contained 13.

### 2 Supplemental Results

Table S1: Cluster instability by method.

| Method | mean frac. unstable spots | total # clusters | fraction of clusters (s/ms/u/hu) | Purity R | Recall R | Jaccard R | Exp. Coherence R | Exp. Variance R |
|---|---|---|---|---|---|---|---|---|
| SEDR | 0.31 ± 0.16 | 132 | 0.32/0.42/0.26/0.01 | -0.41 (0.17) | -0.55 (0.31) | -0.72 (0.52) | -0.55 (0.30) | 0.55 (0.30) |
| GraphST | 0.35 ± 0.20 | 148 | 0.32/0.38/0.29/0.01 | -0.46 (0.21) | -0.41 (0.17) | -0.56 (0.31) | -0.56 (0.32) | 0.56 (0.32) |
| STAGATE | 0.27 ± 0.11 | 127 | 0.32/0.46/0.20/0.02 | -0.43 (0.19) | **-0.37 (0.13)** | **-0.47 (0.22)** | -0.31 (0.10) | 0.31 (0.10) |
| BayesSpace | 0.09 ± 0.09 | 148 | 0.69/0.25/0.06/0.00 | **-0.19 (0.04)** | **-0.34 (0.12)** | **-0.40 (0.16)** | -0.26 (0.07) | 0.26 (0.07) |
| Louvain | 0.25 ± 0.13 | 138 | 0.45/0.36/0.20/0.00 | -0.48 (0.23) | -0.46 (0.21) | -0.57 (0.33) | -0.67 (0.45) | 0.67 (0.44) |
| Leiden | 0.24 ± 0.13 | 146 | 0.45/0.41/0.14/0.01 | -0.46 (0.21) | -0.55 (0.30) | -0.61 (0.38) | -0.65 (0.42) | 0.65 (0.42) |
| SpiceMix | 0.34 ± 0.19 | 114 | 0.33/0.31/0.33/0.03 | **-0.30 (0.09)** | -0.50 (0.25) | -0.58 (0.34) | -0.57 (0.33) | 0.57 (0.32) |
| **SEDR with different backend clustering algorithms (DLPFC samples only)** | | | | | | | | |
| Leiden | 0.35 ± 0.12 | 68 | 0.25/0.44/0.31/0.00 | -0.46 (0.21) | -0.65 (0.42) | -0.75 (0.56) | -0.58 (0.33) | 0.58 (0.33) |
| mclust | 0.22 ± 0.08 | 76 | 0.25/0.51/0.24/0.00 | -0.29 (0.09) | -0.47 (0.22) | -0.55 (0.30) | -0.22 (0.05) | 0.22 (0.05) |

*Note.* Rows correspond to different clustering methods. Second column is the fraction of unstable spots (with scores greater than or equal to 0.4) in each analysis (sample-method pair) averaged across all samples keeping the method fixed (standard deviation also shown). Third column is the total number of clusters produced from running the method on up to 18 samples. Fourth column is the fraction of those clusters in the four categories: $s$ = stable, $ms$ = modestly stable, $u$ = unstable, $hu$ = highly unstable. The remaining columns show the Pearson $r$ ($r^2$) correlation between cluster instability (mean score across spots) versus another cluster-level metric, e.g., cluster purity with respect to the reference annotation (all clusters produced by the method, combining across the different samples, are used in the correlation analysis).

Table S2: Cluster instability by sample.

| Sample | mean frac. unstable spots | # clusters | fraction of clusters (s/ms/u/hu) |
|---|---|---|---|
| MBC | 0.03 ± 0.04 | 48 | 0.88/0.10/0.02/0.00 |
| CRC | 0.10 ± 0.07 | 38 | 0.71/0.26/0.03/0.00 |
| HBC | 0.13 ± 0.09 | 121 | 0.55/0.34/0.08/0.02 |
| MB | 0.09 ± 0.09 | 65 | 0.69/0.26/0.05/0.00 |
| COAD | 0.25 ± 0.10 | 92 | 0.34/0.46/0.21/0.00 |
| OV | 0.35 ± 0.20 | 80 | 0.36/0.41/0.20/0.03 |
| **DLPFC** | | | |
| 151507 | 0.38 ± 0.15 | 53 | 0.28/0.30/0.38/0.04 |
| 151508 | 0.32 ± 0.21 | 55 | 0.38/0.35/0.27/0.00 |
| 151509 | 0.30 ± 0.15 | 50 | 0.32/0.44/0.24/0.00 |
| 151510 | 0.38 ± 0.14 | 53 | 0.30/0.26/0.43/0.00 |
| 151669 | 0.27 ± 0.12 | 40 | 0.15/0.62/0.20/0.03 |
| 151670 | 0.23 ± 0.12 | 43 | 0.35/0.53/0.12/0.00 |
| 151671 | 0.22 ± 0.12 | 40 | 0.45/0.40/0.15/0.00 |
| 151672 | 0.20 ± 0.10 | 41 | 0.39/0.51/0.10/0.00 |
| 151673 | 0.27 ± 0.15 | 52 | 0.37/0.44/0.19/0.00 |
| 151674 | 0.30 ± 0.13 | 54 | 0.22/0.43/0.35/0.00 |
| 151675 | 0.37 ± 0.12 | 52 | 0.23/0.37/0.38/0.02 |
| 151676 | 0.40 ± 0.14 | 52 | 0.19/0.38/0.42/0.00 |

*Note.* Rows correspond to different benchmark samples. Second column is the fraction of unstable spots (with scores greater than or equal to 0.4) in each analysis (sample-method pair) averaged across all methods keeping the sample fixed (standard deviation also shown). Third column is the total number of clusters produced from running up to 8 configurations on the sample. Fourth column is the fraction of those clusters in the four categories: $s$ = stable, $ms$ = modestly stable, $u$ = unstable, $hu$ = highly unstable.

Table S3: Mean ARI across 50 runs for each sample–method pair.

| Sample | SEDR | GraphST | STAGATE | Bayes Space | Louvain | Leiden | SpiceMix | SEDR mclust | Mean ± S.D. |
|---|---|---|---|---|---|---|---|---|---|
| MBC | 0.99 | 0.90 | – | 0.61 | 0.99 | **1.00** | 0.70 | – | 0.86 ± 0.17 |
| CRC | 0.65 | 0.64 | **0.67** | 0.49 | 0.56 | 0.59 | 0.52 | – | 0.59 ± 0.07 |
| HBC | 0.51 | **0.53** | 0.48 | 0.51 | 0.47 | 0.45 | 0.41 | – | 0.48 ± 0.04 |
| MB | 0.40 | 0.43 | 0.43 | 0.41 | 0.49 | **0.50** | 0.42 | – | 0.44 ± 0.04 |
| COAD | 0.16 | 0.19 | 0.07 | 0.19 | 0.17 | 0.17 | **0.26** | – | 0.17 ± 0.05 |
| OV | 0.12 | 0.15 | 0.04 | 0.13 | 0.12 | 0.13 | **0.20** | – | 0.13 ± 0.05 |
| **DLPFC** | | | | | | | | | |
| 151507 | 0.43 | 0.42 | 0.44 | 0.40 | 0.32 | 0.32 | 0.36 | **0.46** | 0.39 ± 0.05 |
| 151508 | 0.39 | 0.35 | 0.41 | 0.34 | 0.27 | 0.27 | 0.32 | **0.46** | 0.35 ± 0.06 |
| 151509 | 0.38 | 0.36 | 0.37 | 0.35 | 0.34 | 0.34 | 0.37 | **0.42** | 0.37 ± 0.03 |
| 151510 | 0.41 | 0.34 | 0.37 | 0.39 | 0.31 | 0.31 | 0.34 | **0.43** | 0.36 ± 0.05 |
| 151669 | 0.38 | 0.36 | 0.36 | 0.43 | 0.27 | 0.33 | 0.28 | **0.47** | 0.36 ± 0.07 |
| 151670 | 0.38 | 0.41 | 0.32 | **0.44** | 0.24 | 0.30 | 0.29 | 0.38 | 0.34 ± 0.07 |
| 151671 | 0.52 | 0.55 | 0.56 | 0.53 | 0.38 | 0.38 | 0.42 | **0.58** | 0.49 ± 0.08 |
| 151672 | 0.51 | 0.53 | 0.52 | **0.61** | 0.39 | 0.38 | 0.39 | 0.57 | 0.49 ± 0.09 |
| 151673 | 0.48 | 0.43 | 0.42 | 0.51 | 0.36 | 0.36 | 0.39 | **0.52** | 0.44 ± 0.06 |
| 151674 | 0.46 | 0.43 | 0.48 | 0.34 | 0.36 | 0.33 | 0.37 | **0.55** | 0.42 ± 0.08 |
| 151675 | 0.50 | 0.41 | 0.43 | 0.36 | 0.39 | 0.37 | 0.39 | **0.56** | 0.42 ± 0.07 |
| 151676 | 0.39 | 0.35 | 0.41 | 0.34 | 0.34 | 0.33 | 0.38 | **0.48** | 0.38 ± 0.05 |
| **Mean ± S.D.** | | | | | | | | | |
| All | 0.45 ± 0.18 | 0.43 ± 0.17 | 0.40 ± 0.15 | 0.41 ± 0.13 | 0.38 ± 0.19 | 0.38 ± 0.19 | 0.38 ± 0.11 | – | |
| DLPFC only | 0.44 ± 0.06 | 0.41 ± 0.07 | 0.42 ± 0.07 | 0.42 ± 0.09 | 0.33 ± 0.05 | 0.33 ± 0.03 | 0.36 ± 0.04 | 0.49 ± 0.06 | |

Table S4: Mean instability (i.e., mean instability score across all spots) for each sample–method pair.

| Sample | SEDR | GraphST | STAGATE | Bayes Space | Louvain | Leiden | SpiceMix | SEDR mclust | Mean ± S.D. |
|---|---|---|---|---|---|---|---|---|---|
| MBC | 0.02 | 0.06 | – | 0.19 | 0.00 | 0.00 | 0.12 | – | 0.06 ± 0.08 |
| CRC | 0.14 | 0.14 | 0.16 | 0.13 | 0.17 | 0.18 | 0.23 | – | 0.17 ± 0.03 |
| HBC | 0.24 | 0.19 | 0.32 | 0.15 | 0.18 | 0.16 | 0.04 | – | 0.18 ± 0.09 |
| MB | 0.29 | 0.11 | 0.27 | 0.06 | 0.11 | 0.09 | 0.07 | – | 0.15 ± 0.09 |
| COAD | 0.35 | 0.28 | 0.24 | 0.17 | 0.26 | 0.25 | 0.21 | – | 0.25 ± 0.06 |
| OV | 0.41 | 0.35 | 0.31 | 0.20 | 0.21 | 0.25 | 0.50 | – | 0.32 ± 0.11 |
| **DLPFC** | | | | | | | | | |
| 151507 | 0.34 | 0.44 | 0.28 | 0.30 | 0.32 | 0.37 | 0.37 | 0.26 | 0.33 ± 0.06 |
| 151508 | 0.37 | 0.41 | 0.24 | 0.21 | 0.24 | 0.31 | 0.43 | 0.17 | 0.30 ± 0.10 |
| 151509 | 0.37 | 0.41 | 0.26 | 0.15 | 0.29 | 0.25 | 0.33 | 0.34 | 0.30 ± 0.08 |
| 151510 | 0.39 | 0.46 | 0.34 | 0.23 | 0.33 | 0.29 | 0.37 | 0.32 | 0.34 ± 0.07 |
| 151669 | 0.30 | 0.42 | 0.27 | 0.12 | 0.37 | 0.34 | 0.33 | 0.32 | 0.31 ± 0.09 |
| 151670 | 0.27 | 0.27 | 0.31 | 0.03 | 0.36 | 0.31 | 0.29 | 0.26 | 0.26 ± 0.10 |
| 151671 | 0.25 | 0.31 | 0.21 | 0.07 | 0.36 | 0.27 | 0.33 | 0.19 | 0.25 ± 0.09 |
| 151672 | 0.28 | 0.23 | 0.24 | 0.08 | 0.26 | 0.24 | 0.36 | 0.23 | 0.24 ± 0.08 |
| 151673 | 0.35 | 0.41 | 0.32 | 0.15 | 0.19 | 0.24 | 0.35 | 0.23 | 0.28 ± 0.09 |
| 151674 | 0.31 | 0.35 | 0.24 | 0.18 | 0.32 | 0.28 | 0.36 | 0.33 | 0.30 ± 0.06 |
| 151675 | 0.34 | 0.38 | 0.36 | 0.25 | 0.29 | 0.33 | 0.44 | 0.25 | 0.33 ± 0.07 |
| 151676 | 0.39 | 0.46 | 0.37 | 0.32 | 0.24 | 0.33 | 0.28 | 0.35 | 0.34 ± 0.07 |
| **Mean ± S.D.** | | | | | | | | | |
| All | 0.30 ± 0.10 | 0.32 ± 0.12 | 0.28 ± 0.05 | 0.17 ± 0.08 | 0.25 ± 0.10 | 0.25 ± 0.09 | 0.30 ± 0.12 | – | |
| DLPFC only | 0.33 ± 0.05 | 0.38 ± 0.07 | 0.29 ± 0.05 | 0.17 ± 0.09 | 0.30 ± 0.06 | 0.30 ± 0.04 | 0.35 ± 0.05 | 0.27 ± 0.06 | |

Table S5: **BayesSpace Entropy and Instability.** Mean instability of BayesSpace is the instability score averaged across all spots in the sample. Mean entropy is the entropy averaged across all spots in the sample, then averaged across all 50 BayesSpace runs.

| Sample | Mean Instability | Mean Entropy |
|---|---|---|
| MBC | 0.19 | 0.10 |
| CRC | 0.13 | 0.06 |
| HBC | 0.15 | 0.14 |
| MB | 0.06 | 0.05 |
| COAD | 0.17 | 0.21 |
| OV | 0.20 | 0.42 |
| **DLPFC** | | |
| 151507 | 0.30 | 0.15 |
| 151508 | 0.21 | 0.21 |
| 151509 | 0.15 | 0.19 |
| 151510 | 0.23 | 0.20 |
| 151669 | 0.12 | 0.14 |
| 151670 | 0.03 | 0.15 |
| 151671 | 0.07 | 0.13 |
| 151672 | 0.08 | 0.13 |
| 151673 | 0.15 | 0.14 |
| 151674 | 0.18 | 0.12 |
| 151675 | 0.25 | 0.21 |
| 151676 | 0.32 | 0.16 |

Table S6: **Runtime of instability score calculation from 50 runs.** Runtimes are given in seconds and correspond to the single-thread computational time for each individual sample–method combination. NA indicates that the method was not run on the sample. Runtimes include loading and preprocessing of the saved clustering assignments, instability-score computation, and saving of the resulting instability scores to disk. All benchmarks were performed on the NIH Biowulf cluster on a CPU-only compute node with two AMD EPYC 9454 processors, 96 physical CPU cores (192 hardware threads), and approximately 3 TB of memory. Independent sample–method calculations were run concurrently as separate single-threaded processes. The Slurm job used `--exclusive` and `--mem=0` so that the allocated node was not shared with other Slurm jobs.

| Dataset | # Spots | Leiden | Louvain | Bayes-Space | GraphST | STAGATE | SpiceMix | SEDR | SEDR-mclust |
|---|---|---|---|---|---|---|---|---|---|
| **DLPFC** | | | | | | | | | |
| 151507 | 4,020 | 0.06 | 0.04 | 0.08 | 0.05 | 0.06 | 0.07 | 0.05 | 0.06 |
| 151508 | 4,001 | 0.07 | 0.06 | 0.07 | 0.06 | 0.07 | 0.06 | 0.07 | 0.07 |
| 151509 | 4,696 | 0.06 | 0.07 | 0.09 | 0.06 | 0.06 | 0.06 | 0.07 | 0.09 |
| 151510 | 4,552 | 0.07 | 0.06 | 0.08 | 0.08 | 0.08 | 0.07 | 0.05 | 0.08 |
| 151669 | 3,022 | 0.06 | 0.07 | 0.10 | 0.07 | 0.07 | 0.08 | 0.06 | 0.09 |
| 151670 | 2,731 | 0.05 | 0.06 | 0.07 | 0.04 | 0.07 | 0.04 | 0.07 | 0.05 |
| 151671 | 3,461 | 0.08 | 0.06 | 0.10 | 0.05 | 0.08 | 0.08 | 0.05 | 0.06 |
| 151672 | 3,402 | 0.05 | 0.04 | 0.08 | 0.07 | 0.07 | 0.07 | 0.07 | 0.08 |
| 151673 | 3,534 | 0.08 | 0.07 | 0.07 | 0.05 | 0.06 | 0.05 | 0.05 | 0.08 |
| 151674 | 3,649 | 0.04 | 0.07 | 0.08 | 0.07 | 0.04 | 0.05 | 0.07 | 0.09 |
| 151675 | 3,173 | 0.07 | 0.07 | 0.08 | 0.07 | 0.06 | 0.29 | 0.04 | 0.04 |
| 151676 | 3,149 | 0.05 | 0.06 | 0.08 | 0.08 | 0.07 | 0.04 | 0.04 | 0.04 |
| MB | 2,264 | 0.04 | 0.04 | 0.08 | 0.03 | 0.03 | 0.07 | 0.03 | NA |
| HBC | 3,795 | 0.04 | 0.04 | 0.07 | 0.07 | 0.07 | 0.07 | 0.07 | NA |
| MBC | 25,548 | 0.18 | 0.18 | 0.22 | 0.22 | NA | 0.19 | 0.19 | NA |
| **Cropped Visium HD Samples** | | | | | | | | | |
| CRC | 21,656 | 0.17 | 0.17 | 0.25 | 0.17 | 0.16 | 0.17 | 0.40 | NA |
| COAD | 27,494 | 0.19 | 0.25 | 0.26 | 0.30 | 0.23 | 0.29 | 0.19 | NA |
| OV | 27,291 | 0.20 | 0.20 | 0.23 | 0.19 | 0.23 | 0.19 | 0.20 | NA |
| **Full Visium HD Samples** | | | | | | | | | |
| CRC | 110,402 | 0.81 | 0.78 | NA | NA | NA | NA | NA | NA |
| OV | 410,006 | 2.74 | 2.80 | NA | NA | NA | NA | NA | NA |
| COAD | 503,622 | 3.35 | 3.34 | NA | NA | NA | NA | NA | NA |

Table S7: **Mean runtime for clustering methods across 50 runs.** Runtimes are given in seconds. The "Leiden+Louvain" runtime includes the time for preprocessing data and then running both of the two clustering algorithms, as implemented in scanpy. NA indicates the method was not run on the sample, with the exception that STAGATE had a memory error on the MBC sample. All clustering analyses were performed on the NIH Biowulf cluster. All other methods were run on compute nodes with GPUs (NVIDIA Tesla V100 GPUs), unless otherwise noted. Jobs were submitted via Slurm with flags `--exclusive`, and `--mem=0` to ensure there were not competing processes on the node. Better timings could be achieved through more careful optimization of the compute environment for each clustering method.

| Dataset | Spots | Leiden+ Louvain (no gpu) | BayesSpace (no gpu) | GraphST | STAGATE | SpiceMix | SEDR |
|---|---|---|---|---|---|---|---|
| **DLPFC** | | | | | | | |
| 151507 | 4,020 | 4.24 | 6.46 | 40.18 | 136.87 | 1760.70 (no gpu) | 40.73 |
| 151508 | 4,001 | 4.05 | 6.24 | 42.92 | 137.06 | 2138.54 (no gpu) | 41.45 |
| 151509 | 4,696 | 4.44 | 7.20 | 54.65 | 151.16 | 2542.45 (no gpu) | 50.69 |
| 151510 | 4,552 | 4.18 | 7.05 | 49.42 | 139.80 | 2115.86 (no gpu) | 46.66 |
| 151669 | 3,022 | 3.57 | 5.22 | 35.20 | 93.14 | 1557.78 (no gpu) | 37.73 |
| 151670 | 2,731 | 3.16 | 4.76 | 32.66 | 92.31 | 1339.13 (no gpu) | 30.04 |
| 151671 | 3,461 | 4.06 | 5.73 | 36.75 | 130.07 | 1816.93 (no gpu) | 36.22 |
| 151672 | 3,402 | 3.98 | 5.63 | 34.14 | 106.48 | 1610.31 (no gpu) | 34.56 |
| 151673 | 3,534 | 3.33 | 5.92 | 36.24 | 112.34 | 2016.46 (no gpu) | 34.91 |
| 151674 | 3,649 | 3.58 | 6.14 | 38.31 | 123.28 | 2322.81 (no gpu) | 36.31 |
| 151675 | 3,173 | 4.05 | 5.40 | 32.73 | 97.93 | 1857.59 (no gpu) | 30.99 |
| 151676 | 3,149 | 3.22 | 5.35 | 32.59 | 113.40 | 1869.53 (no gpu) | 31.06 |
| MB | 2,264 | 2.78 | 9.17 | 43.79 | 84.73 | 1118.78 (no gpu) | 26.17 |
| HBC | 3,795 | 4.39 | 12.03 | 41.19 | 119.90 | 1605.47 (no gpu) | 38.75 |
| MBC | 25,548 | 11.50 | 39.55 | 601.18 | failed | 4052.87 | 261.62 |
| **Cropped Visium HD Samples** | | | | | | | |
| CRC | 21,656 | 42.53 | 39.26 | 453.85 | 338.16 | 1960.31 | 404.26 |
| COAD | 27,494 | 83.97 | 79.07 | 845.09 | 413.64 | 2217.63 | 504.87 |
| OV | 27,291 | 94.77 | 77.68 | 882.90 | 1076.41 | 4572.86 | 561.75 |
| **Full Visium HD Samples** | | | | | | | |
| CRC | 110,402 | 421.94 | NA | NA | NA | NA | NA |
| OV | 410,006 | 1766.05 | NA | NA | NA | NA | NA |
| COAD | 503,622 | 2387.68 | NA | NA | NA | NA | NA |

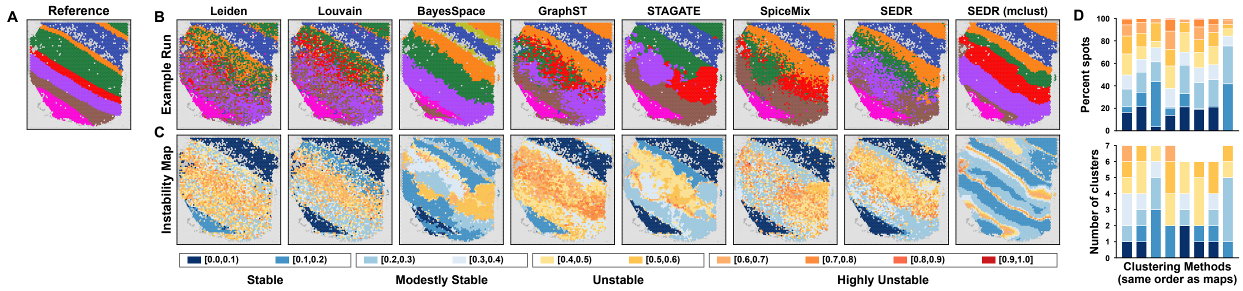

Figure S1: **Spatially resolved clustering instability for DLPFC sample 151507. A)** Reference annotation distributed with the benchmark. **B)** The example run, defined as the first clustering run (out of 50) with the fewest clusters, is shown for all methods. Accuracy (ARI) of example run with different clustering methods going from left to right is 0.32, 0.32, 0.43, 0.41, 0.43, 0.40, 0.42, and 0.47. **C)** Spot-level clustering instability maps estimated from 50 stochastic runs for each method. Mean instability for clustering methods going from left to right is 0.37, 0.32, 0.30, 0.44, 0.28, 0.37, 0.34, and 0.26. **D)** Stacked bars (one per method in the same order as the maps) show the fraction of spots assigned to each instability interval (top) and the number of clusters assigned to each instability interval based on the mean score (bottom).

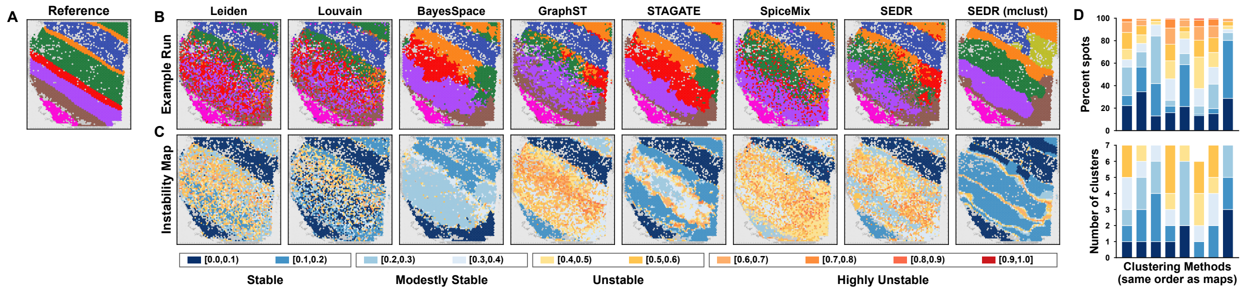

Figure S2: **Spatially resolved clustering instability for DLPFC Visium sample 151508.** Accuracy (ARI) of methods going from left to right is 0.28, 0.28, 0.35, 0.33, 0.41, 0.37, 0.40, and 0.43. Mean instability going from left to right is 0.31, 0.24, 0.21, 0.41, 0.24, 0.43, 0.37, and 0.17.

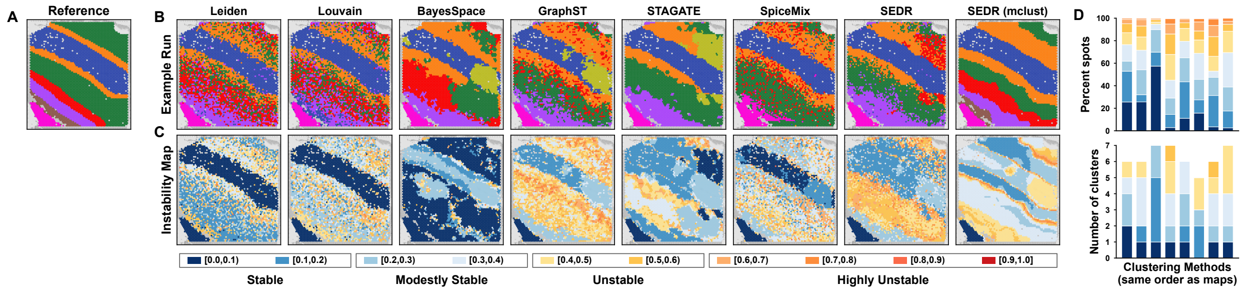

Figure S3: **Spatially resolved clustering instability for DLPFC Visium sample 151509.** The accuracy (ARI) for clustering methods going from left to right is 0.36, 0.34, 0.35, 0.34, 0.50, 0.37, 0.46, and 0.53. Mean instability going from left to right is 0.25, 0.29, 0.15, 0.41, 0.26, 0.33, 0.37, and 0.34.

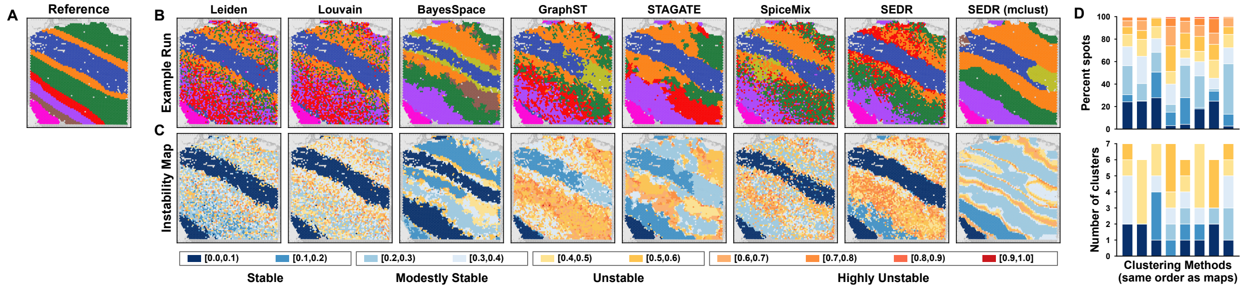

Figure S4: **Spatially resolved clustering instability for DLPFC Visium sample 151510.** Accuracy (ARI) for clustering methods going from left to right is 0.31, 0.32, 0.38, 0.31, 0.40, 0.33, 0.45, and 0.38. Mean instability going from left to right is 0.29, 0.33, 0.23, 0.46, 0.34, 0.37, 0.39, and 0.32.

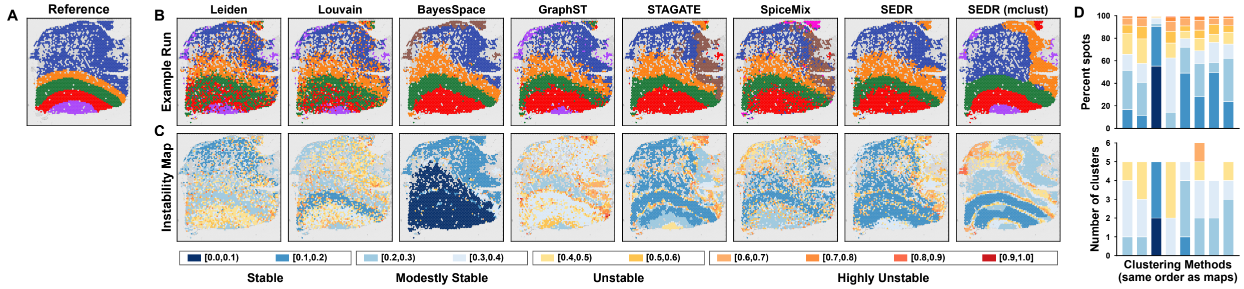

Figure S5: **Spatially resolved clustering instability for DLPFC Visium sample 151669.** Accuracy (ARI) for clustering methods going from left to right is 0.34, 0.33, 0.44, 0.45, 0.37, 0.32, 0.51, and 0.44. Mean instability going from left to right is 0.34, 0.37, 0.12, 0.42, 0.27, 0.33, 0.30, and 0.32.

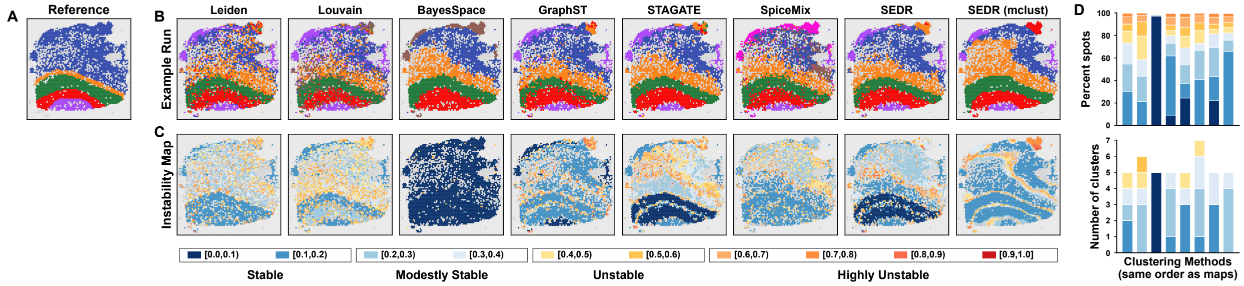

Figure S6: **Spatially resolved clustering instability for DLPFC Visium sample 151670.** Accuracy (ARI) for clustering methods going from left to right is 0.31, 0.26, 0.44, 0.41, 0.38, 0.29, 0.43, and 0.37. Mean instability going from left to right is 0.31, 0.36, 0.03, 0.27, 0.31, 0.29, 0.27, and 0.26.

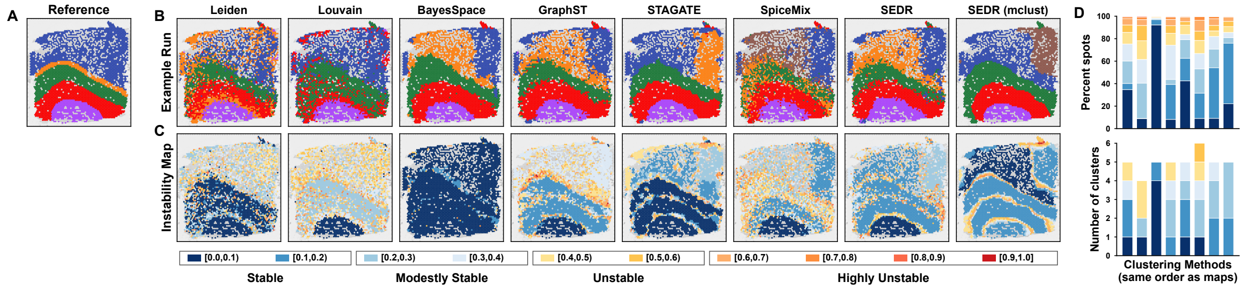

Figure S7: **Spatially resolved clustering instability for DLPFC Visium sample 151671.** Accuracy (ARI) for clustering methods going from left to right is 0.40, 0.47, 0.52, 0.55, 0.60, 0.47, 0.53, and 0.56. Mean instability going from left to right is 0.27, 0.36, 0.07, 0.31, 0.21, 0.33, 0.25, and 0.19.

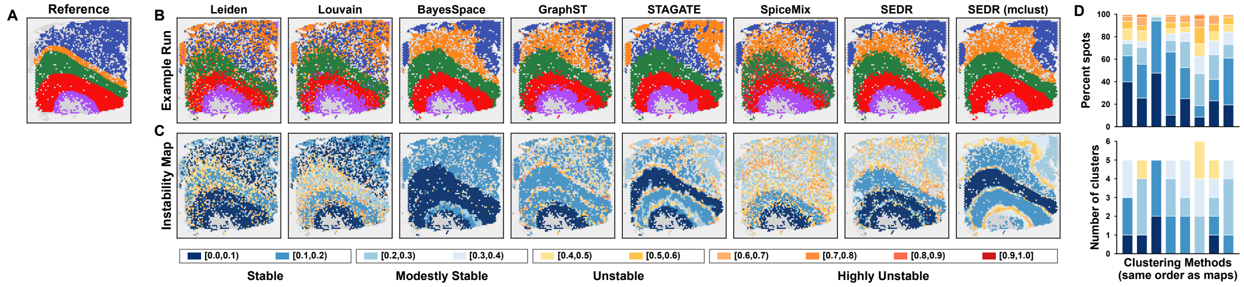

Figure S8: **Spatially resolved clustering instability for DLPFC Visium sample 151672.** Accuracy (ARI) for clustering methods going from left to right is 0.39, 0.39, 0.59, 0.54, 0.56, 0.42, 0.56, and 0.58. Mean instability going from left to right is 0.24, 0.26, 0.08, 0.23, 0.24, 0.36, 0.28, and 0.23.

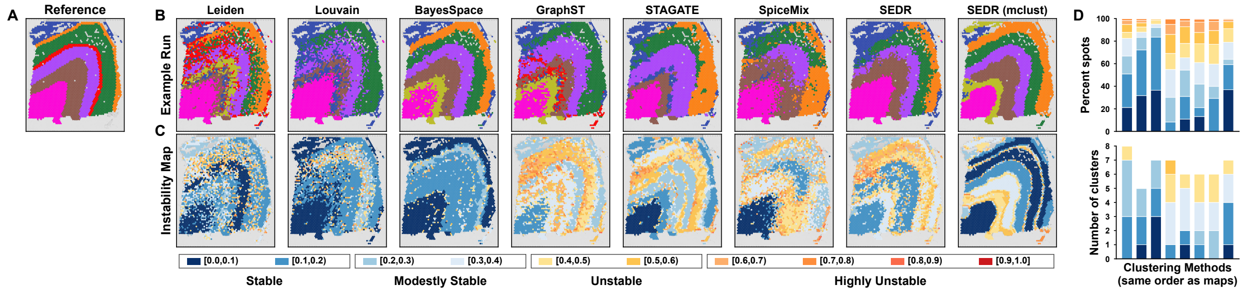

Figure S9: **Spatially resolved clustering instability for DLPFC Visium sample 151673.** Accuracy (ARI) for clustering methods going from left to right is 0.38, 0.37, 0.53, 0.42, 0.44, 0.41, 0.48, and 0.53. Mean instability going from left to right is 0.24, 0.19, 0.15, 0.41, 0.32, 0.35, 0.35, and 0.23.

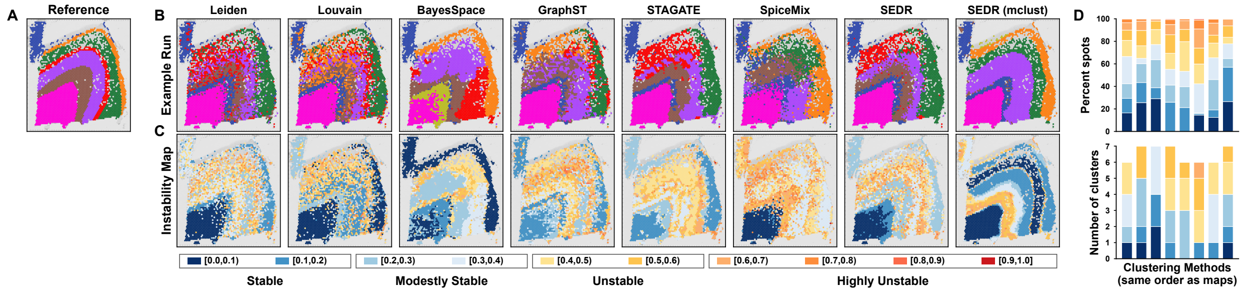

Figure S10: **Spatially resolved clustering instability for DLPFC Visium sample 151675.** Accuracy (ARI) for clustering methods going from left to right is 0.39, 0.41, 0.37, 0.43, 0.50, 0.36, 0.52, and 0.52. Mean instability going from left to right is 0.33, 0.29, 0.25, 0.38, 0.36, 0.44, 0.34, and 0.25.

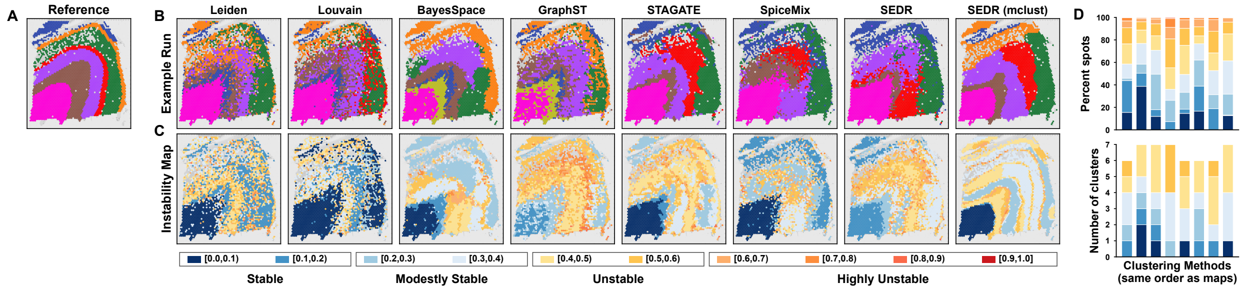

Figure S11: **Spatially resolved clustering instability for DLPFC Visium sample 151676.** Accuracy (ARI) for clustering methods going from left to right is 0.37, 0.35, 0.34, 0.31, 0.41, 0.36, 0.38, and 0.42. Mean instability going from left to right is 0.33, 0.24, 0.32, 0.46, 0.37, 0.28, 0.39, and 0.35.

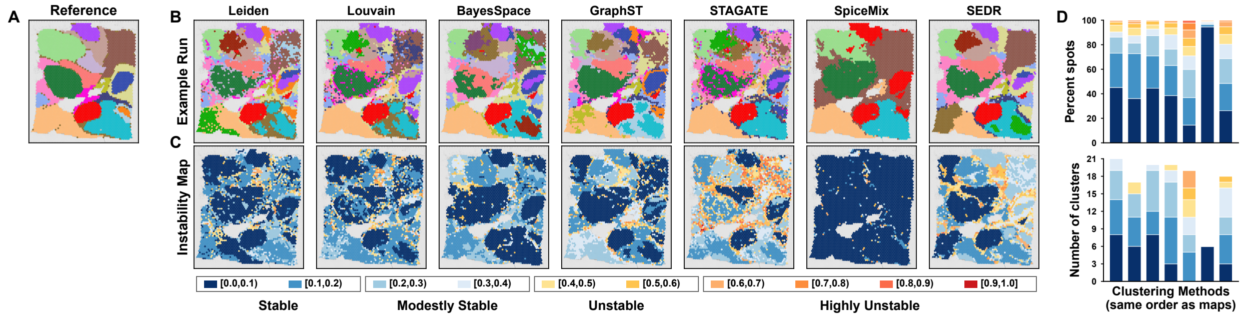

Figure S12: **Spatially resolved clustering instability for HBC Visium sample.** Accuracy (ARI) for clustering methods going from left to right is 0.47, 0.50, 0.51, 0.51, 0.53, 0.39, and 0.54. Mean instability going from left to right is 0.16, 0.18, 0.15, 0.19, 0.32, 0.04, and 0.24.

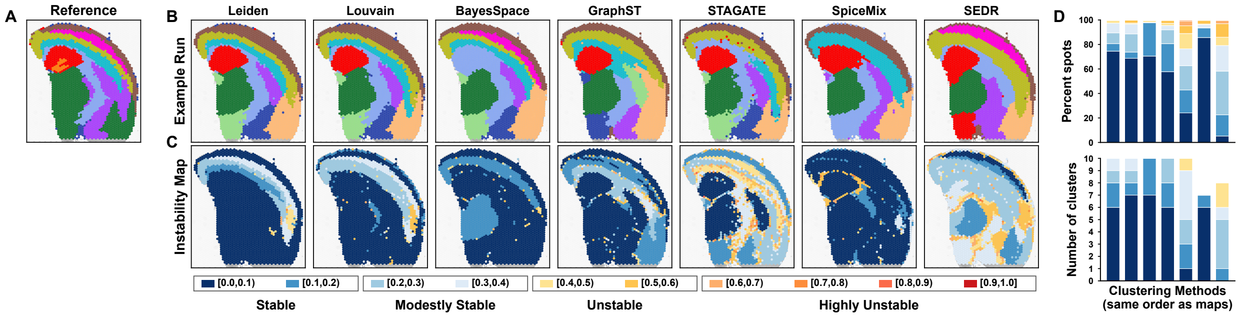

Figure S13: **Spatially resolved clustering instability for MB Visium sample.** Accuracy (ARI) for clustering methods going from left to right is 0.51, 0.50, 0.42, 0.41, 0.43, 0.41, and 0.38. Mean instability going from left to right is 0.09, 0.11, 0.06, 0.11, 0.27, 0.07, and 0.29.

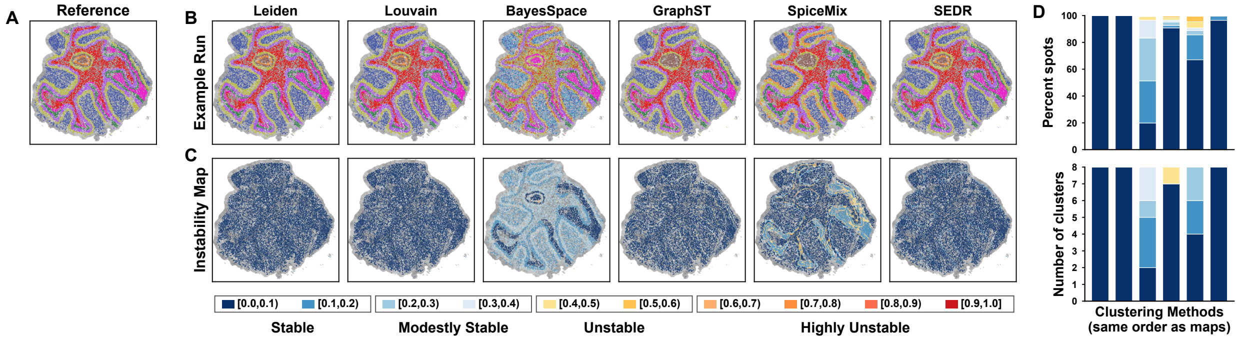

Figure S14: **Spatially resolved clustering instability for MBC Slide-seq sample.** Accuracy (ARI) for clustering methods going from left to right is 1.00, 0.99, 0.62, 0.91, NA, 0.67, and 0.99. Mean instability going from left to right is 0.00, 0.00, 0.19, 0.06, NA, 0.12, and 0.02. STAGATE failed to run on this sample due to a memory error.

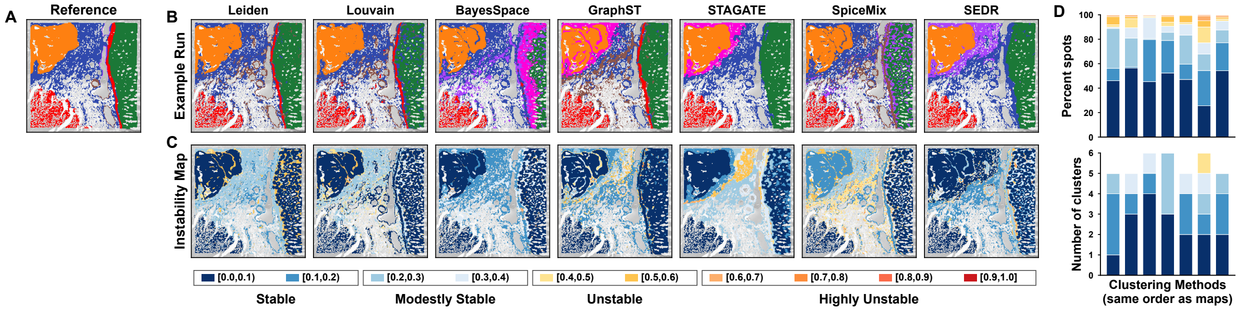

Figure S15: **Spatially resolved clustering instability for CRC Visium HD sample.** Accuracy (ARI) for clustering methods going from left to right is 0.72, 0.60, 0.49, 0.62, 0.71, 0.55, and 0.67. Mean instability going from left to right is 0.18, 0.17, 0.13, 0.14, 0.16, 0.23, and 0.14. This sample was cropped so that all methods could be run.

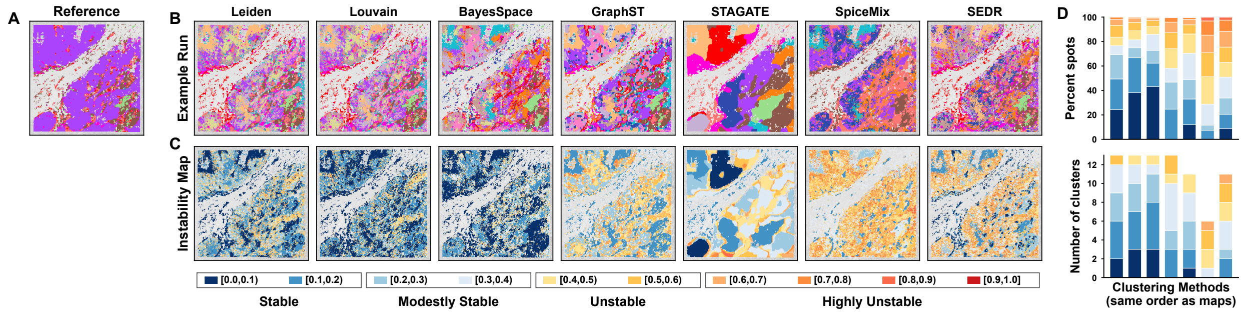

Figure S16: **Spatially resolved clustering instability for OV Visium HD sample.** Accuracy (ARI) for clustering methods going from left to right is 0.13, 0.13, 0.13, 0.15, 0.06, 0.14, and 0.13. Mean instability going from left to right is 0.25, 0.20, 0.20, 0.35, 0.31, 0.50, and 0.41. This sample was cropped so that all methods could be run.

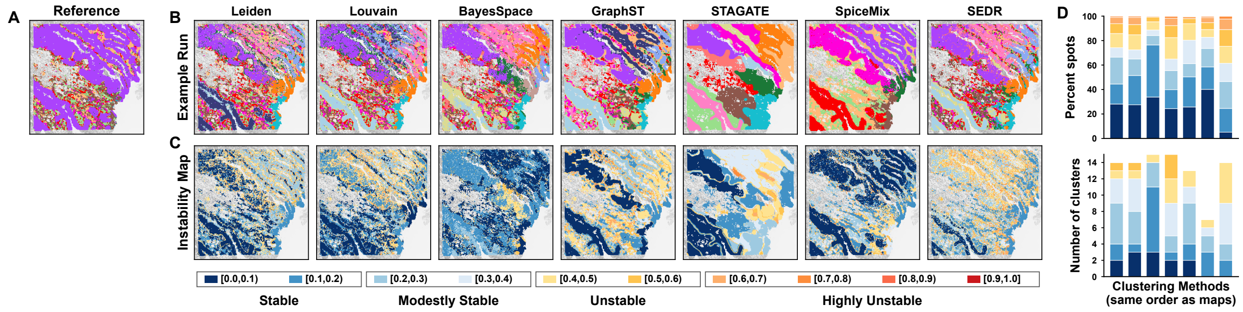

Figure S17: **Spatially resolved clustering instability for COAD Visium HD sample.** Accuracy (ARI) for clustering methods going from left to right is 0.18, 0.18, 0.19, 0.20, 0.07, 0.25, and 0.17. Mean instability going from left to right is 0.26, 0.26, 0.17, 0.28, 0.24, 0.21, and 0.35. This sample was cropped so that all methods could be run.

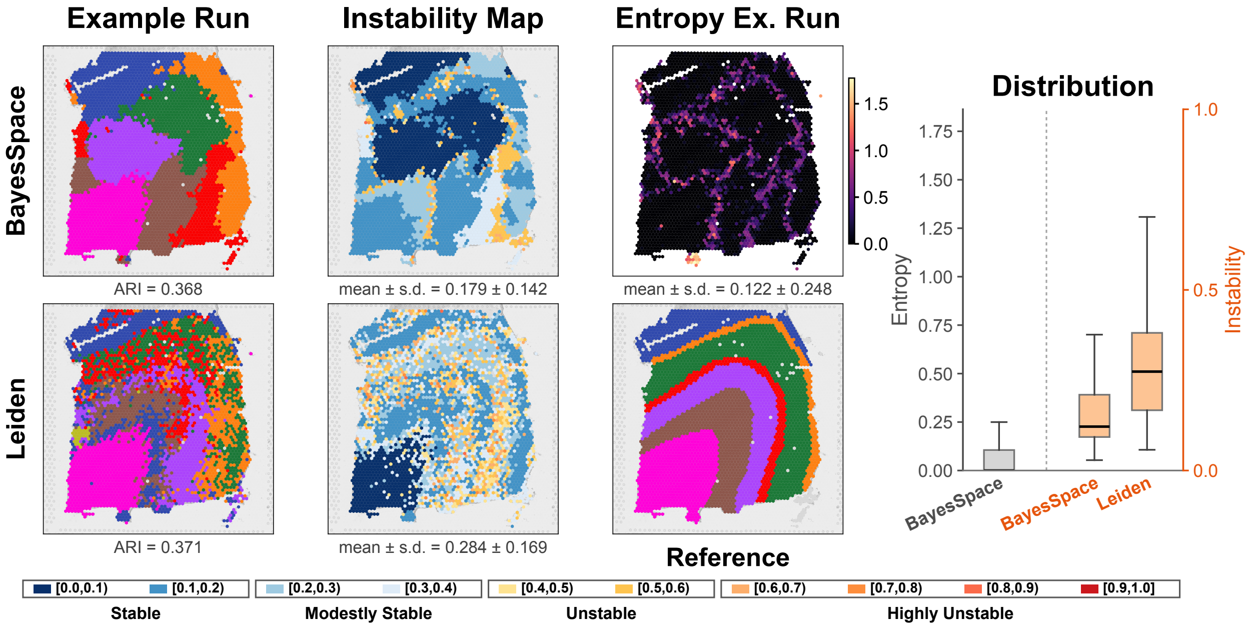

Figure S18: **Instability versus entropy for DLPFC 151674 sample.** **A)** Reference annotation distributed with the benchmark. **B)** Example clustering run for BayesSpace (ARI: 0.37), followed by the per-spot instability score, computed from 50 runs, followed by the per-spot entropy score derived from its cluster membership. Entropy is lower in the middle of clusters and higher at the boundaries, unlike instability. **C)** Example clustering run for Leiden (ARI: 0.37), followed by the per-spot instability score, computed from 50 runs. **D)** Box plot of BayesSpace entropy (gray), BayesSpace per-spot instability (orange), and Leiden per-spot instability (orange); outliers are not shown. Median entropy is zero.

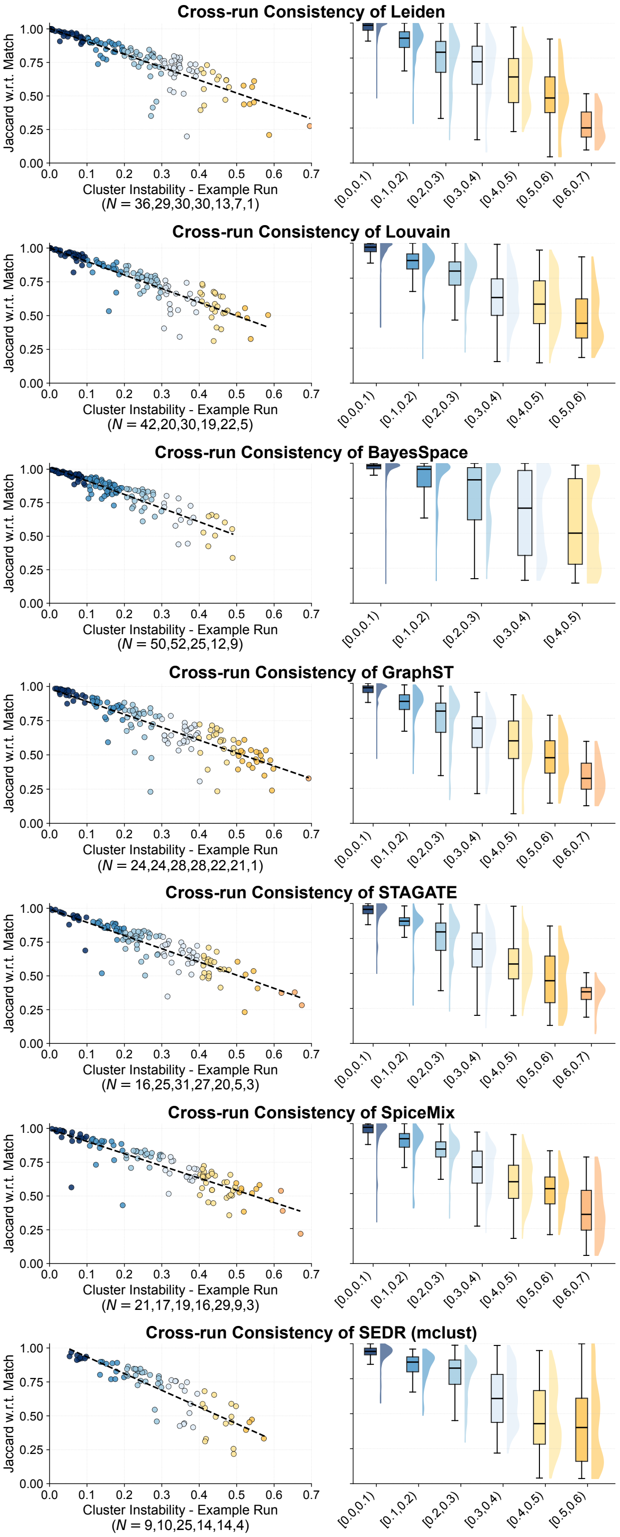

Figure S19: **Cluster cross-run consistency vs. instability.** The SEDR result is shown in Fig. 4B and is therefore not repeated here.

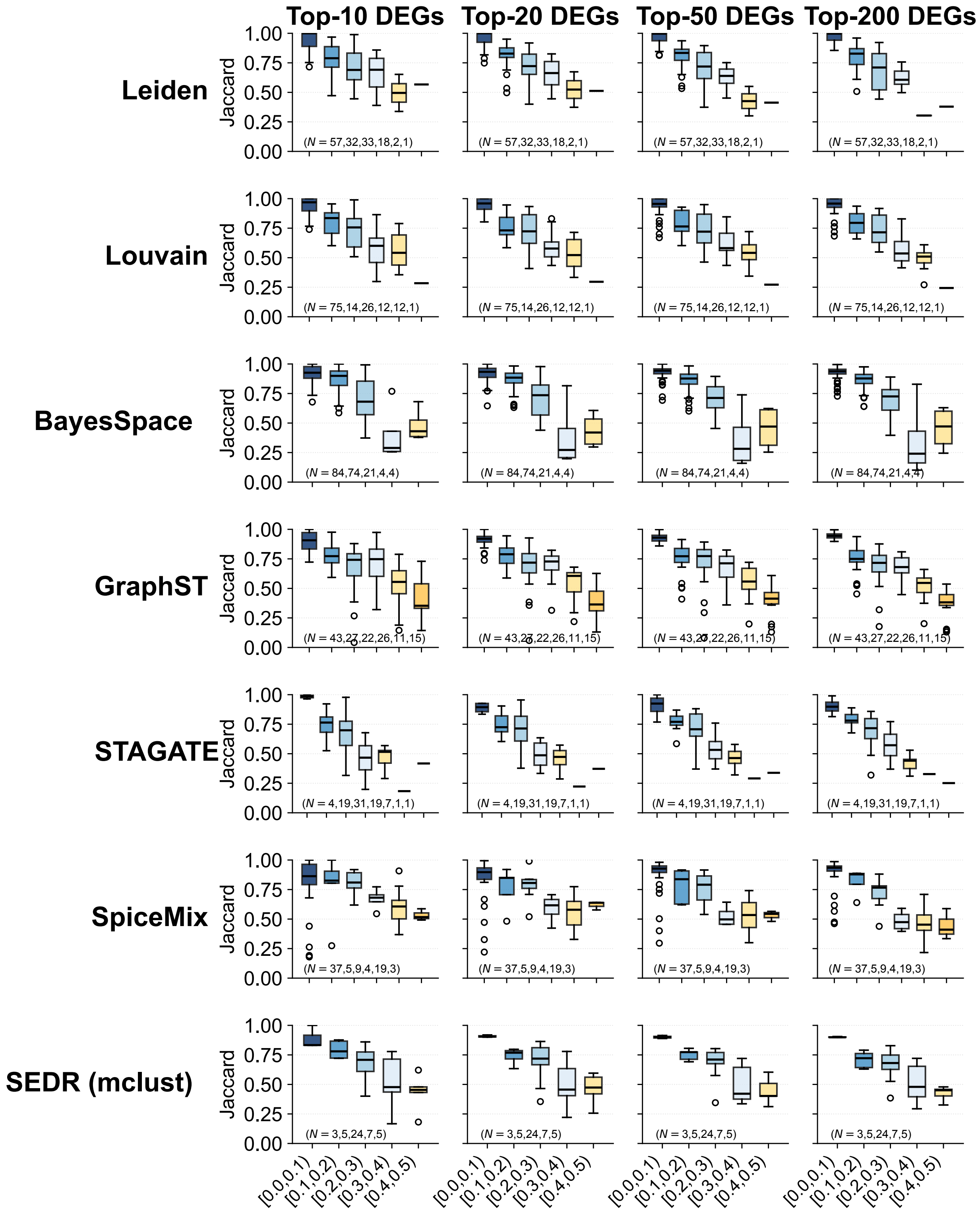

Figure S20: **DEG cross-run consistency (Jaccard) vs. instability.** The SEDR result is shown in Fig. 4C (top) and is therefore not repeated here.

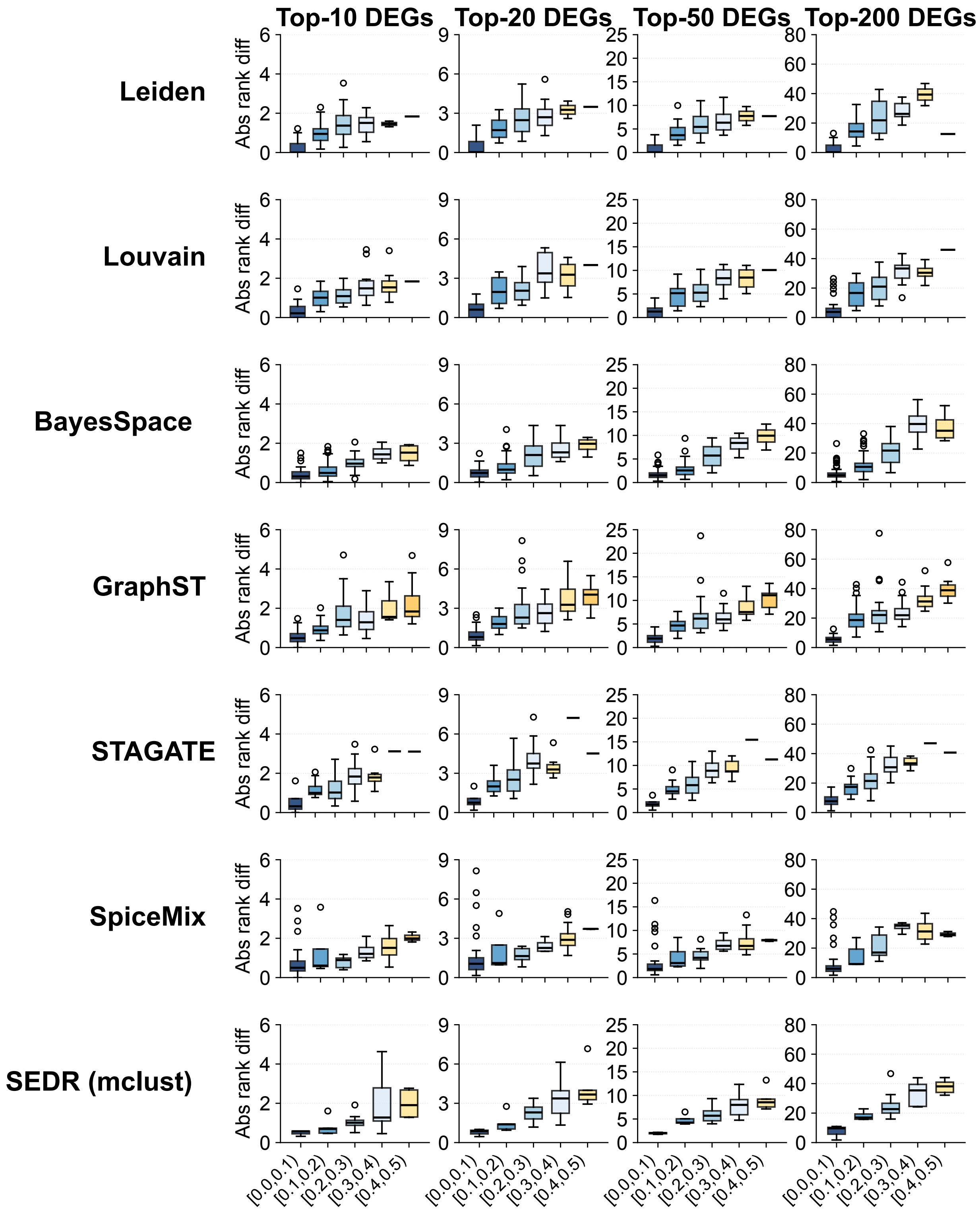

Figure S21: **DEG rank cross-run consistency vs. instability.** Absolute rank difference is for shared DEGs only. The SEDR result is shown in Fig. 4C (bottom) and is therefore not repeated here.

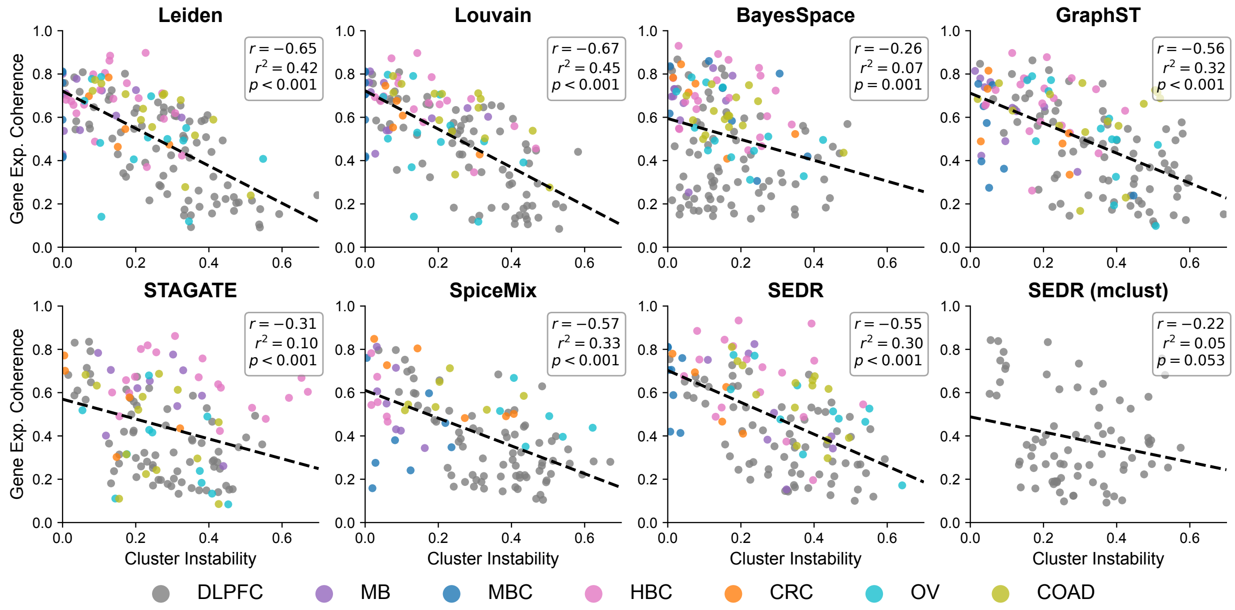

Figure S22: **Gene expression coherence vs. cluster instability.** Each dot represents a cluster produced by the method (subplot) on the indicated sample (dot color). Cluster gene expression coherence is defined as the mean pairwise cosine similarity among spots in this 20-dimensional PCA space (note that this definition is best aligned with the input to Leiden and Louvain and may not be suitable for all methods tested). Cluster instability is the mean score of its spots.

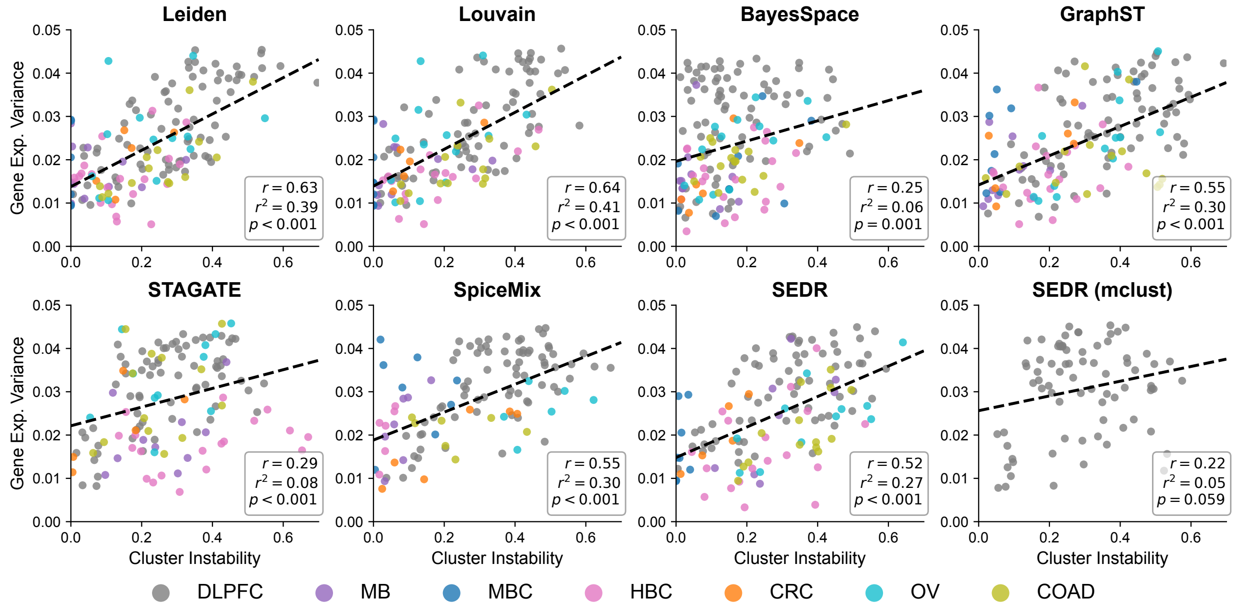

Figure S23: **Gene expression variance vs. cluster instability.** Each dot represents a cluster produced by the method (subplot) on the indicated sample (dot color). Cluster gene expression variance is defined as the mean within-cluster variance across the same 20 principal-component dimensions (note that this definition is best aligned with the input to Leiden and Louvain and may not be suitable for all methods tested). Cluster instability is the mean score of its spots.

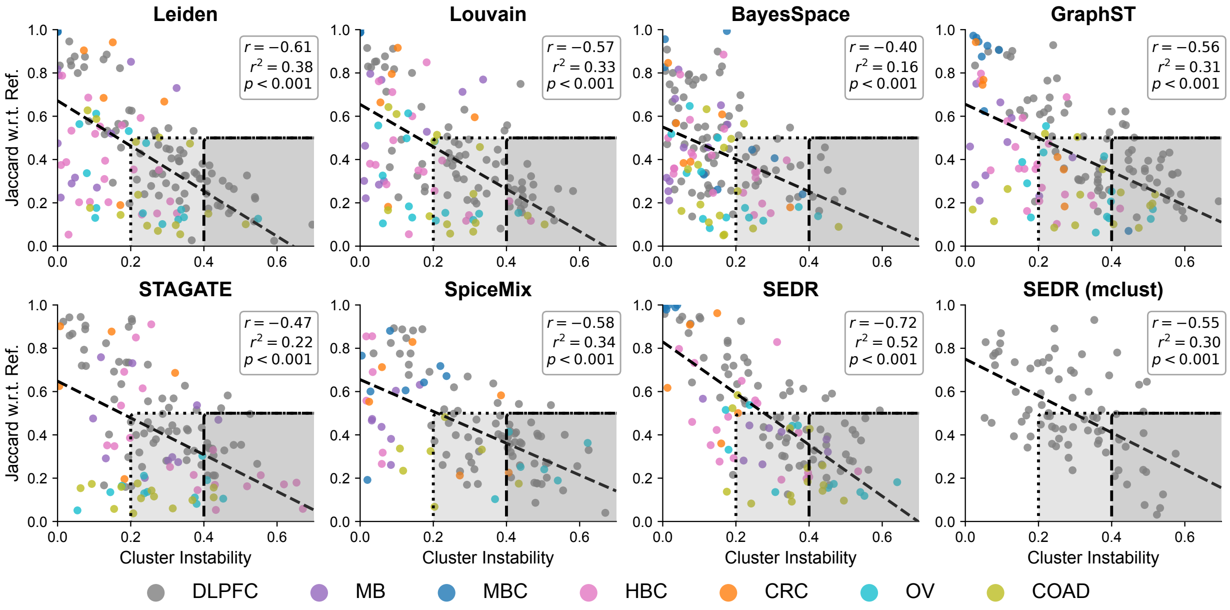

Figure S24: **Cluster similarity w.r.t. reference (Jaccard) vs. instability.** Each dot represents a cluster in the example run produced by the method (subplot) on the indicated sample (dot color). Cluster Jaccard is the number of spots shared by both the predicted cluster and matched reference cluster divided by the number of spots in their union. The best-matching cluster in the reference annotation is one with maximal Jaccard. Cluster instability is the mean score of its spots. Dashed line is linear best fit. The dark gray box highlights unstable clusters (score > 0.4) with low similarity (Jaccard < 0.5). Light grey box is extension to modestly stable clusters.

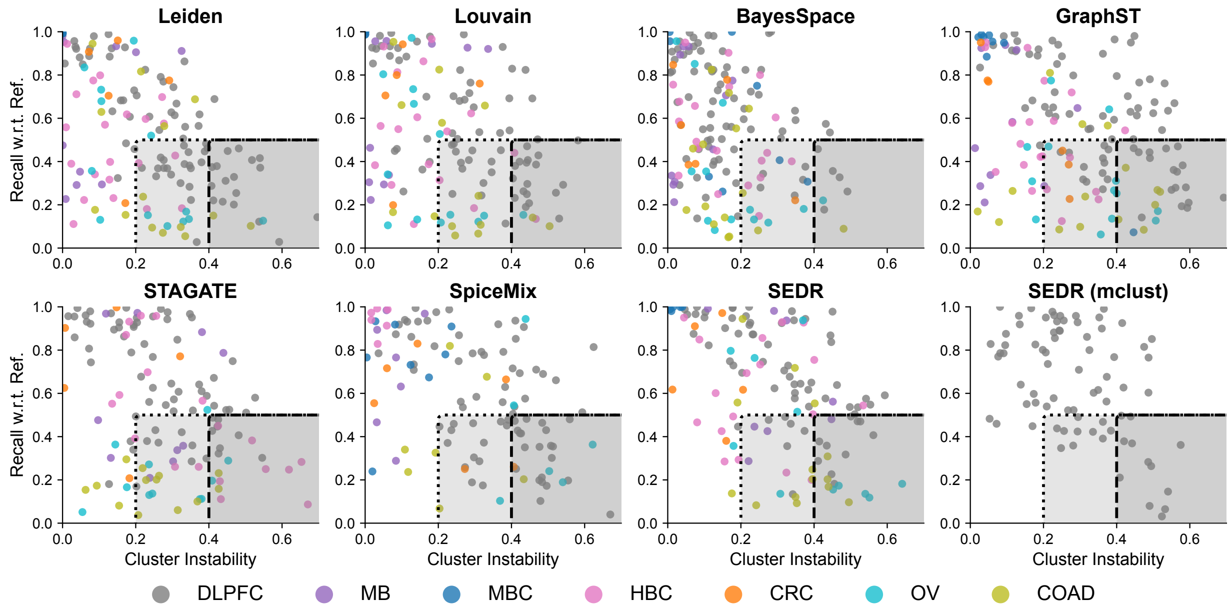

Figure S25: **Cluster recall w.r.t. reference vs. instability.** Each dot represents a cluster in the example run produced by the method (subplot) on the indicated sample (dot color). Cluster recall is the number of spots shared by both the predicted cluster and matched reference cluster divided by the number of spots in the reference cluster. The best-matching cluster in the reference annotation is one with maximal Jaccard. Cluster instability is the mean score of its spots. Dark grey box highlights unstable clusters with low recall ($< 0.5$). Light grey box is extension to modestly stable clusters.

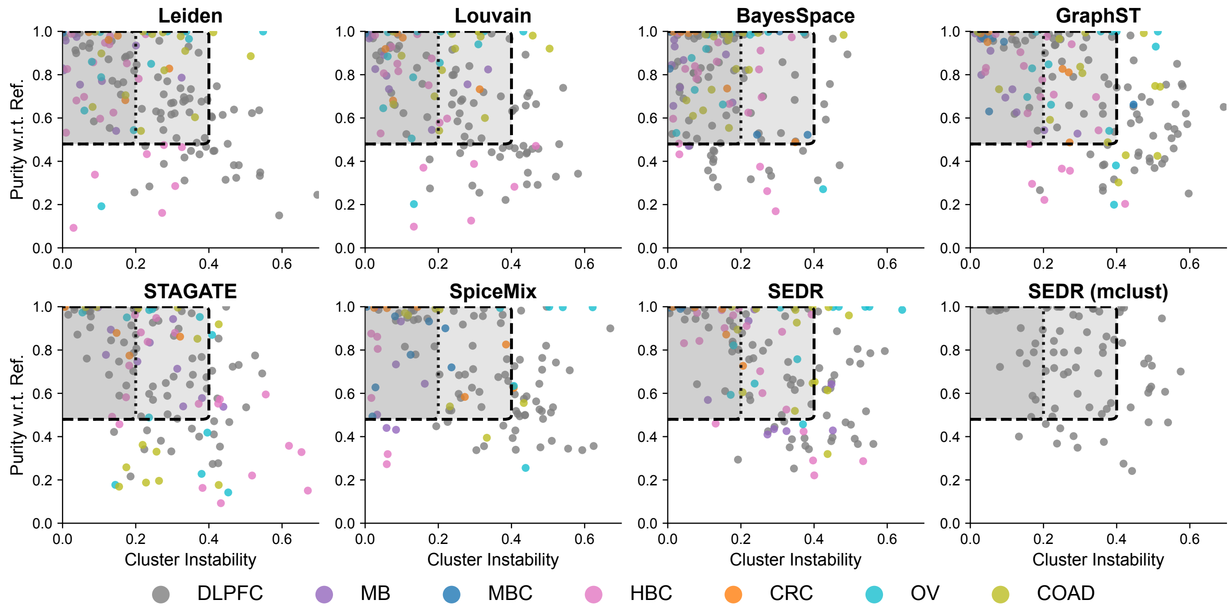

Figure S26: **Cluster purity (precision w.r.t. reference) versus instability.** Each dot represents a cluster in the example run produced by the method (subplot) on the indicated sample (dot color). Cluster purity (i.e., precision) is the number of spots shared by both the predicted cluster and matched reference cluster divided by the number of spots in the predicted cluster. The best-matching cluster in the reference annotation is one with maximal Jaccard. Cluster instability is the mean score of its spots. Dark grey box highlights stable clusters with high purity ($> 0.5$). Light grey box is extension to modestly stable clusters.

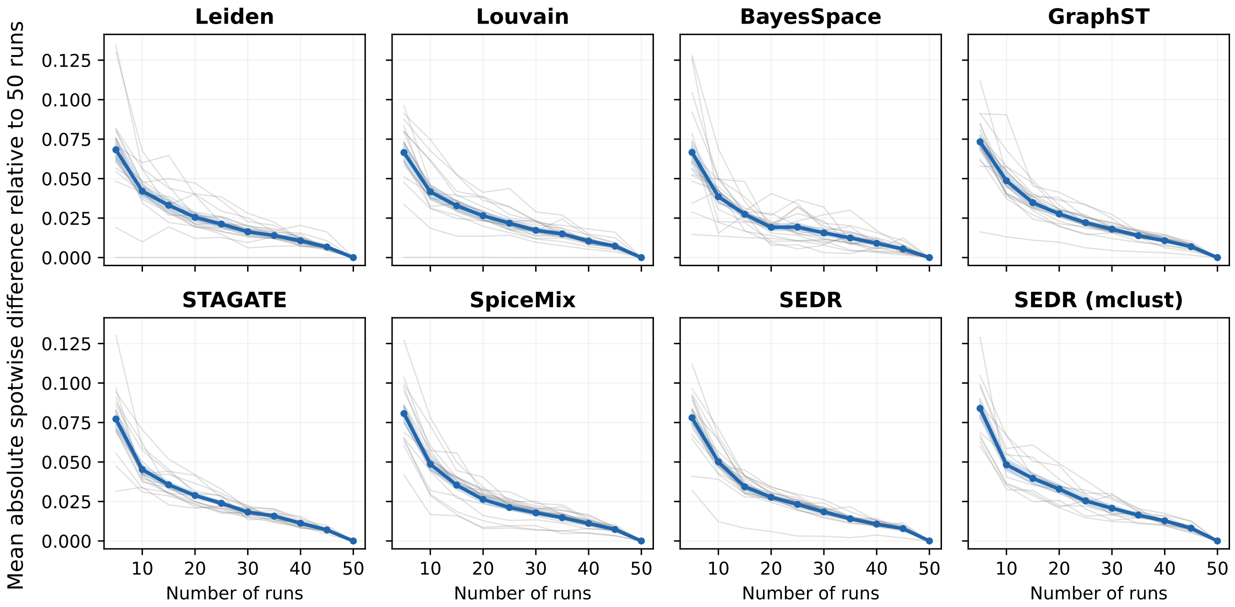

(a) **Instability score convergence using the first $K$ runs.** For each $K$, the instability score was computed using the first $K$ of 50 clustering runs, and the mean absolute spotwise difference from the 50-run estimate was calculated for each sample.

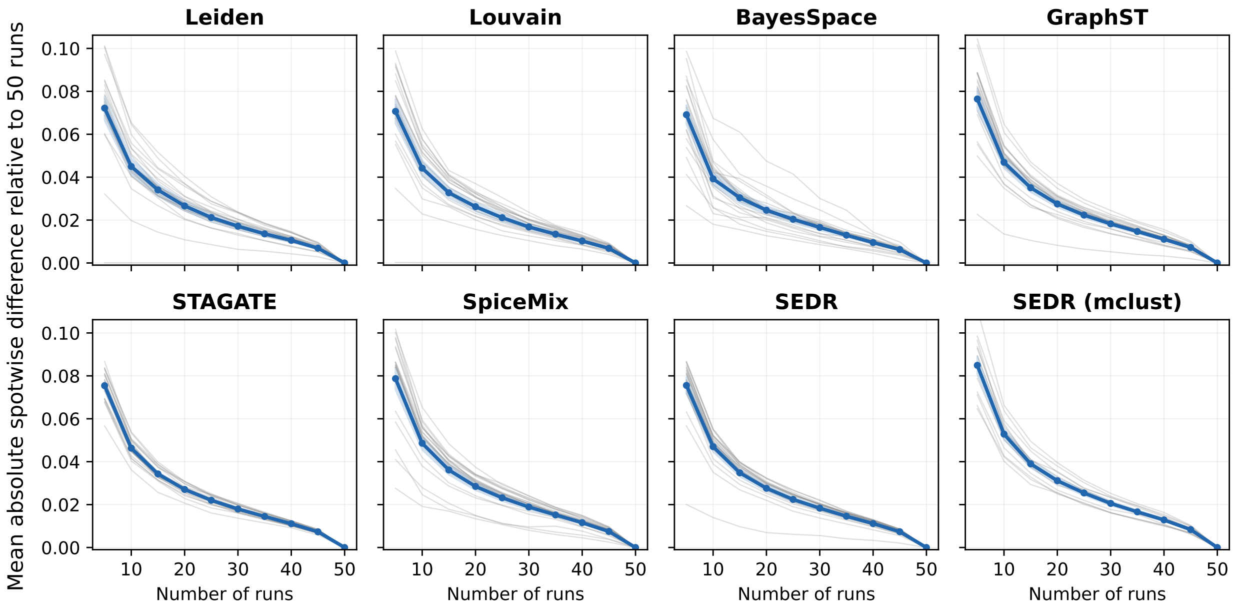

(b) **Instability score convergence with random run sampling.** For each $K$, instability was computed separately for 25 random subsets of $K$ runs. The mean absolute spotwise difference from the 50-run estimate was calculated for each subset and then averaged across the 25 subsets within each sample.

Figure S27: **Instability score convergence with increasing numbers of clustering runs.** Each subplot corresponds to a clustering method, and each gray line represents a sample. The $x$-axis shows the number of runs for computing instability. The $y$-axis shows the mean absolute spotwise difference from the 50-run estimate. Blue lines show the mean across samples, with shading indicating the standard error of the mean. At 25 runs, the mean difference across methods was $\approx 0.02$ in both the first-$K$ and random-subset analyses, indicating that the convergence pattern was not specific to the ordering of clustering runs.

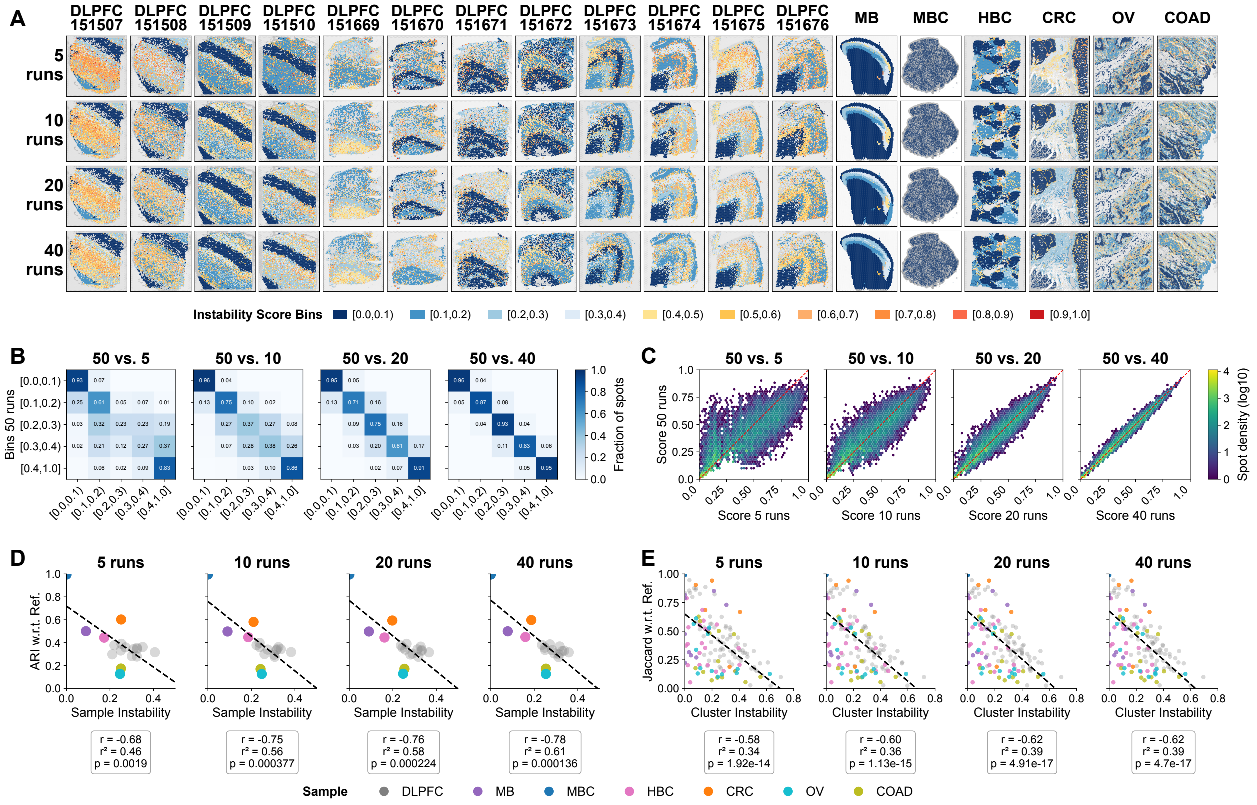

Figure S28: **Impact of varying number of runs on Leiden instability maps.** Note that going from 5 runs to 10 runs means that 5 additional runs are added to the first 5, and similarly for the other increments.

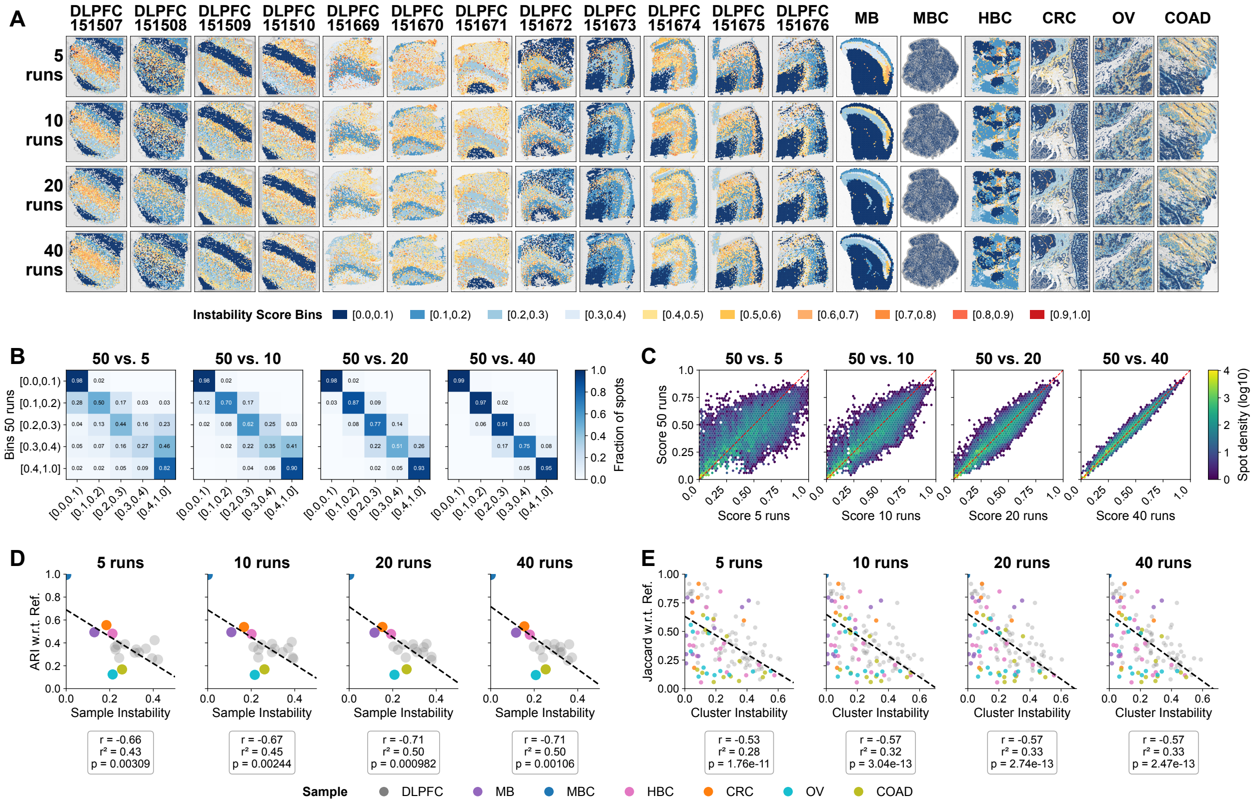

Figure S29: **Impact of varying number of runs on Louvain instability maps.** Note that going from 5 runs to 10 runs means that 5 additional runs are added to the first 5, and similarly for the other increments.

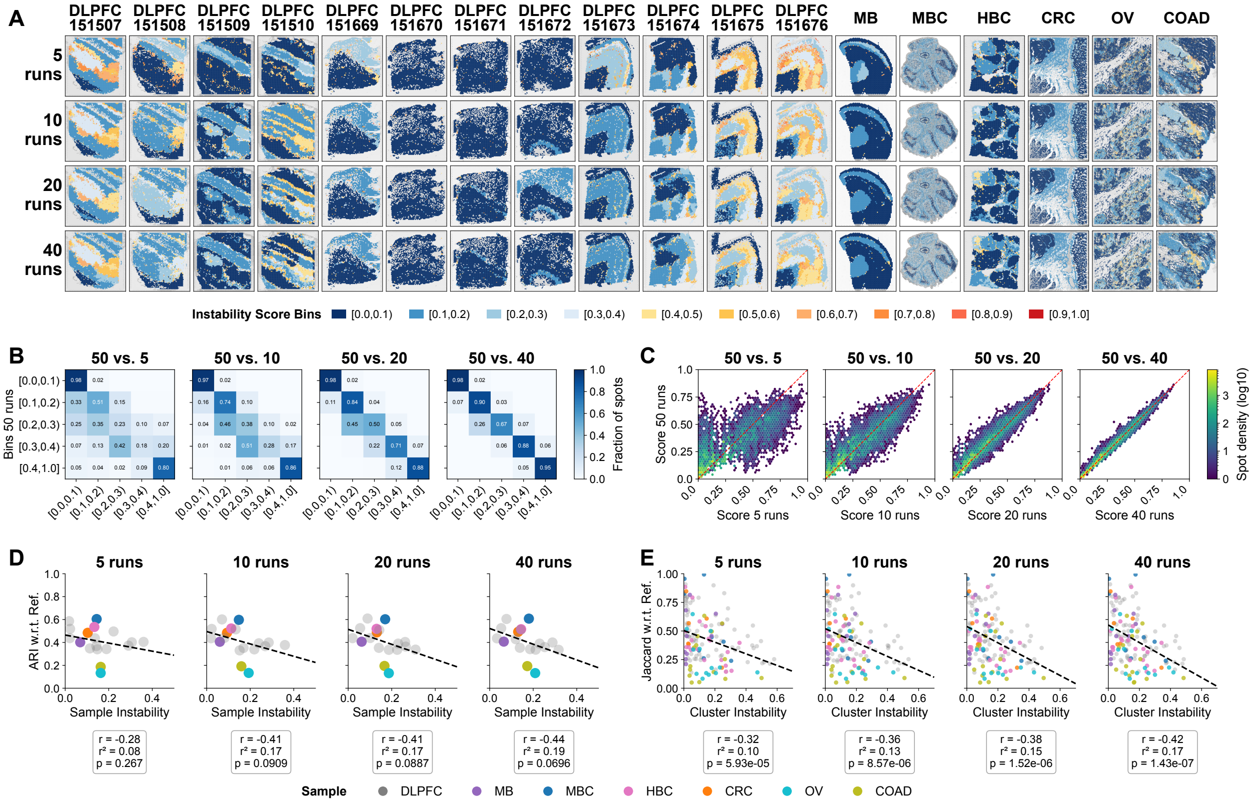

Figure S30: **Impact of varying number of runs on BayesSpace instability maps.** Note that going from 5 runs to 10 runs means that 5 additional runs are added to the first 5, and similarly for the other increments.

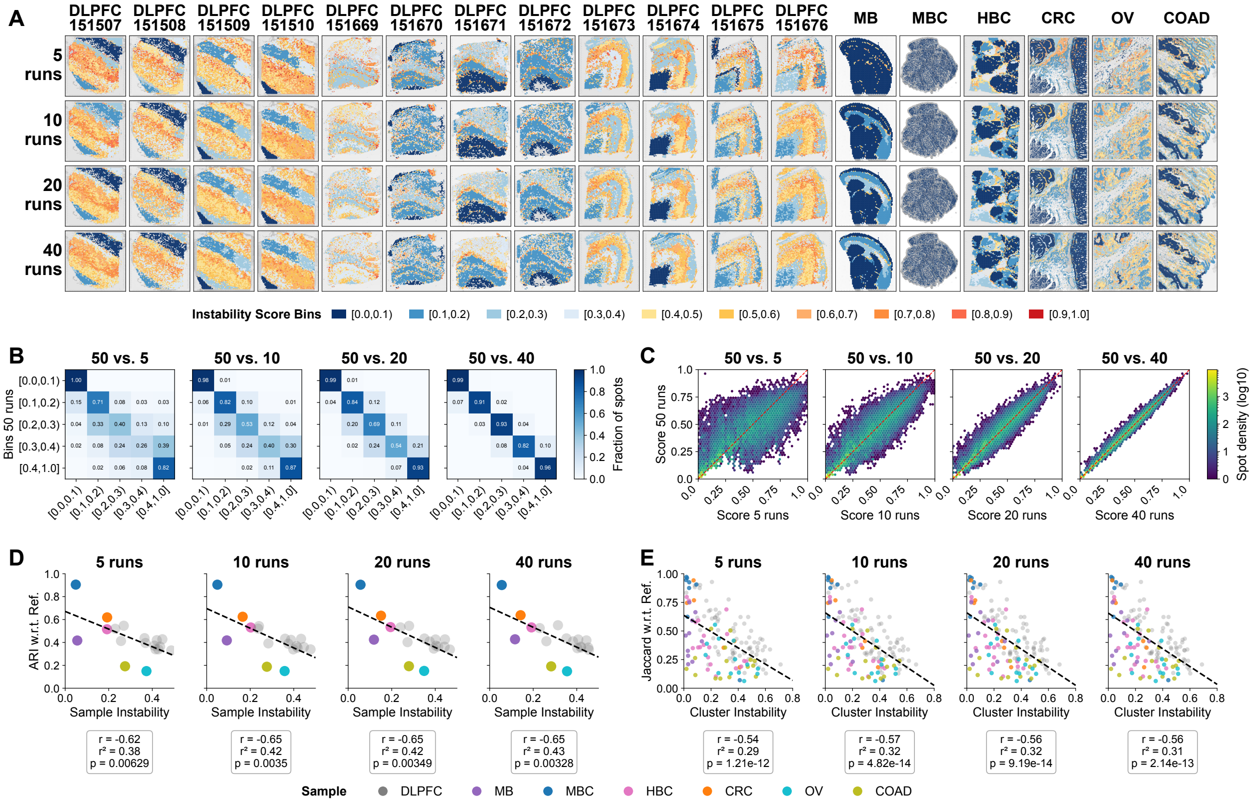

Figure S31: **Impact of varying number of runs on GraphST instability maps.** Note that going from 5 runs to 10 runs means that 5 additional runs are added to the first 5, and similarly for the other increments.

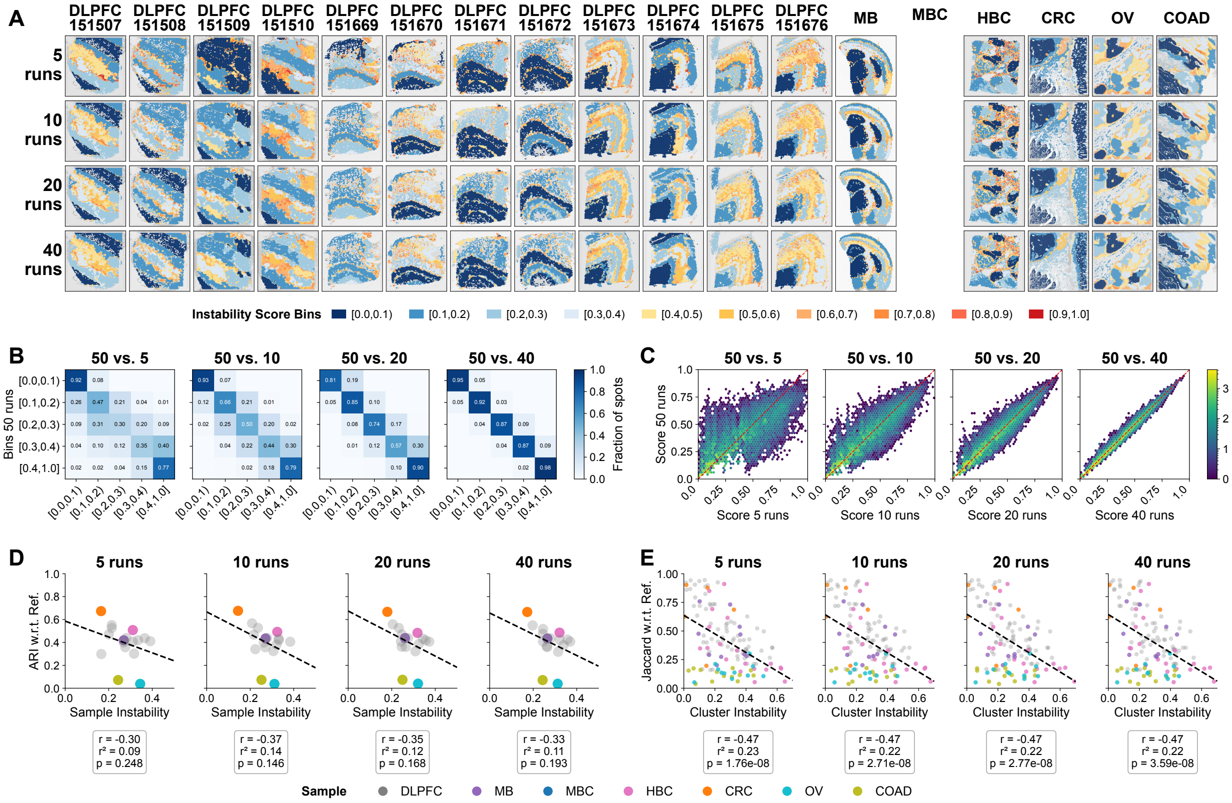

Figure S32: **Impact of varying number of runs on STAGATE instability maps.** Note that going from 5 runs to 10 runs means that 5 additional runs are added to the first 5, and similarly for the other increments. Missing panels are due to STAGATE encountering a memory error on the MBC sample.

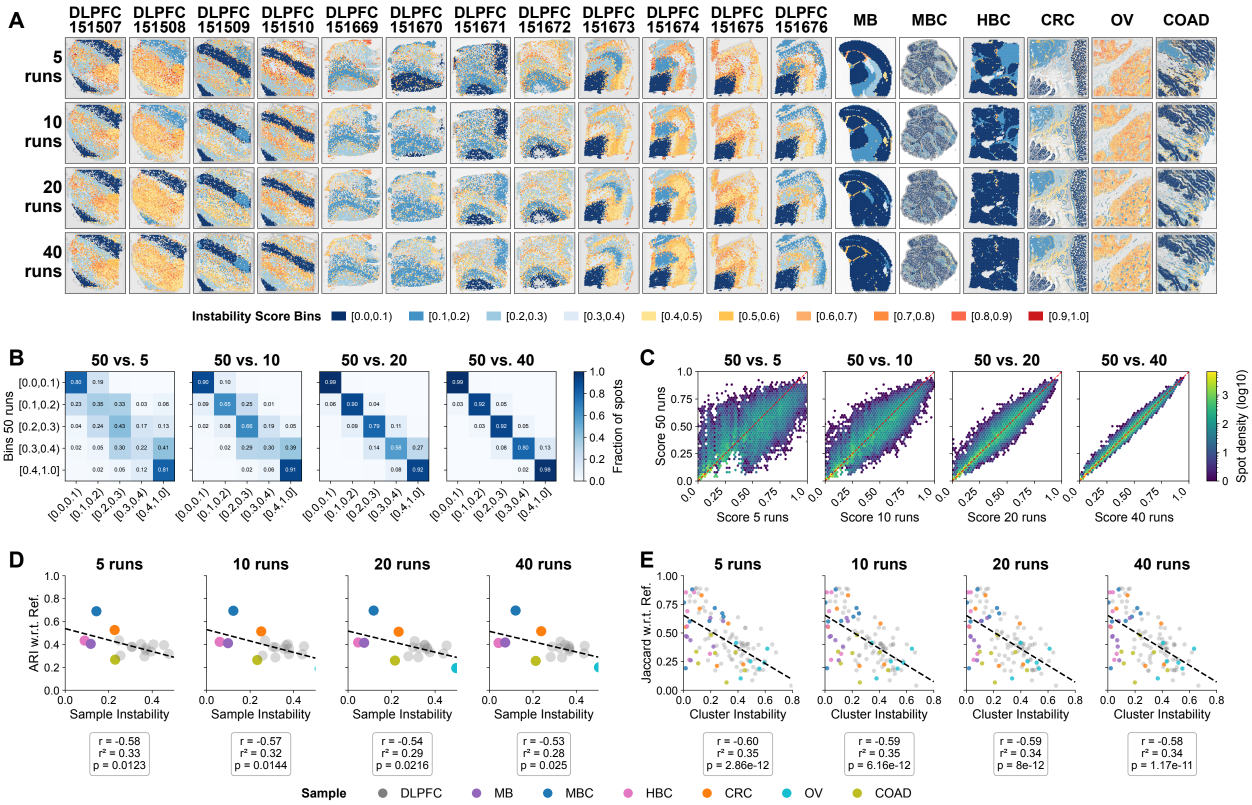

Figure S33: **Impact of varying number of runs on SpiceMix instability maps.** Note that going from 5 runs to 10 runs means that 5 additional runs are added to the first 5, and similarly for the other increments.

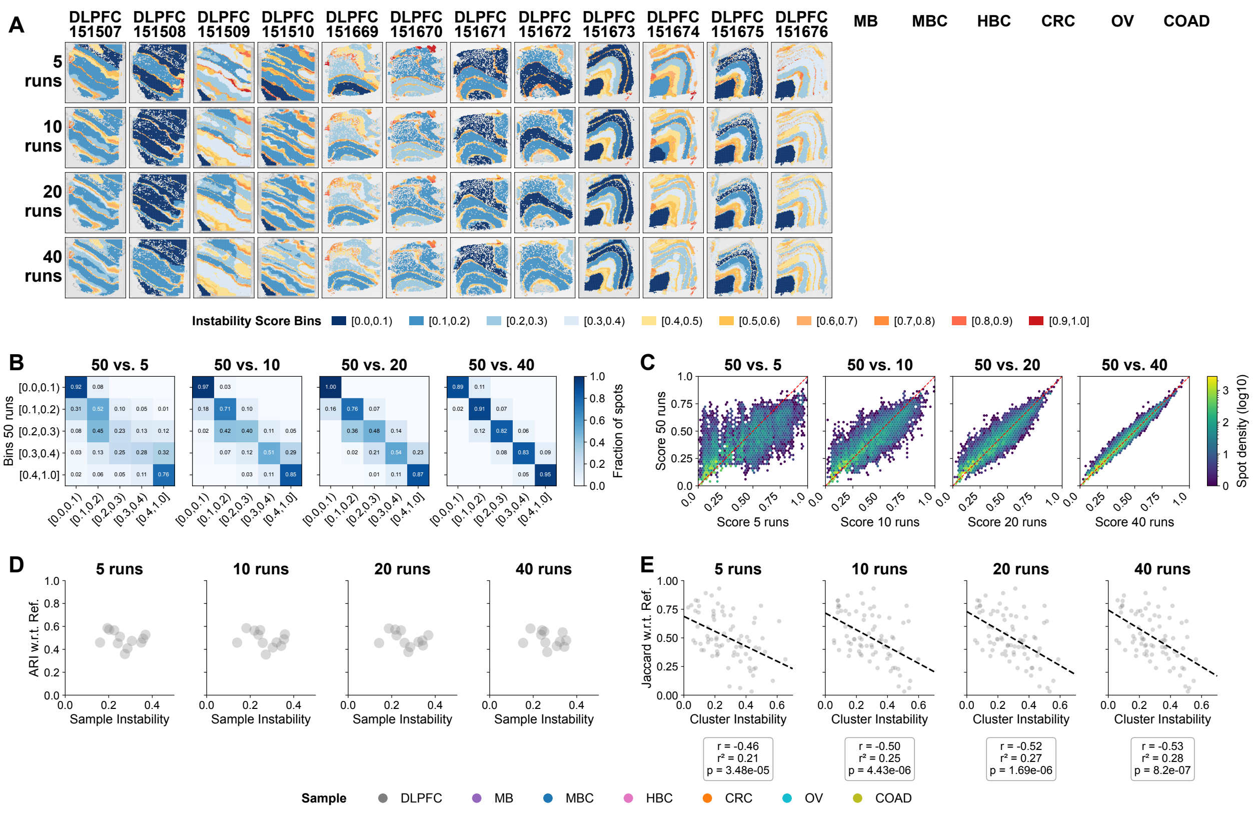

Figure S34: **Impact of varying number of runs on SEDR-mclust instability maps.** Note that going from 5 runs to 10 runs means that 5 additional runs are added to the first 5, and similarly for the other increments. Missing panels are due to SEDR with mclust backend being run on the DLPFC samples only.

### References

[1] 10x Genomics. Human breast cancer (block a section 1), June 2020.

[2] 10x Genomics. Adult mouse brain (FFPE). https://www.10xgenomics.com/datasets/adult-mouse-brain-ffpe-1-standard-1-3-0, 2021. Dataset published 16 August 2021.

[3] 10x Genomics. Visium HD spatial gene expression library, human colorectal cancer (FFPE), March 2024. Dataset published 25 March 2024.

[4] Andrew Cheng, Guanyu Hu, and Wei Vivian Li. Benchmarking cell-type clustering methods for spatially resolved transcriptomics data. *Briefings in Bioinformatics*, 24(1):bbac475, 2023.

[5] Paul Kiessling, Amin El-Heliebi, and Naveed Ishaque. Visium HD human colorectal cancer (FFPE) data release pathologist annotation, May 2024.

[6] Kristen R Maynard, Leonardo Collado-Torres, Lukas M Weber, Cedric Uytingco, Brianna K Barry, Stephen R Williams, Joseph L Catallini, Matthew N Tran, Zachary Besich, Madhavi Tippani, Jennifer Chew, Yifeng Yin, Joel E Kleinman, Thomas M Hyde, Nikhil Rao, Stephanie C Hicks, Keri Martinowich, and Andrew E Jaffe. Transcriptome-scale spatial gene expression in the human dorsolateral prefrontal cortex. *Nature Neuroscience*, 24(3):425–436, 2021.

[7] Pengfei Ren, Rui Zhang, Yunfeng Wang, Peng Zhang, Ce Luo, Suyan Wang, Xiaohong Li, Zongxu Zhang, Yanping Zhao, Yufeng He, Haorui Zhang, Yufeng Li, Zhidong Gao, Xiuping Zhang, Yahui Zhao, Zhihua Liu, Yuanguang Meng, Zhe Zhang, and Zexian Zeng. Systematic benchmarking of high-throughput subcellular spatial transcriptomics platforms across human tumors. *Nature Communications*, 16(1):9232, 2025.

[8] Samuel G Rodriques, Robert R Stickels, Aleksandrina Goeva, Carly A Martin, Evan Murray, Charles R Vanderburg, Joshua Welch, Linlin M Chen, Fei Chen, and Evan Z Macosko. Slide-seq: A scalable technology for measuring genome-wide expression at high spatial resolution. *Science*, 363(6434):1463–1467, 2019.

[9] Han Shu. Hugging Face spatial transcriptomics datasets, 2024.

[10] Hang Xu, Huazhu Fu, Yahui Long, Kok Siong Ang, Raman Sethi, Kelvin Chong, Mengwei Li, Rom Uddamvathanak, Hong Kai Lee, Jingjing Ling, Ao Chen, Ling Shao, Longqi Liu, and Jinmiao Chen. Unsupervised spatially embedded deep representation of spatial transcriptomics. *Genome Medicine*, 16(1):12, 2024.

[11] Fangyuan Zhao. 10x Visium datasets, April 2023.
